# Pulsed priming with the FAK inhibitor narmafotinib enhances both gemcitabine/Abraxane and FOLFIRINOX chemotherapy response in pancreatic cancer

**DOI:** 10.64898/2026.07.23.740234

**Authors:** Kendelle J Murphy, Cecilia R Chambers, Daniel A Reed, Lily M Channon, Niamh EP Mills, Sophie E McKay, Victoria Lee, Anna E Howell, Alice MH Tran, Max Nobis, Astrid Magenau, Janett Stoehr, Brooke A Pereira, Nadia Kuepper, Shona Ritchie, Katie Gordon, Michael Trpceski, Victoria M Tyma, Shanna Hafiz, Vrinda Johri, Alexis Ang, Deborah S Barkauskas, Claire Vennin, Xiao Q Wang, Marjan M Naeini, Braydon Meyer, Amelia L Parker, Sriram Gummadi, Sai V Chitti, Diego Chacon Fajardo, Anaiis Zaratzian, Michael Tayao, Andrew Da Silva, Australian Pancreatic Genome Initiative (APGI), Australian Pancreatic Matrix Atlas (APMA), Anthony J Cesare, Suresh Mathivanan, Clare Stirzaker, Sharissa L Latham, David R Croucher, George Sharbeen, Phoebe A Phillips, Leonard D Goldstein, Ruth J Lyons, Adnan Nagrial, Nick Pavlakis, Anthony J Gill, Andrew V Biankin, Jaswinder Samra, C Elizabeth Caldon, Thomas R Jeffry Evans, Lorraine Chantrill, Lisa G Horvath, Owen J Sansom, Jennifer P Morton, Tri G Phan, Yingxiao Wang, Terrie-Anne Cock, Sarah A Kinkel, Anthony Bishop, Mark Devlin, John Lambert, Thomas R Cox, Marina Pajic, Christopher J Burns, David Herrmann, Paul Timpson

## Abstract

**Background:** Pancreatic ductal adenocarcinoma (PDAC) is a particularly lethal malignancy with few treatment options available. Extensive remodelling of extracellular matrix (ECM) generates a highly fibrotic tumour landscape, which impairs therapeutic response.

**Objective:** We investigated whether stromal priming *via* the highly specific Focal Adhesion Kinase (FAK) inhibitor narmafotinib (AMP945) in combination with the two major standard-of-care chemotherapies in PDAC, gemcitabine/Abraxane and FOLFIRINOX, reduces fibrosis and enhances treatment efficacy.

**Design:** 3D organotypic matrices, intravital imaging, and *in vivo* subcutaneous and orthotopic PDAC models were used to provide a rationale for a first-line priming regimen of narmafotinib prior to chemotherapy.

**Results:** Neoadjuvant chemotherapy induces fibrosis in PDAC indicating a need for upfront first-line priming of the ECM to normalise the stroma for optimal treatment response. Narmafotinib is a new potent small molecule FAK inhibitor. Phase I safety data shows excellent safety, tolerability, and pharmacokinetics following oral administration in humans. We reveal that narmafotinib treatment during early ECM remodelling (‘priming’) reduces fibrosis, while limiting subsequent PDAC invasion. Moreover, intravital imaging demonstrates real-time FAK inactivation and cell cycle stalling, leading to improved chemotherapeutic efficacy upon narmafotinib priming *in vivo*. Long-term assessment in patient-derived models shows that narmafotinib priming prior to gemcitabine/Abraxane or FOLFIRINOX reduces PDAC progression and extends survival in both chemotherapy settings.

**Conclusions:** Our results using these Phase II-ready drug combinations strongly support the clinical assessment of narmafotinib in PDAC. Narmafotinib is currently in Phase Ib/IIa trials, assessing a pulsed dosing regimen prior to gemcitabine/Abraxane, and warrants further clinical assessment in combination with FOLFIRINOX.

**SIGNIFICANCE OF THIS STUDY:** *What is already known on this topic:*

- Pancreatic cancer (PC) is one of the most lethal malignancies and is characterised by a dense, fibrotic stroma, which impairs chemotherapy efficacy.
- The non-receptor tyrosine kinase FAK is known to promote cancer fibrosis and therefore represents a therapeutic target to normalise the PC stroma and to improve chemotherapy performance.

*What this study adds:*

- Neoadjuvant chemotherapy induces early fibrosis indicating a need for upfront first-line priming of the ECM to blunt or normalise stromal fibrosis for optimal response to therapy.
- The small molecule inhibitor narmafotinib (which is currently under Phase Ib/IIa clinical trial assessment) shows high specificity towards FAK as well as desirable pharmacokinetics and pharmacodynamics in healthy human volunteers.
- Early short-term narmafotinib priming reduces fibrosis and improves the efficacy of subsequent standard-of-care gemcitabine/Abraxane chemotherapy.
- FOLFIRINOX (oxaliplatin, irinotecan, leucovorin and 5-fluorouracil) is a multi-agent chemotherapy preferentially used in PDAC patients with good performance status. Our results demonstrate that narmafotinib priming also improves FOLFIRINOX efficacy, leading to extended survival in patient-derived PDAC models.

*How this study might affect research, practice, or policy:*

- This study supports the clinical development of narmafotinib in combination with both gemcitabine/Abraxane (ACCENT trial) and further FOLFIRINOX standard-of-care chemotherapies for PDAC patient treatment.
- The first-line priming strategy and early ECM normalisation used in this study may also be applicable to other combination therapy settings and warrants further investigation in ongoing clinical studies.

## INTRODUCTION

Pancreatic ductal adenocarcinoma (PDAC) is a highly fibrotic and aggressive malignancy with a 5-year survival rate of 13% (*1*). Due to a lack of early symptoms, most patients present with inoperable locally advanced or metastatic disease (*1, 2*). Clinically, standard-of-care is predominantly one of two systemic chemotherapy regimens: gemcitabine and Abraxane (nab-paclitaxel) (*2, 3*), or FOLFIRINOX (oxaliplatin, irinotecan, leucovorin and 5-fluorouracil), which both result in modest survival benefits (*4–6*). FOLFIRINOX is generally only provided to patients with good Eastern Cooperative Oncology Group (ECOG) performance status due to tolerability issues, and combination therapies with FOLFIRINOX have not been widely explored (*4*). As such, there is an urgent need to enhance treatment efficacy in both chemotherapy settings to improve PDAC patient outcomes.

PDAC progression is accompanied by a fibrotic response, involving remodelling of extracellular matrix (ECM), which can promote tissue stiffness, disease progression and chemoresistance (*7–9*). Importantly, while complete ablation of stromal elements has conflicting effects on PDAC survival (*8, 10, 11*), recent work suggests that subtle, fine-tuned manipulation of the ECM architecture can improve drug response and outcomes (*7, 8, 12–16*).

Focal Adhesion Kinase (FAK) is a non-receptor tyrosine kinase that modulates interactions between cancer cells and the ECM (*17, 18*). FAK expression and activity are frequently upregulated in multiple cancers, where FAK was shown to regulate proliferation, survival and invasion as well as ECM remodelling and stiffening (*12, 14, 16–24*). Here, we provide preclinical data showing that the small molecule, narmafotinib, is a highly potent, selective FAK inhibitor, with desirable potency, selectivity, and pharmacokinetics (PK) in healthy human volunteers, which can be utilised to improve chemotherapy response in PDAC. Using 3D organotypic matrices we show anti-fibrotic effects of narmafotinib during ECM remodelling, which restricts PDAC invasion. Moreover, real-time intravital (*in vivo*) imaging using a FAK biosensor and FUCCI cell cycle reporter in the live PDAC microenvironment demonstrates that narmafotinib reduces fibrosis, while potentiating therapeutic efficacy (*25–27*). Lastly, we reveal the preclinical efficacy of narmafotinib in combination with gemcitabine/Abraxane and, for the first time, FOLFIRINOX, to extend survival in long-term patient-derived models.

## MATERIALS AND METHODS

### Study Design

Amplia Therapeutics Phase Ia clinical trial AMP945-101 (ACTRN12620000894998) was a randomised, double-blind, placebo-controlled study of the safety, tolerability, and pharmacokinetics of single and repeat doses of narmafotinib administered orally to healthy adult volunteers.

For *in vivo* experiments, treatment start and regimen as well as study endpoints are outlined in the Figures and Supplementary Material. Animal numbers are defined in the corresponding Figure Legends and in the Supplementary Material. All animal experiments were performed in accordance with the Garvan/St. Vincent’s Animal Ethics Committee guidelines (Animal Research Authorities 19/10, 19/13, 22/08, 22/09, 22/10, 25/10, 25/11, 25/12) and in compliance with the Australian code of practice for care and use of animals for scientific purposes.

Tumour microarrays from the APGI ICGC cohort were used to assess stromal collagen levels *via* Picrosirius Red staining and pTyr-397-FAK *via* immunohistochemistry (IHC) in a cohort of 226 patients. Following quantification of average collagen length by TWOMBLI (*28*) and pTyr-397-FAK levels *via* QuPath (*29*), samples were segregated into high or low fibre length and pTyr-397-FAK, and Kaplan-Meier curves were generated.

Further information on Methods is detailed in the Supplementary Material.

## RESULTS

### Fibrosis is increased in PC following neoadjuvant chemotherapy, while FAK activation and ECM remodelling correlates with outcomes

The ECM provides structural support to solid tumours, with a transition from isotropic (poorly aligned) to anisotropic (highly aligned) structures often associated with aggressive cancers (*7, 8, 12–14, 24, 28, 30–33*). Previous studies have shown that standard-of-care chemotherapy can stimulate ECM remodelling, driving further fibrosis (*31, 33–35*). Therefore, we sought to assess fibrosis in PC tumours isolated from treatment-naïve patients compared to patients who received neoadjuvant gemcitabine/Abraxane or FOLFIRINOX (Figure 1A,B, Table S1). PC samples from the Royal North Shore Hospital/Australian Pancreatic Matrix Atlas (APMA (*12, 33*)) cohort (Table S1) were stained with Picrosirius Red to determine collagen I/III organisation, and revealed an increase in fibrillar collagen in both neoadjuvant gemcitabine/Abraxane– and FOLFIRINOX-treated patients compared to treatment-naïve individuals (Figure 1A). Further analysis using polarised light imaging showed that while overall collagen fibre orientation was not affected, neoadjuvant chemotherapy significantly increased collagen birefringence signal (Figure S1A,B (*36, 37*)). This increased collagen deposition and remodelling was further confirmed by Second Harmonic Generation (SHG) imaging (Figure 1B, Movie S1, (*25, 32, 38*)), which suggests that fibrosis is an early stromal response to chemotherapy in PC and warrants the assessment of anti-fibrotic priming prior to chemotherapy in a first-line setting.

**Figure 1.**
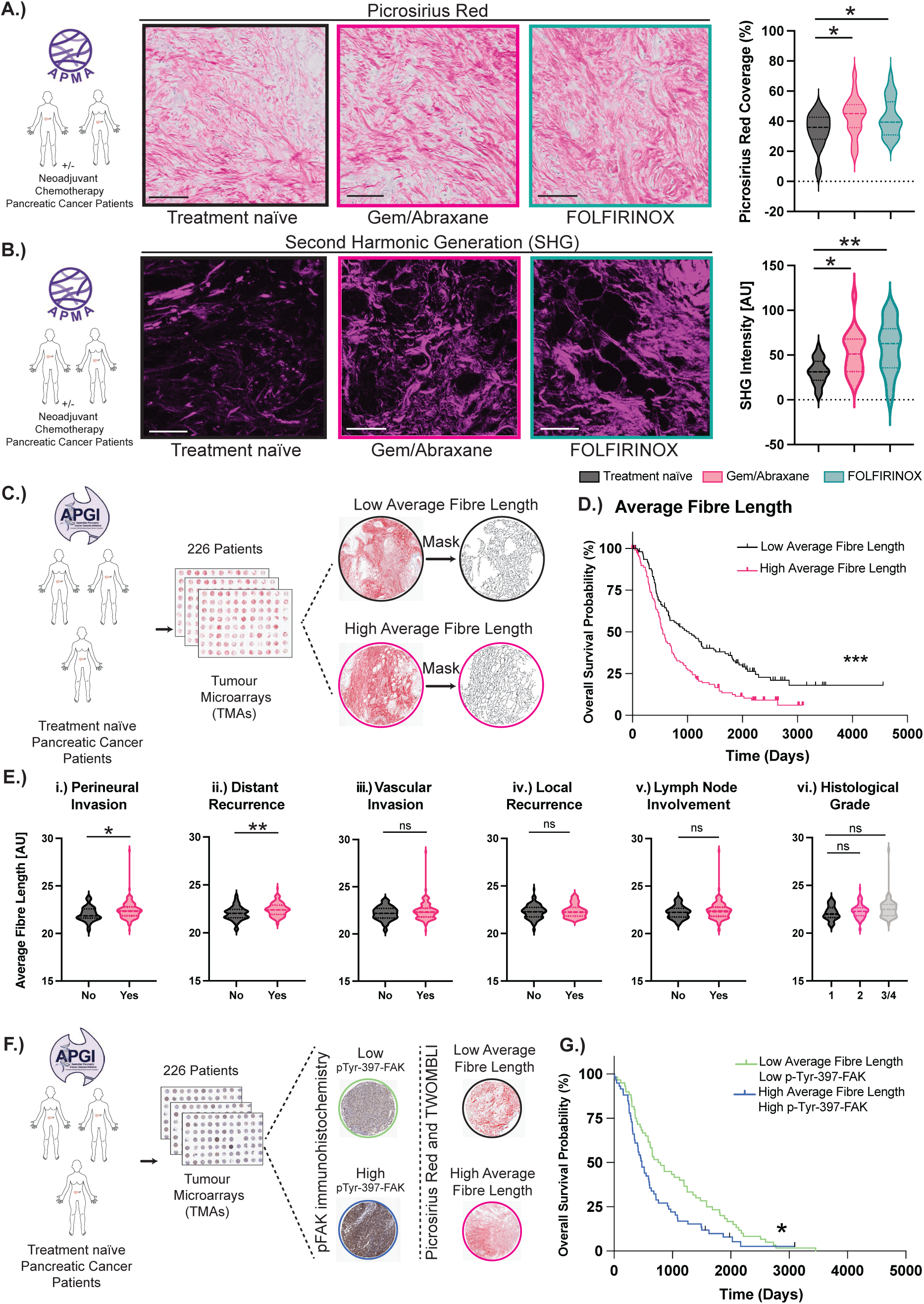
ECM organisation and FAK signature correlate with clinico-pathological features in PC patients. (**A,B**) Representative images and quantification of Picrosirius Red brightfield (**A.**) or SHG (**B.**) from treatment-naïve *versus* neoadjuvant gemcitabine/Abraxane– or FOLFIRINOX-treated PC patients in the Royal North Shore Hospital/APMA cohort. Scale bar, 50 µm (**A.**), 100 µm (**B.**). n=22 treatment-naïve, n=20 neoadjuvant gemcitabine/Abraxane, n=19 neoadjuvant FOLFIRINOX patients, with ≥5 ROIs per tumour. (**C.**) Schematic of treatment-naïve patient TMA staining with Picrosirius Red (three cores per patient). (**D.**) Kaplan-Meier analysis of human PC patient overall survival based on high (pink, n=113) *versus* low average fibre length (black, n=113). (**E.**) Analysis of clinico-pathological differences (**i.)** perineural invasion, **ii.)** distant recurrence, **iii.)** vascular invasion, **iv.)** local recurrence, **v.)** lymph node involvement, **vi.)** histological grade) between patients with high and low average fibre length (n=186 patients). (**F.**) Schematic of treatment-naïve patient TMA with high or low Picrosirius Red or pTyr-397-FAK. (**G.**) Kaplan-Meier analysis of human PC patient overall survival based on fibre length and pFAK levels (high [blue, n=56] *versus* low [green, n=60]). Violin plots are median (dashed line) with quartiles (dotted line), p-values determined using (**A**) Kruskal-Wallis test with Dunn’s multiple comparisons, (**B**) one-way ANOVA with Sídák multiple comparison test, (**E**) unpaired two-tailed t-test with Welch correction for unequal variance or (**D,G**) log-rank Mantel-Cox test. ns, P≥0.05, *P<0.05, **P<0.01, ***P<0.001, ****P<0.0001.

Subtle changes in ECM organisation may affect PC progression and outcomes (*7, 20, 31, 34, 35*). To systematically examine the influence of ECM ultrastructure/organisation on PC prognosis, human tumour microarrays (TMAs) from the Australian Pancreatic Genome Initiative (APGI) and International Cancer Genome Cohort (ICGC) of 226 treatment-naïve patient samples (*39–42*) were stained with Picrosirius Red (Figure 1C) and assessed *via* semi-supervised TWOMBLI analysis to extract ultrastructural features of the ECM and their association with long-term follow-up (Figure S1C,D (*28*)). This revealed that treatment-naïve PC tissues with high collagen fibre length correlated with poor overall patient survival (Figure 1D), as well as enhanced perineural invasion and distant recurrence (Figure 1E), which have previously been linked to unfavourable prognosis (*43, 44*). Assessment of additional collagen features in the APGI cohort of PC patients revealed that while general ECM density metrics, such as fibre area, box-counting fractal dimension and lacunarity, did not correlate with clinico-pathological features assessed (Figure S2A-C), other ultrastructural fibre metrics beyond fibre length, such as the number of fibre end– and branchpoints correlate with distant recurrence (Figure S2D-F). While hyphal growth unit (HGU, number of endpoints per unit length) correlated with both the presence of perineural invasion and distant recurrence (Figure S2G). These findings indicate that higher-order features of collagen network topology, rather than bulk matrix abundance, can capture clinically relevant aspects of stromal remodelling associated with tumour aggressiveness. Together, these results demonstrate that fibrotic collagen features can be linked to patient prognosis, positioning collagen network topology as an informative marker of disease aggressiveness, and support the rationale for co-targeting fibrosis alongside chemotherapy to improve outcomes in PC.

FAK is well-known to regulate ECM organisation and cancer cell response to ECM cues (*12, 14, 17–20*). Parallel to our TWOMBLI analysis of ECM ultrastructure, we stained TMAs from the APGI cohort for FAK activity (pTyr-397-FAK). Association between FAK activity and clinico-pathological features in the APGI patient cohort showed significant correlation between stromal FAK activity and distant recurrence (Table S2) in line with previously published literature (*14, 19, 24*). Interestingly, quantification of FAK activity in combination with TWOMBLI quantification of collagen features (Figure 1F), revealed that high collagen fibre length and high FAK activity correlated with poor overall patient survival (Figure 1G). These results emphasise the potential role FAK plays in PC fibrosis and outcomes, suggesting that FAK inhibition could be utilised to reduce fibrosis and improve response to chemotherapy in PC.

### Narmafotinib is a selective, orally available FAK inhibitor

Narmafotinib (AMP945: Amplia Therapeutics) was discovered through a structure-guided drug discovery program (Figure 2A). This drug-like compound is an ATP-competitive small molecule inhibitor that in cell-free assays possesses excellent potency for FAK (IC_50_ 2.2 nM) and is selective over the closely related kinase Pyk2 (IC_50_ 550 nM, Figure 2B) showing a superior selectivity for FAK (>200-fold) compared to other commonly used FAK inhibitors, such as PF562271 (∼10-fold(*45*)) or defactinib (near equal inhibitor of FAK and Pyk2 (*46*)). In a panel of PDAC cell lines and fibroblasts, 10 nM to 50 nM narmafotinib was sufficient to reduce FAK activity (Figure S3A-E). Narmafotinib displays high selectivity across a panel of 403 kinases (Figure 2C, Table S3) with kinome selectivity superior to that reported for defactinib (*47*). The compound has good pharmacokinetics in mice, where orally dosed narmafotinib possesses ∼7 hours half-life and 45% bioavailability (Table S4). Given the excellent preclinical activity and safety profile of narmafotinib, a Phase I randomised double-blind, placebo-controlled study in 56 healthy volunteers was undertaken (ACTRN12620000894998) to investigate safety, tolerability, pharmacokinetic (PK) and pharmacodynamic (PD) effects of orally dosed drug. In a single ascending dose (SAD) study narmafotinib (15-125 mg) could be detected in the plasma of fasted volunteers for 24 h (Figure 2D). Prior food consumption had no effect on the mean time (2-6 h) to maximum plasma concentration. The median half-life of narmafotinib in plasma was 15.7-23 h (Figure 2D,E). Plasma concentrations from the multiple ascending dose (MAD) study indicated that a steady state of circulating narmafotinib was achieved for all assessed drug dosages after 3-4 days (Figure 2F,G). Lastly, in an exploratory PD analysis using skin punch biopsies from healthy volunteers, baseline pTyr-397-FAK levels were compared to drug target levels after dosing with narmafotinib. Here dose– and exposure-related reductions in pTyr-397-FAK were detected, indicating that drug target engagement had occurred in proportion to both the administered dose and the total area under the curve plasma exposure to narmafotinib (AUC_0-inf_, (Figure 2H,I)). No serious or severe treatment emergent adverse events (TEAEs), nor any TEAEs leading to study withdrawal were observed. In brief, 16 MAD participants reported a total of 24 TEAEs during the study (Figure 2J). TEAE frequency did not appear to be dose-related, with the highest number of TEAEs reported in Cohort 1 at 25 mg narmafotinib (9 events reported by 5 [83.3%] participants), followed by Cohort 2 at 50 mg narmafotinib (6 events reported by 4 [66.7%] participants), pooled placebo (5 events reported by 4 [66.7%] participants) and Cohort 3 at 100 mg narmafotinib (4 events reported by 3 [50.0%] participants) (Figure 2J). There was no relationship observed between treatment-related TEAE frequency and dose, no serious or severe TEAEs, nor any TEAEs leading to dosing discontinuation or study withdrawal. The most frequently reported TEAEs during the study were headache and backpain, with 4 events reported by 4 (16.7%) participants, and 3 events reported by 3 (12.5%) participants, across all dose groups, respectively. Taken together, the Phase I study data supports the clinical development of narmafotinib.

**Figure 2.**
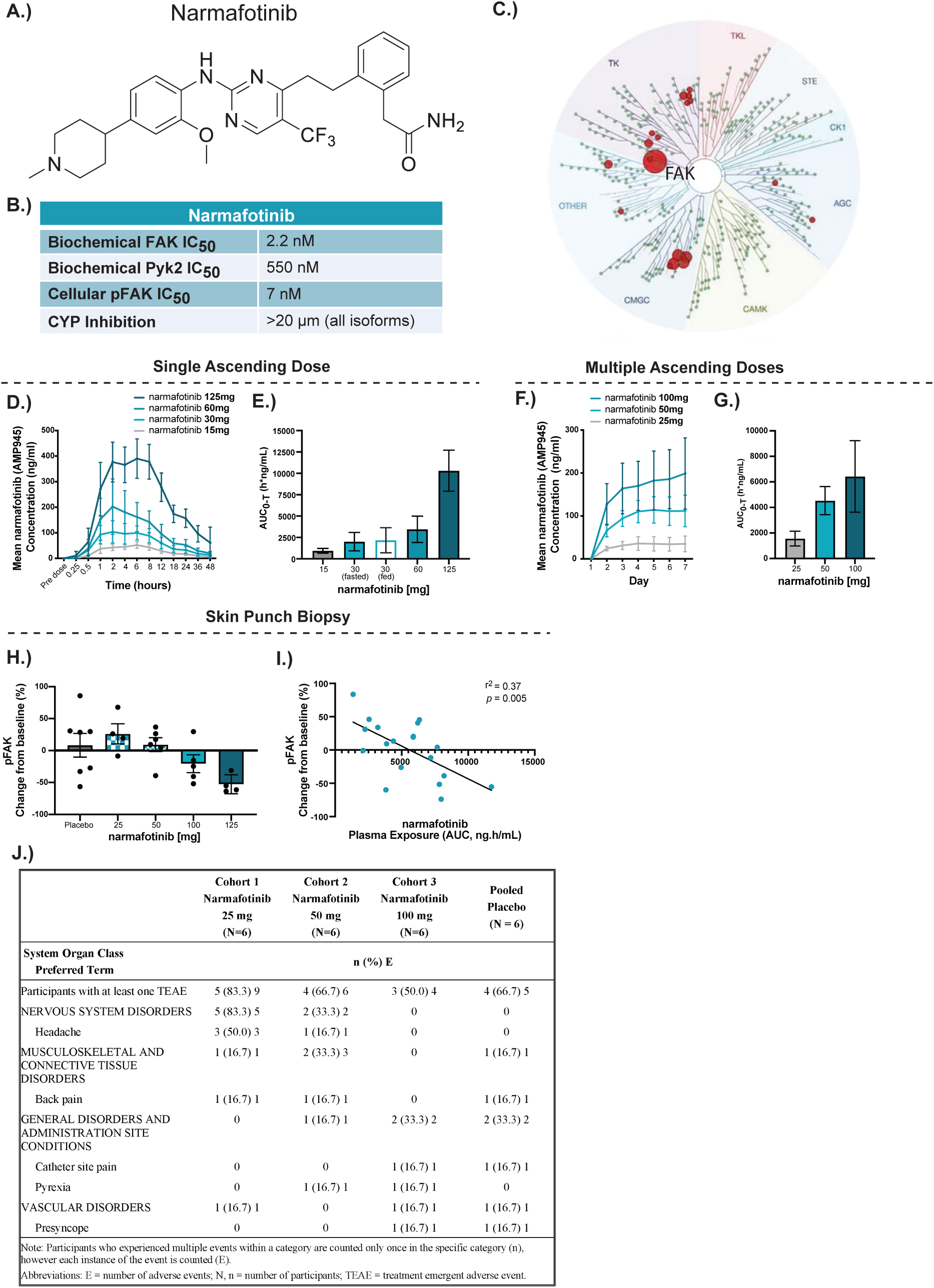
Narmafotinib, a highly selective FAK inhibitor. (**A.**) Chemical structure of narmafotinib. (**B.**) Determination of IC_50_ values for narmafotinib against FAK, and Pyk2, cellular pFAK inhibition (n=12). (**C.**) Kinase array data for narmafotinib against 468 kinases (including 403 non-mutant kinases). Circle size corresponds to % control. (**D-G**) Pharmacokinetics of narmafotinib in healthy volunteers following SAD (**D,E**) or MAD administration of narmafotinib (**F,G**). (**H.**) pTyr-397-FAK levels in skin punch biopsies from healthy volunteers. (**I.**) Linear relationship between change in FAK activity from baseline and AUC_0-inf_ following narmafotinib dosing. AUC_0-T_: Area under the plasma concentration-time curve from the time of dosage administration over the dosage interval. n ≥ 6. (**J.**) Summary of TEAEs in Phase Ia AMP945-101 trial. For (**E,G**) AUC graphs depict geometric mean ± geometric %CV. For (**D,F**) results are mean ± SD. pFAK changes from baseline r^2^ and p-value determined by simple linear regression fit, assessed for departure from zero (baseline).

### Narmafotinib reduces fibrosis and PDAC invasion

3D organotypic matrices enable the assessment of matrix remodeling to assess the anti-fibrotic potential of narmafotinib *in vitro* (*12, 13, 48*). Telomerase-immortalised fibroblasts (TIFs) were initially embedded into acid-extracted fibrillar collagen and allowed to contract collagen in the presence of narmafotinib over 12 days (Figure 3A (*12, 13, 48*)). Here, narmafotinib significantly reduced matrix contraction (Figure 3B,C) independent of changes in fibroblast proliferation (Ki67-positive) and apoptosis (cleaved caspase-3-positive, Figure S4A,B). SHG imaging was used to visualise fibrillar collagen, where we observed reduced SHG intensity in narmafotinib-treated conditions compared to control (Figure 3D,E, Movie S2). This was confirmed using Picrosirius Red staining and polarised light microscopy of collagen birefringence (Figure 3F-I). Further analysis of high, medium, and low birefringence signal (indicating a range of high-to-low density fibres, respectively) showed a decrease in fibrillar collagen density upon narmafotinib treatment (Figure 3I, Figure S4C). We also confirmed the anti-fibrotic efficacy of narmafotinib in a panel of cancer-associated fibroblast (CAF) lines, where we observed that narmafotinib decreased CAF-mediated collagen matrix contraction and reduced SHG signal intensity in murine KPC-educated CAFs and human PDAC patient CAFs (Figure S4D-I). Collectively, these data show that narmafotinib can reduce ECM contraction and remodelling mediated by murine and human pancreatic CAFs.

**Figure 3.**
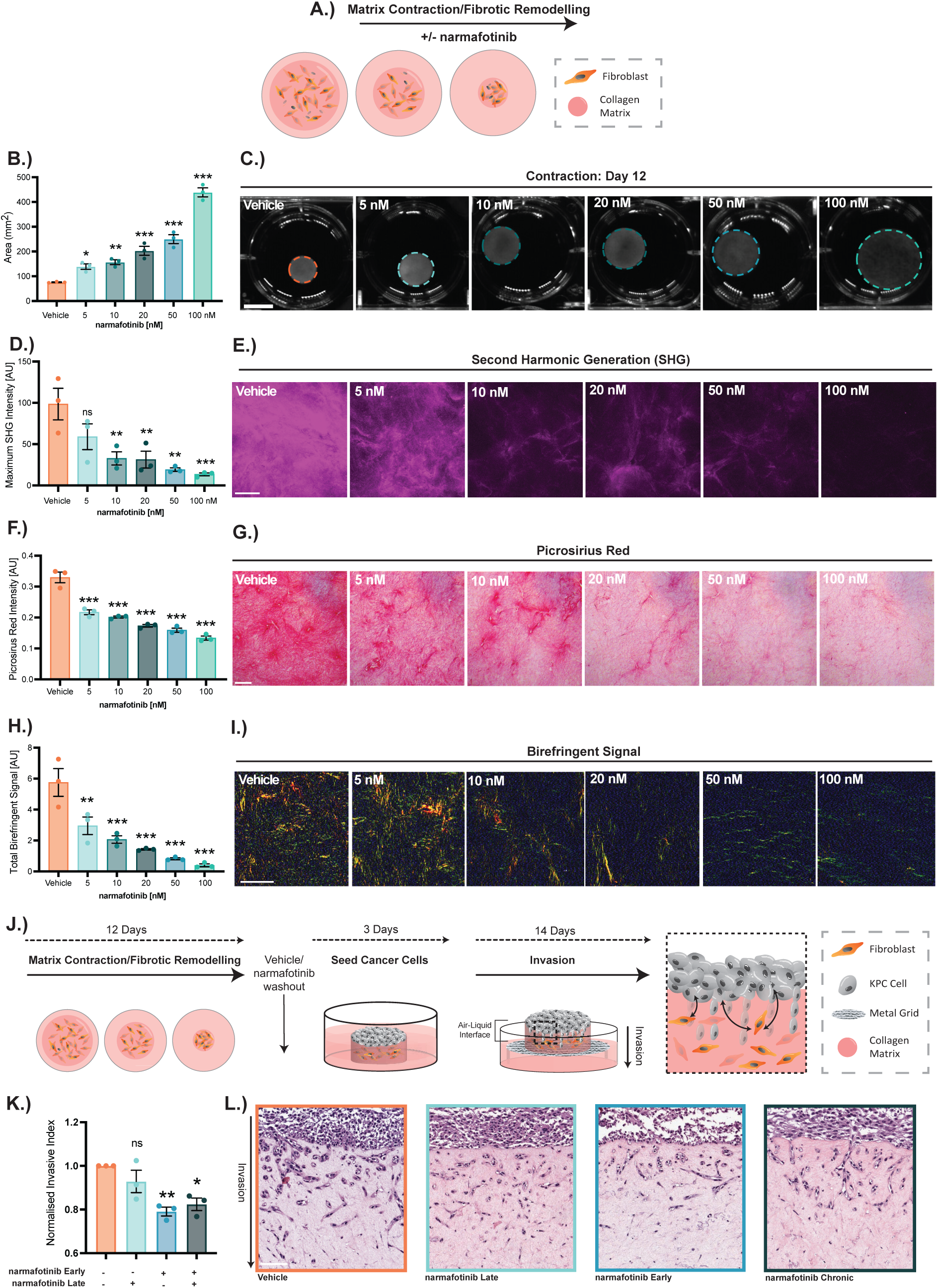
Narmafotinib reduces collagen remodelling and KPC cancer cell invasion. (**A.**) Schematic of 3D organotypic contraction assays. (**B,C**) Quantification (**B.**) and representative images (**C.**) of matrix contraction (Day 12, scale bar, 1 cm). (**D,E**) Quantification (**D.**) and representative images (**E.**) of SHG (Day 12). (**F,G**) Quantification (**F.**) and representative brightfield images (**G.**) of Picrosirius Red intensity (**F.**) in matrices (Day 12). (**H,I**) Quantification of total birefringence signal (**H.**) and representative birefringence images (**I.**) of Picrosirius Red stained matrices (Day 12). (**J.**) Schematic of 3D organotypic invasion assays (see Figure S5A for treatment schedule). (**K,L**) Quantification (**K.**) and representative images (**L.**) of KPC cell invasion into 3D organotypic matrices upon treatment with vehicle or 20 nM narmafotinib during invasion, matrix contraction or matrix contraction and invasion (chronic treatment). Scale bar, 100 µm (**E,G,I,L**). n=3 biological repeats, with three matrices per repeat and three FOVs per matrix. Data represent means ± SEM, p-values determined using (**B,D,F,H**) ordinary one-way ANOVA with Dunnett’s multiple comparisons test or (**K**) one-sample t test. Unless otherwise stated, significance compared to vehicle. ns, P≥0.05, *P<0.05, **P<0.01, ***P<0.001.

Given the anti-fibrotic effect of narmafotinib, we next assessed the effect on subsequent cancer cell invasion into 3D organotypic matrices using PDAC cells isolated from the genetically engineered KPC mouse model of PDAC (*Pdx1-Cre; LSL-Kras^G12D/+^; LSL-p53^R172H/+^* (*49, 50*)). Here, cancer cells were allowed to grow on TIF-contracted collagen matrices for 3 days. Matrices were then transferred onto a grid to generate an air-liquid interface and expose the KPC cells to a chemotactic gradient, to which they invaded (Figure 3J, Figure S5A (*12, 13, 48*)). Organotypic matrices were treated with vehicle or narmafotinib during fibroblast-mediated collagen contraction only (early ‘priming’), during KPC cell invasion only (late treatment) or both in a chronic setting (Figure 3J,K, Figure S5A). Interestingly, late treatment with narmafotinib during cancer cell invasion was not sufficient to reduce invasion (Figure 3K,L, orange *versus* light green). However, early priming during ECM remodelling significantly decreased cancer cell invasion (Figure 3K,L, orange *versus* light blue), which was not improved following chronic treatment (Figure 3K,L, light blue *versus* grey). Ki67 and cleaved caspase-3 staining showed that the decrease in cancer cell invasion was independent of changes in cell proliferation and apoptosis (Figure S5B,C). Moreover, when organotypic matrices were de-cellularised to remove the fibroblasts prior to cancer cell invasion, we observed that both early narmafotinib priming during collagen contraction as well as late narmafotinib treatment during cancer cell invasion significantly reduces cancer cell invasion (Figure S5D-F). This indicates that narmafotinib can have effects on both cancer and stromal cells depending on the context. Further work could also assess whether narmafotinib reduces cancer cell invasion, for example by decreasing the formation of invasion-permissive ECM tracks as previously described (*24, 51–53*). Overall, these results suggest that short-term priming with narmafotinib during ECM remodelling is sufficient to impair PDAC cell invasion without a need for chronic treatment. Therefore, this priming approach could be combined with chemotherapies to maximise outcome, while minimising side effects associated with chronic treatments (*8, 12, 13, 54*). Thus, we sought to assess the effect of narmafotinib priming on PDAC fibrosis and progression in combination with chemotherapy *in vivo*.

### Narmafotinib priming reduces fibrosis and metastasis while also improving response to gemcitabine/Abraxane and FOLFIRINOX

As the organotypic data demonstrated that narmafotinib treatment during stromal remodelling reduced fibrosis, we investigated if short-term pulsed narmafotinib priming could reduce fibrosis *in vivo*. KPC cells expressing the well-validated, reversible FAK-FRET biosensor (*12, 27*) were implanted subcutaneously into nude mice to monitor FAK inhibition in live PDAC tumours by intravital imaging (Figure 4A,B). Once tumours were palpable, mice were treated with vehicle or 10 mg/kg narmafotinib for 3 days. Intravital FLIM-FRET imaging of the FAK biosensor in surgically exposed tumours (Figure 4A,B), at 4 h post-final dosage of narmafotinib, confirmed that short-term treatment was sufficient to reduce FAK activity in live tumours, as seen by a decrease in fluorescence lifetime (Figure 4C, FLIM images showing long lifetimes [active FAK] in yellow/green colours, and short lifetimes [inactive FAK] in blue). Additionally, we also observed a concomitant decrease in ECM remodelling following narmafotinib treatment as assessed by SHG and birefringence imaging (Figure 4D, Figure S6A,B).

**Figure 4.**
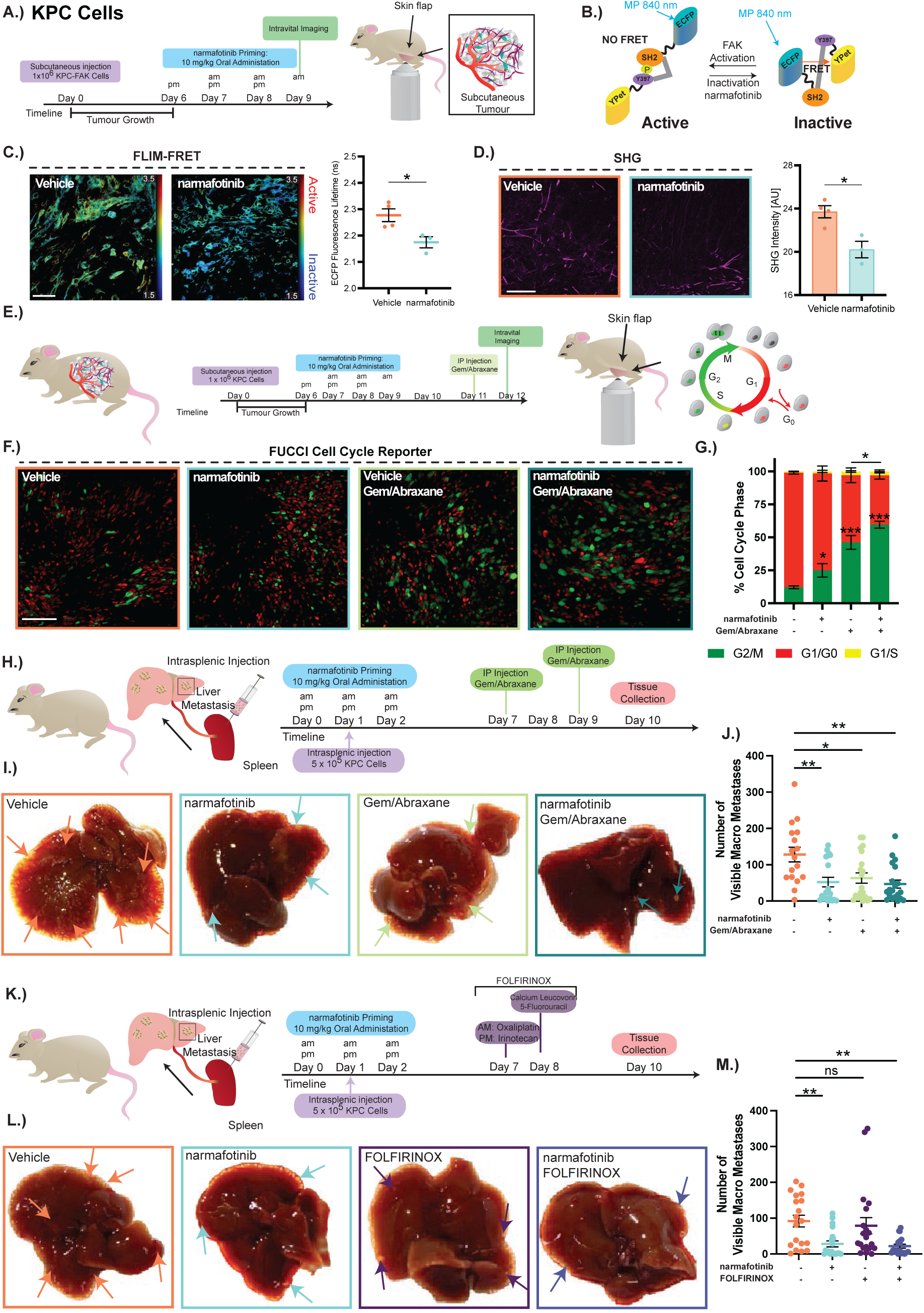
Narmafotinib priming improves KPC response to gemcitabine/Abraxane *in vivo*. (**A,B**) Timeline of KPC-FAK-FRET intravital imaging study (**A.**) using the Lyn-FAK-FRET biosensor (**B.**). (**C.**) Images and quantification of fluorescence lifetime (nanoseconds) in mice treated with vehicle or narmafotinib (10 mg/kg). n=50 cells per animal. (**D.**) Representative images and quantification of SHG in tumours treated with vehicle or narmafotinib. 5 FOV per animal, n=4 vehicle and n=3 narmafotinib animals (**C,D**). (**E.**) Timeline of KPC-FUCCI intravital imaging study. (**F,G**) Representative FUCCI images (**F.**) and quantification of cell cycle distribution (**G.**) in tumours treated with vehicle or narmafotinib followed by saline or gemcitabine/Abraxane. n=10 FOV per animal, 4 animals per treatment group. (**H-J**) Timeline (**H.**), representative liver images (**I.**) and quantification of visible liver macrometastases (**J.**) following intrasplenic KPC cell injection into mice treated with vehicle, narmafotinib, gemcitabine/Abraxane alone or narmafotinib prior to gemcitabine/Abraxane. n ≥ 17 mice per treatment group. (**K-M**) Timeline (**K.**), representative liver images (**L.**) and quantification of visible liver macrometastases (**M.**) following intrasplenic KPC cell injection into mice treated with vehicle, narmafotinib, FOLFIRINOX alone or narmafotinib prior to FOLFIRINOX. n ≥ 18 mice per treatment group. Arrows point at macrometastases in the liver tissue (**I,L**). Scale bar, 50 µm (**C.**), 100 µm (**D,F**). Results are mean ± SEM, p-values determined using (**C,D**) unpaired two-tailed t-test with Welch’s correction for unequal variance, (**G.**) two-way ANOVA with Tukey’s multiple comparisons test and (**J,M**) Kruskal-Wallis test with Dunn’s multiple comparisons. ns, P≥0.05, *P<0.05, **P<0.01, ***P<0.001.

Bi-directional cancer cell-ECM signalling has been shown to reduce response to chemotherapy (*7, 19, 20*). Given the anti-fibrotic effects of narmafotinib, we sought to assess the effect of narmafotinib priming on gemcitabine/Abraxane efficacy in live tumours. KPC cells were engineered to express the Fluorescence ubiquitin cell cycle indicator (FUCCI) reporter (KPC-FUCCI), which changes colour from green-to-yellow-to-red depending on the cell cycle phase (Figure 4E) (*26*). This allows for real-time examination of cell cycle progression or stalling in response to chemotherapy (*12, 26, 40*). Following establishment of subcutaneous KPC-FUCCI tumours, mice were treated with vehicle or 10 mg/kg narmafotinib for 3 days (Days 6-9) prior to a single treatment with gemcitabine/Abraxane or saline control on Day 11 (Figure 4E). Tumours were exposed using skin-flap surgery 24 hours post-chemotherapy for intravital imaging. Quantification of the FUCCI cell cycle reporter demonstrated that narmafotinib alone induced an increase in the proportion of cells in G_2_/M phase compared to control (Figure 4F,G, Movie S3). In line with its clinical application, gemcitabine/Abraxane alone also resulted in a higher proportion of cells stalling in G_2_/M compared to control. Importantly, this was further increased upon narmafotinib priming (Figure 4F,G, Movie S3), indicating an improvement in chemotherapeutic efficacy. Moreover, SHG imaging demonstrated reduced fibrillar collagen following narmafotinib treatment alone and in combination with gemcitabine/Abraxane (Figure S6C,D, Movie S4). Lastly, cleaved caspase-3, and Ki67 analysis revealed that narmafotinib priming prior to gemcitabine/Abraxane increased cell death and reduced cell proliferation, respectively, compared to gemcitabine/Abraxane only (Figure S6E-H), confirming that narmafotinib priming can be utilised to improve chemotherapeutic outcome in PDAC.

To evaluate the effects of this enhanced response to gemcitabine/Abraxane on survival, KPC cells were subcutaneously injected into mice. Upon palpable tumour formation, mice were enrolled into treatment cycles of narmafotinib priming for 3 days prior to gemcitabine/Abraxane (Days 7,10) until study endpoint (Figure S7A). Promisingly, narmafotinib priming prior to gemcitabine/Abraxane resulted in a significant increase in median survival compared to gemcitabine/Abraxane alone (Figure S7B). Analysis of collagen deposition by Picrosirius Red staining in survival endpoint tumours revealed that repeated treatments with gemcitabine/Abraxane induce fibrosis compared to vehicle, which was reverted to baseline levels upon priming with narmafotinib prior to gemcitabine/Abraxane (Figure S7C,D). Of note, while this subcutaneous KPC model is very aggressive, which likely contributes to overall shortened survival times and survival differences between the treatment groups (Figure S7A,B), we were able to show in this setting that narmafotinib priming prior to gemcitabine/Abraxane (Figure S7A,B) performs better than chronic narmafotinib treatment in combination with gemcitabine/Abraxane (Figure S7E,F, compare survival benefits highlighted for combination treatment compared to chemotherapy only between Figures S7B and S7F).

PDAC is a highly metastatic cancer type, where patients are predominantly diagnosed with advanced-stage disease (*1*). To assess potential additional effects of FAK inhibition *via* narmafotinib on liver metastasis, we injected KPC cells into the spleen of mice, which leads to rapid colonisation and metastatic outgrowth in the liver, as previously described (*12, 13, 55*). Here, mice were treated with narmafotinib for three days to mimic FAK priming while cells are transiting in the circulation prior to administration of gemcitabine/Abraxane or FOLFIRINOX (Figure 4H,K), which is a second standard-of-care chemotherapy option shown to impact PDAC patient survival (*4–6*). At study endpoint, livers were collected for quantification of liver metastases. Our analysis showed that narmafotinib alone, and in combination with either chemotherapy, reduced metastatic burden at the visible and histological level (Figure 4H-M and S8A-D) suggesting that narmafotinib can provide anti-metastatic benefits in addition to its effects on primary tumours.

Previous studies have highlighted the role of the immune system in PDAC progression and response to treatment (*8, 56*). In order to assess the effect of narmafotinib in combination with chemotherapy on immune cell populations, we derived primary syngeneic KPC cancer cells and KPC-educated CAFs from the genetically engineered KPC mouse model backcrossed for 10 generations onto the C57BL6/J background (Figure S9A), as previously achieved (*48, 49, 57*). RNA-sequencing confirmed that *α-Sma*, *Pdgfrα* and *Pdpn* were highly expressed in syngeneic KPC-educated CAFs, while *Cdh1*, *Pdx1* and *Krt19* were highly expressed in syngeneic KPC cancer cells (Figure S9B, Table S5). This was also confirmed using immunofluorescence (Figure S9C) and is in line with well-validated control KPC cancer cells and KPC-educated CAFs (Figure S9D) as previously described (*33, 48, 49*). We next co-injected syngeneic KPC cancer cells and KPC-educated CAFs orthotopically into the pancreas of immunocompetent C57BL/6 mice (Figure S10A) as described previously (*48*). Of note, combination of narmafotinib with gemcitabine/Abraxane was not assessed in this immunocompetent setting to avoid any potential immune response to Abraxane, which is paclitaxel bound to human albumin (*58*). We therefore assessed narmafotinib in combination with FOLFIRINOX (Figure S10A). Following tumour establishment, mice were enrolled into treatment cycles consisting of narmafotinib priming prior to FOLFIRINOX chemotherapy with timed endpoint tumour isolation following 4 cycles of treatment (Figure S10A). Interestingly, while the abundance of CD4^+^ T cells, FoxP3^+^ T cells and F4/80 macrophages was unchanged (Figure S10B-G), we observed a significant increase in CD8^+^ T cell abundance in tumours treated with narmafotinib and FOLFIRINOX compared to all other treatment groups (Figure S10H,I). This suggests that narmafotinib in combination with chemotherapy may boost anti-tumour immunity.

Overall, our data from the short-term KPC mouse models show that narmafotinib priming can reduce fibrosis and improve PDAC response to subsequent chemotherapy while also limiting metastasis. We next sought to assess this combination therapy in human PDAC tissue using patient-derived tumours from the APGI/ICGC cohort (*12, 13, 39–42*).

### Narmafotinib improves response to gemcitabine/Abraxane in patient-derived settings

PDAC is known to be a highly aggressive disease, with intratumoural cancer cell-ECM communication influencing therapy performance and outcome (*39, 41, 42, 59, 60*). To assess the impact of narmafotinib priming we utilised TKCC10lo patient-derived cells (PDCL) from the APGI cohort (*39–41*), which form highly fibrotic tumours (Picrosirius Red, Figure 5A, Table S6 showing complete mutational status (*61*)). Upon establishment of subcutaneous TKCC10lo tumours, mice were enrolled into treatment cycles (Figure 5B). In line with clinical PDAC management, gemcitabine/Abraxane chemotherapy resulted in an increased survival compared to control (Figure 5C, orange *versus* light green). Importantly, survival was further extended upon narmafotinib priming prior to gemcitabine/Abraxane compared to chemotherapy only (Figure 5C, light green *versus* dark green), demonstrating that this first-line priming approach can delay PDAC progression in human PDCL-based xenografts.

**Figure 5.**
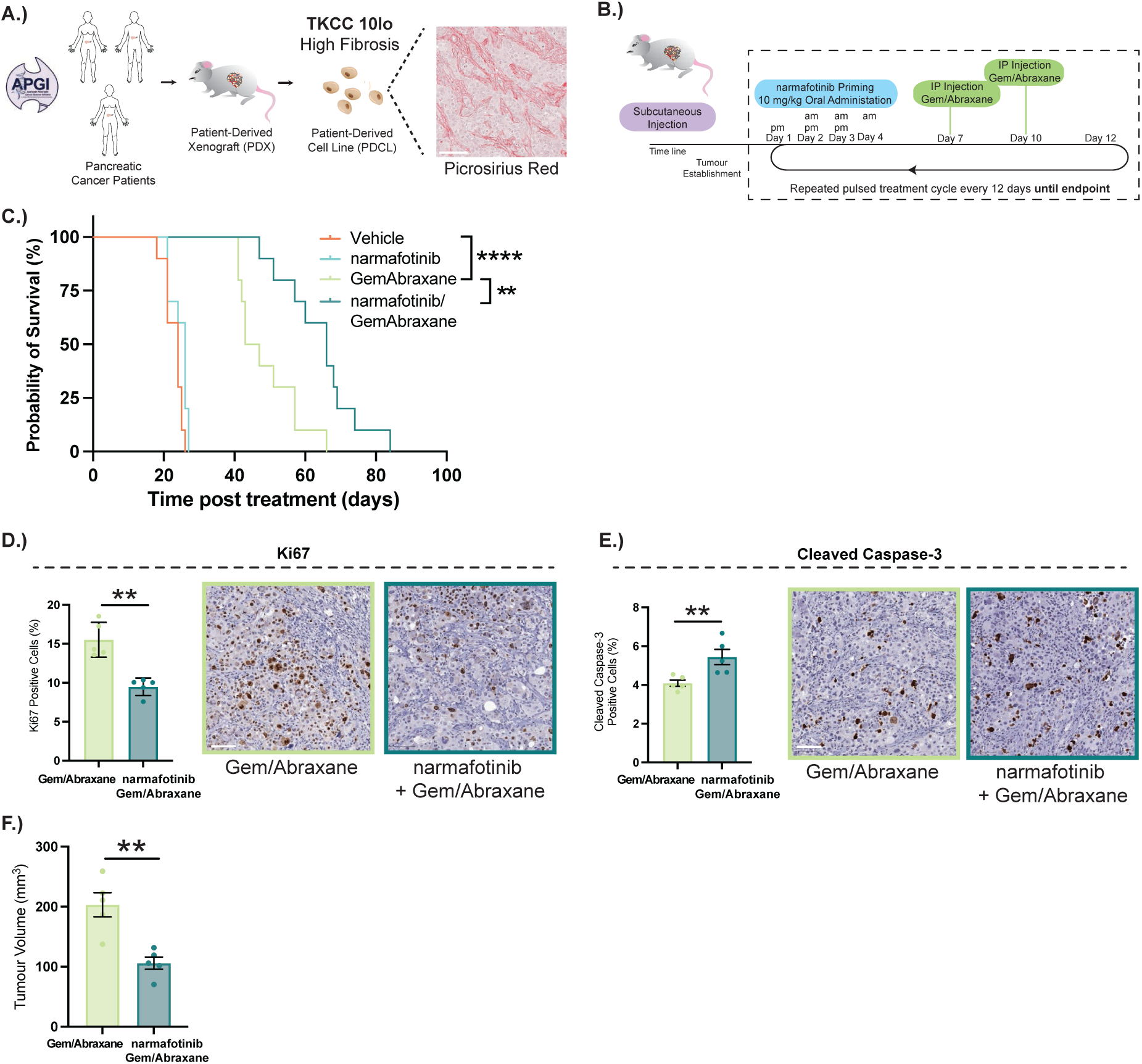
Narmafotinib priming prior to gemcitabine/Abraxane extends survival in subcutaneous patient-derived models. (**A.**) Patient-Derived Xenografts (PDXs) and Patient-Derived Cell Lines (PDCLs): TKCC10lo PDCL-based tumour, with high collagen levels (Picrosirius Red). (**B,C**) Timeline (**B.**) and Kaplan-Meier analysis of survival (**C.**) in mice with subcutaneous TKCC10lo tumours treated with vehicle, narmafotinib, gemcitabine/Abraxane alone or narmafotinib prior to gemcitabine/Abraxane. n=10 mice per treatment group. (**D-F**) Quantification and representative images of TKCC10lo tumours stained with Ki67 (**D.**) and cleaved caspase-3 (**E.**) as well as quantification of subcutaneous tumour size (**F.**) at timed endpoint after 3 treatment cycles with gemcitabine/Abraxane alone or narmafotinib prior to gemcitabine/Abraxane. Scale bar, 100 µm. n=5 mice per treatment group, 6 FOV per tumour. Results are mean ± SEM, p-values determined using (**C.**) log-rank Mantel-Cox test or (**D-F**) unpaired two-tailed t-test with Welch’s correction for unequal variance. ns, P≥0.05, *P<0.05, **P<0.01, ***P<0.001.

To further characterise the effects of narmafotinib priming on fibrosis and PDAC progression in this patient-derived setting, mice harbouring subcutaneous TKCC10lo tumours were subjected to short-term priming as before with timed study endpoints for all conditions (2 cycles for treatment groups without chemotherapy and 3 cycles for chemotherapy treatment groups). At timed endpoint, we observed a subtle decrease in tumour volume following narmafotinib alone compared to vehicle (Figure S11A). In addition, FAK activity was confirmed to be reduced compared to vehicle (Figure S11B), while SHG imaging confirmed reduced fibrosis in tumours treated with narmafotinib compared to vehicle (Figure S11C). This was confirmed by Picrosirius Red staining and birefringence imaging (Figure S11D,E). As previously observed in the KPC model (Figure S6E-H), tumours primed with narmafotinib prior to gemcitabine/Abraxane exhibited reduced proliferation (Figure 5D), enhanced apoptosis (Figure 5E) and a decrease in tumour growth (Figure 5F) compared to gemcitabine/Abraxane alone.

We next assessed changes in gene expression *via* RNA-sequencing to determine how narmafotinib may improve response to gemcitabine/Abraxane (Figure S12). Comparing the combination therapy with chemotherapy alone, we identified 142 significantly deregulated genes with 102 downregulated genes and 40 upregulated genes in tumours treated with narmafotinib prior to chemotherapy compared to chemotherapy alone (Figure S12A, Table S7). Gene set enrichment analysis (GSEA) revealed the upregulation of Hallmark gene sets ‘Myogenesis’ and ‘Protein Secretion’ in the combination therapy setting compared to chemotherapy alone in addition to previously reported tumour suppressors, such as *OTUD1* (*62*) and *SEMA3B* (*63*) (Figure S12A,B, Table S7). Downregulated transcripts included genes, such as *LOX*, which we and others have previously shown to contribute to increased cancer fibrosis and metastasis as well as reduced response to chemotherapy (*33, 64, 65*). Similarly, we found secreted proteins *SPON1* and *CHRDL2* to be significantly downregulated, which have been shown to increase collagen production and remodelling as well as metastasis in lung cancer (*66*) and to aid cancer cell survival during chemotherapy treatment in colorectal cancer (*67*). *FOXM1*, which has been demonstrated to regulate pro-desmoplastic gene expression programs in pancreatic stellate cells (*68*) and to mediate resistance to anoikis and gemcitabine in PC (*69, 70*), was also significantly downregulated. Similarly, downregulated expression of *BUB1* and *CEP55* aligns with their previously described roles in mediating chemotherapy resistance in pancreatic, lung, and breast cancer (*71–75*), and with the prolonged G_2_/M arrest observed by FUCCI imaging (Figure 4F,G).

Collectively, these data suggest that narmafotinib priming reduces the expression of known drivers of chemoresistance thus improving chemotherapy performance in line with the downregulation of Hallmark gene sets ‘E2F Targets’, ‘G_2_/M Checkpoint’ and ‘Mitotic Spindle’ identified *via* GSEA in this dataset (Figure S12B, Table S7). Moreover, using a consensus gene set for cancer hallmarks defined by Menyhart *et al.* (*76*), we observe a trending reduction in all cancer hallmarks in the combination treatment setting compared to chemotherapy alone (Figure S12C) with significant downregulation of ‘Genome Instability’ and ‘Resisting Cell Death’, both of which are highly enriched in PC (*76*), as well as ‘Reprogramming Energy Metabolism’ (Figure S12C). Further studies are needed to elucidate how these changes in gene expression contribute to the enhanced chemotherapy efficacy we observe in our PDCL-based *in vivo* model.

To further assess the efficacy of our combination therapy on transplanted whole patient-derived xenograft (PDX) tumours, we next implanted TKCC10lo-matched PDXs (from which TKCC10lo PDCLs were originally isolated) into mice and enrolled them into treatment cycles consisting of vehicle or narmafotinib priming prior to gemcitabine/Abraxane (Figure S13A,B). Here waterfall plots revealed decreased tumour growth as well as disease stabilisation and regression upon narmafotinib priming prior to gemcitabine/Abraxane compared to chemotherapy alone (Figure S13C). Promisingly, this resulted in a significant survival benefit for mice treated with the combination therapy in this matched PDX model (Figure S13D) emphasising that narmafotinib priming improves chemotherapy performance in both tumours based on PDCLs and PDX tumour samples. Given this promising result, we next assessed if narmafotinib priming could enhance the anti-tumour effects of FOLFIRINOX in patient-derived models.

### Narmafotinib improves response to FOLFIRINOX in patient-derived settings

FOLFIRINOX is a second standard-of-care chemotherapy option for PDAC patients (*4–6*). Critically, FOLFIRINOX is often associated with tolerability issues (*4*) and has not been widely assessed in combination therapies (*8*). We therefore wanted to test if we can overcome this obstacle, as our priming approach avoids drug combination at the same time as chemotherapy and could further shift the clinical benefits of FOLFIRINOX in PDAC (*4*). Upon establishment of subcutaneous TKCC10lo PDCL-based tumours, mice were subjected to treatment cycles consisting of narmafotinib priming (Days 1-4) prior to FOLFIRINOX (Days 8-9) until endpoint (Figure 6A,B (*4, 77*)). Consistent with its clinical use, FOLFIRINOX increased survival in this setting (Figure 6C, orange *versus* purple). Promisingly, mouse survival was further increased upon narmafotinib priming prior to FOLFIRINOX compared to FOLFIRINOX alone (Figure 6C, purple *versus* dark blue). To further characterise the effects of narmafotinib priming prior to FOLFIRINOX, TKCC10lo tumour-bearing mice were subjected to two treatment cycles, and timed endpoint samples were collected 24 h after administration of FOLFIRINOX or narmafotinib and FOLFIRINOX (Figure 6B). Here, we observed a subtle decrease in tumour volume when mice were primed with narmafotinib alone *versus* vehicle (Figure S11F). Moreover, while narmafotinib prior to FOLFIRINOX had no effect on proliferation (Figure 6D), it enhanced apoptosis (Figure 6E) and reduced tumour growth (Figure 6F) demonstrating that narmafotinib priming can also improve FOLFIRINOX efficacy.

**Figure 6.**
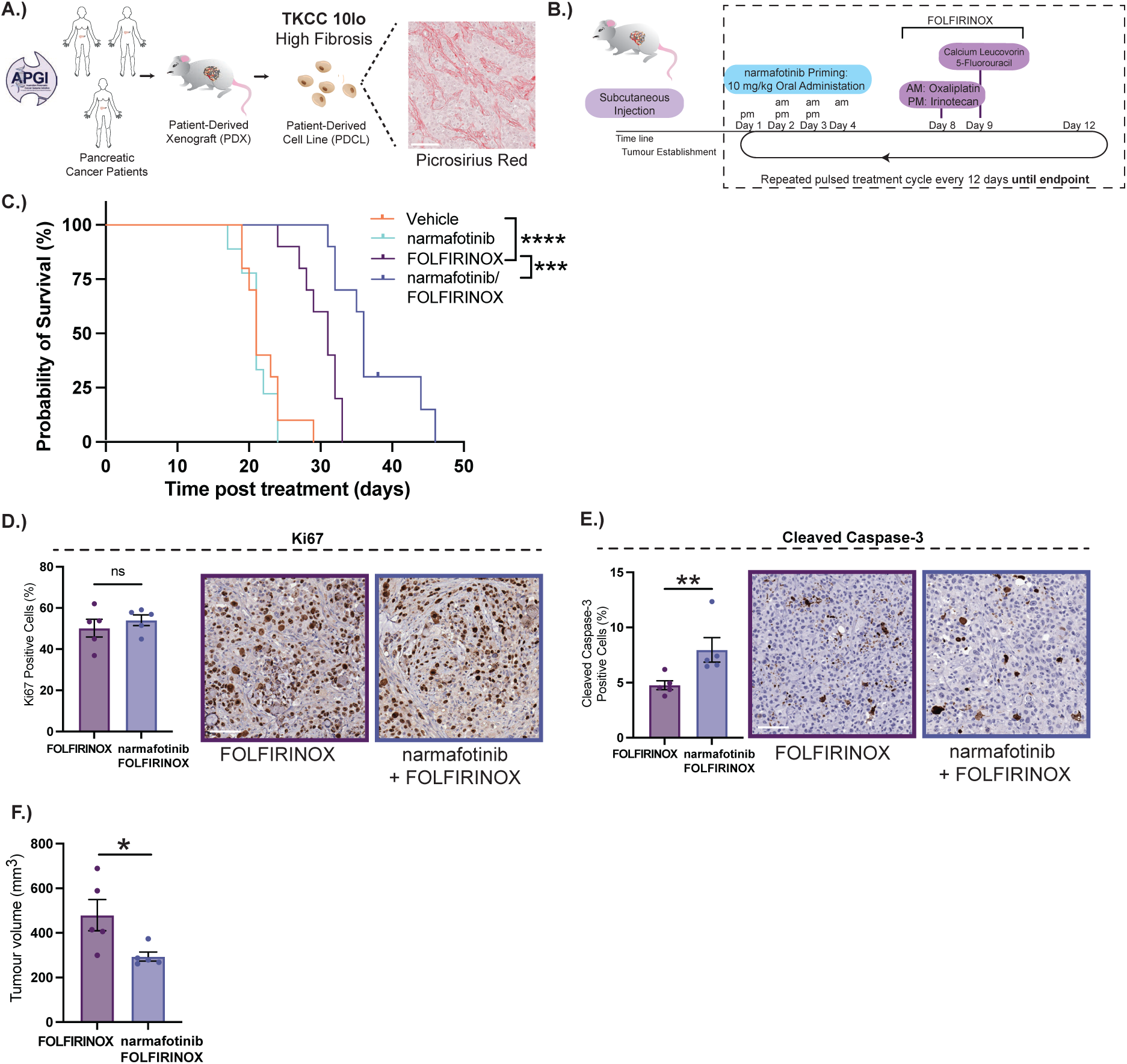
Narmafotinib priming prior to FOLFIRINOX extends survival in subcutaneous patient-derived models. (**A.**) Patient-Derived Xenografts (PDXs) and Patient-Derived Cell Lines (PDCLs): TKCC10lo PDCL-based tumour. (**B,C.**) Timeline (**B.**) and Kaplan-Meier analysis of survival (**C.**) in mice with TKCC10lo subcutaneous tumours treated with vehicle, narmafotinib, FOLFIRINOX alone or narmafotinib prior to FOLFIRINOX. n=10 mice per group. (**D-F**) Quantification and representative images of TKCC10lo tumours stained with Ki67 (**D.**) and cleaved caspase-3 (**E.**) as well as quantification of subcutaneous tumour size (**F.**) at timed endpoint after two treatment cycles with FOLFIRINOX or narmafotinib prior to FOLFIRINOX. Scale bar, 100 µm. n=5 mice per treatment group, 6 FOV per tumour. Results are mean ± SEM, p-values determined using (**C.**) log-rank Mantel-Cox test, (**D.**) an unpaired two-tailed t-test with Welch’s correction for unequal variance or (**E,F**) Mann-Whitney test for non-normally distributed data. ns, P≥0.05, *P<0.05, **P<0.01, ***P<0.001.

### Long-term orthotopic assessments of narmafotinib priming prior to chemotherapies in patient-derived settings

We next assessed the long-term effect of narmafotinib in combination with chemotherapy on survival in an orthotopic patient-derived model. Following orthotopic (intrapancreatic) injection of TKCC10lo cells expressing luciferase into mice (Figure 7A,B), tumour growth was monitored *via* whole-body In Vivo Imaging System (IVIS) imaging. Upon reaching an average flux of 1×10^9^ (Figure 7A,B (*13, 48*)), mice received pulsed treatment cycles consisting of 3 days narmafotinib priming prior to either gemcitabine/Abraxane (Days 7, 10, Figure 7A) or FOLFIRINOX (Days 8-9, Figure 7B). As expected, gemcitabine/Abraxane significantly extended survival in this long-term patient-derived model (Figure 7C, orange *versus* light green). Importantly, narmafotinib priming had an additional survival benefit over the gemcitabine/Abraxane only group (Figure 7C, light green *versus* dark green), demonstrating that narmafotinib priming improves gemcitabine/Abraxane efficacy and encouragingly, extends survival in an orthotopic patient-derived setting.

**Figure 7.**
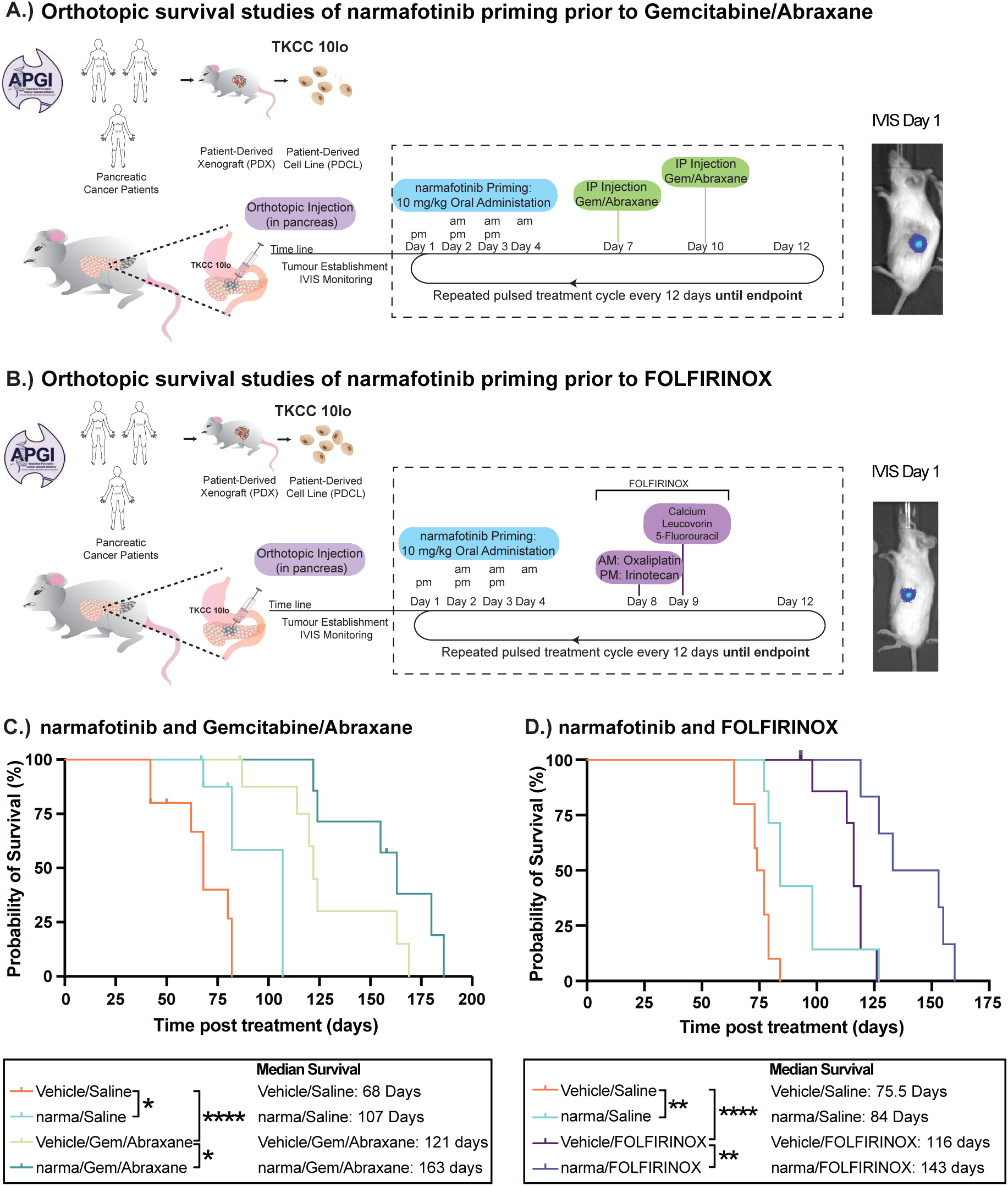
Narmafotinib priming prior to gemcitabine/Abraxane or FOLFIRINOX extends survival in long-term orthotopic patient-derived models. (**A,B**) Timeline for TKCC10lo orthotopic survival study with gemcitabine/Abraxane (**A.**) or FOLFIRINOX (**B.**). (**C.**) Kaplan-Meier analysis of survival in mice with TKCC10lo orthotopic tumours treated with vehicle (n=10), narmafotinib (n=9), gemcitabine/Abraxane alone (n=9) or narmafotinib prior to gemcitabine/Abraxane (n=8). (**D.**) Kaplan-Meier analysis of survival in mice with TKCC10lo orthotopic tumours treated with vehicle (n=10), narmafotinib (n=7), FOLFIRINOX alone (n=8) or narmafotinib prior to FOLFIRINOX (n=7). Groups compared using log-rank Mantel-Cox test. *P<0.05, **P<0.01, ****P<0.0001.

In the FOLFIRINOX cohort (Figure 7B) we observed that FOLFIRINOX extends survival compared to control (Figure 7D, orange *versus* purple). Importantly, narmafotinib priming prior to FOLFIRINOX, provided a survival benefit over FOLFIRINOX alone, demonstrating that narmafotinib priming can also improve FOLFIRINOX efficacy (Figure 7D, purple *versus* dark blue). These data indicate that FOLFIRINOX response rates in PDAC patients could also be enhanced by including anti-fibrotic priming with narmafotinib.

To further characterise the effects of narmafotinib priming prior to chemotherapy in the orthotopic setting, TKCC10lo cells were injected into the pancreas of mice and allowed to form palpable tumours (Figure S14A). As before, mice were subjected to treatment cycles consisting of narmafotinib priming prior to chemotherapy, and timed endpoint samples were collected (5 cycles for treatment groups without chemotherapy and 6 cycles for chemotherapy treatment groups, Figure S14A). As before, narmafotinib was shown to reduce FAK activity (Figure S14B,C). Analyses of Picrosirius Red brightfield and polarised light imaging confirmed that narmafotinib alone also reduced collagen deposition compared to control (Figure S14D-G). In agreement with our analysis in patient tissues (Figure 1A,B), we observed a significant increase in collagen deposition following chemotherapy treatment, which was decreased upon narmafotinib priming prior to chemotherapy (Figure S14D-G). Moreover, in line with its clinical applications, chemotherapy alone reduced cell proliferation (Figure S14H,I) with narmafotinib priming prior to chemotherapy significantly enhancing apoptosis (Figure S14J,K).

### Narmafotinib priming prior to gemcitabine/Abraxane reduces tumour progression in lowly fibrotic patient-derived settings

While our data demonstrate that narmafotinib priming prior to chemotherapy reduces tumour progression in an initially ECM-high patient-derived setting, we also aimed to assess this priming in an ECM-low patient-derived setting to determine whether narmafotinib priming could also counteract chemotherapy-induced fibrosis in this setting. Here, we subcutaneously injected TKCC2.1lo PDCLs, which form lowly fibrotic tumours (Picrosirius Red, Figure S15A, Table S6 showing complete mutational status (*61*)) into mice and, upon establishment of palpable tumours, enrolled the mice into treatment cycles (narmafotinib priming for 3 days prior to gemcitabine/Abraxane on Days 7 and 10, Figure S15B). At timed endpoint (2 cycles for treatment groups without chemotherapy and 6 cycles for chemotherapy treatment groups) narmafotinib treatment alone decreased FAK activity (Figure S15C) but did not affect tumour growth (Figure S15D). In line with our human PDAC patient data (Figure 1A,B) and data from the TKCC10lo model (Figure S14D-G), analysis of Picrosirius Red brightfield and polarised light imaging showed that gemcitabine/Abraxane alone increased fibrosis in this lowly fibrotic PDCL-based model, which was significantly reduced upon narmafotinib priming prior to chemotherapy (Figure S15E-H). Further analysis showed that narmafotinib had a small monotherapy effect on cell proliferation (Figure S15I,J) and enhanced cell apoptosis when given prior to chemotherapy (Figure S15K,L) leading to significantly reduced tumour size in the combination setting (Figure S15M).

Collectively, these data support first-line narmafotinib priming in PDAC as a clinically relevant strategy that not only enhances chemotherapy response rates without added toxicity but also has the potential to improve response rate to chemotherapy in both lowly and highly fibrotic settings.

## DISCUSSION

Reciprocity between cancer cells and the ECM plays an essential role in PDAC progression and response rates (*13, 15, 20, 35*). Disease management by combination therapies has increased in recent years; though the precise timing of multimodal targeting needs to be optimised to maximise benefit (*8, 54*). In this study, we demonstrate that neoadjuvant gemcitabine/Abraxane and FOLFIRINOX, both induce an early fibrotic reaction in PDAC patients, warranting further investigation as to whether ECM priming could be part of early PDAC treatment regimens to blunt the inevitable fibrosis occurring during clinical management. These findings highlight that the therapeutic impact of stromal targeting can be highly dependent on temporal context, where early, transient modulation of ECM architecture may be sufficient to reprogram tumour-stroma interactions, in contrast to chronic inhibition strategies that can promote adaptive resistance (*78*). Excitingly, our priming approach might lead to improved response over daily treatment and minimise potential side effects associated with multiple drug combinations, while avoiding known caveats associated with chronic anti-stromal therapies, which in the case of FAK inhibition have recently been shown to lead to stromal resistance (*13, 48, 78*). Our study introduces a temporally defined priming strategy, in which transient FAK inhibition is applied prior to chemotherapy to counteract treatment-induced fibrosis and enhance therapeutic response. Fine-tuned ECM-based priming could therefore facilitate a more nuanced anti-fibrotic therapy in PDAC as an alternative to chronic stromal targeting (*8, 15*).

Importantly, we present clinically relevant drug combinations, which could be used with other chemotherapy iterations in PDAC. For example, FOLFIRINOX-related treatment variations, such as FOLFOX, or FOLFIRI.3, as well as combinations of FOLFIRINOX variations with gemcitabine and/or Abraxane, such as FIRGEM, NAB-FOLFIRI, NALIRIFOX and NAB-FOLFOX, could be tested to investigate whether narmafotinib priming also improves response to these therapies (*6, 79*). Moreover, pulsed stromal priming in this way may be of similar benefit when used with immunotherapy approaches to improve response rates in PDAC (*14, 15, 80, 81*) as suggested by our data from immunocompetent mice where we observed a significant increase in CD8^+^ T cells upon combination of narmafotinib with FOLFIRINOX. These results may indicate a shift toward a more immunologically permissive tumour microenvironment, potentially reflecting enhanced T cell recruitment or intratumoural penetration/retention, and supports a future rationale for combining stromal priming with immune checkpoint blockade strategies.

Additional studies are needed to define the optimal timing and clinical context for the use of FAK inhibitors or other anti-fibrotic agents in combination with standard-of-care therapies, including their most effective placement within a combination treatment regimen and the overall treatment sequence. However, the concordance between our preclinical findings and early clinical outcomes from the first-line narmafotinib priming and gemcitabine/Abraxane ACCENT trial (NCT05355298) further supports the translational potential of this approach, particularly given the preservation of tolerability and absence of additive toxicity (*82, 83*), which has so far limited some combination strategies in PDAC.

In line with this, using 3D organotypic assays, we demonstrated that narmafotinib, has similar anti-fibrotic and anti-invasive efficacy when used in a priming setting compared to chronic assessment. We therefore investigated whether pulsed narmafotinib priming could improve chemotherapy response in PDAC, while avoiding potential side effects associated with concurrent drug exposure.

Similarly, narmafotinib priming may be assessed in other cancer types where multiple chemotherapeutics are given (*84, 85*); or where fibrosis is a key element hindering effective treatment, such as ovarian, breast, head and neck or squamous skin carcinomas (*57, 86–88*). Our results suggest that narmafotinib priming leads to increased accumulation of cells in G_2_/M phase of the cell cycle, rendering them more vulnerable to subsequent chemotherapy. We previously showed that anti-fibrotic ROCK1/2 inhibition also induces mitotic defects *in vivo* (*13*), sensitising cells to anti-mitotic Abraxane. Future studies may assess whether narmafotinib-mediated FAK inhibition targets additional aspects of PDAC progression beyond fibrosis due to the bi-directional signalling that FAK mediates at the tumour-ECM interface (*17, 18, 22, 23*). Interestingly, our RNA-seq analysis showed that narmafotinib priming with chemotherapy led to the downregulation of multiple genes previously associated with chemoresistance as well as cell cycle associated pathways, such as E2F targets, G_2_/M checkpoint and mitotic spindle programs, compared to chemotherapy alone, thus further supporting the assessment of narmafotinib priming with other clinically relevant chemotherapies. Moreover, and in line with previous studies (*14, 24*), we show that narmafotinib alone and in combination with chemotherapy also reduces PDAC metastasis suggesting additional benefits of this combination treatment beyond primary tumour targeting. Our data also suggest that patients who receive chemotherapy could benefit from transient narmafotinib priming independent of their initial tumour fibrosis status, as chemotherapy can increase fibrosis in both lowly and highly fibrotic tumours, which can be targeted with narmafotinib priming to reduce PDAC progression in either context.

Collectively, we have revealed a combination therapy approach to improve both gemcitabine/Abraxane and FOLFIRINOX response rates in PDAC. Importantly, the Phase I data demonstrates that narmafotinib is safe and well-tolerated supporting co-treatment with chemotherapies. This temporal separation of drug treatment offers an additional advantage to enhance the clinical benefits of both gemcitabine/Abraxane and FOLFIRINOX, which we have assessed in complementary *in vitro* as well as *in vivo* mouse, PDCL– and PDX-based models. While each of these models has its advantages and limitations (*8*), the efficacy benefit of narmafotinib priming is compelling. Building on these findings, a Phase Ib/IIa clinical trial (NCT05355298) has been initiated to evaluate a pulsed dosing regimen of narmafotinib priming with gemcitabine/Abraxane as first-line therapy in patients with unresectable or metastatic PDAC, which aligns with our preclinical work. This drug combination has been well tolerated by patients and early clinical data indicates favourable outcomes over historical data for gemcitabine/Abraxane alone (*2, 3, 82, 83*). In contrast to prior stromal depletion strategies, which have yielded mixed or adverse clinical outcomes, our data support a shift towards temporally defined ECM normalisation approaches. Moreover, data from our study warrants future, similar temporal assessments of this approach in FOLFIRINOX treatment regimens. Overall, these results position first-line stromal priming as a clinically translatable strategy to enhance therapeutic response and outcomes in PDAC.

## ABBREVIATIONS

AEs: adverse effects
APGI: Australian Pancreatic Genome Initiative
APMA: Australian Pancreatic Matrix Atlas
CAF: Cancer-associated fibroblast
ECM: extracellular matrix
ECG: electrocardiogram
ECOG: Eastern Cooperative Oncology Group
FAK: Focal Adhesion Kinase
FLIM: Fluorescence Lifetime Imaging Microscopy
FRET: Förster Resonance Energy Transfer
FOV: field of view
FUCCI: Fluorescence Ubiquitin Cell Cycle Indicator
ICGC: International Cancer Genome Cohort
IHC: immunohistochemistry
KPC: *Pdx1-Cre*
LSL-Kras^G12D/+^: LSL-Trp53^R172H/+^
MAD: multiple ascending dose
MSD: mesoscale discovery
PC: pancreatic cancer
PD: pharmacodynamic
PDAC: pancreatic ductal adenocarcinoma
PDCL: patient-derived cell line
PDX: patient-derived xenograft
PK: pharmacokinetics
RNA-seq: RNA-sequencing
ROI: region of interest
RT: room temperature
SAD: single ascending dose
SHG: Second Harmonic Generation
TEAE: treatment emergent adverse event
TIF: telomerase-immortalised fibroblast
TMA: tumour microarray
TWOMBLI: The Workflow of Matrix Biology Informatics

## FUNDING

This study was supported by the National Health and Medical Research Council (NHMRC), Australian Research Council (ARC), Cancer Council NSW, Cancer Institute NSW (CINSW), Cancer Australia, Tour de Cure, UNSW Medicine Cancer Theme, Perpetual Impact Philanthropy, St. Vincent’s Clinic Foundation, Sydney Catalyst (the Translational Cancer Research Centre of Central Sydney and Regional New South Wales), an Australian Cancer Research Foundation (ACRF INCITe) infrastructure grant and Suttons family and Len Ainsworth foundation philanthropy. This work was made possible by a PanKind Pancreatic Cancer Foundation Grant. P.T. is supported by the Len Ainsworth Fellowship in Pancreatic Cancer Research and a NHMRC Investigator Grant. K.J.M. (ECF1384), M.N. (ECF012), B.A.P. (ECF1309, CDF1266), S.R. (ECF1006), A.L.P. (CDF1167, CDF2307) and D.H. (ECF011) are supported by CINSW Research Fellowships. G.S. is supported by a CINSW Career Development Fellowship (CDF181166) and a Maridulu Budyari Gumal Sydney Partnership for Health, Education, Research and Enterprise [SPHERE] Cancer Clinical Academic Group Senior Research Fellowship (funded by CINSW Translational Cancer Research Capacity Building Grant 2021/CBG0003). G.S., P.A.P., M.P. and P.T. are supported by a CINSW Translational Program Grant (2020/TPG2100). C.R.C. and D.A.R. are supported by Baxter Family Scholarships. M.Tr. is supported by a White Walker Cancer Research PhD scholarship. V.T. is supported by an Elevate: Boosting Diversity in STEM (ATSE) PhD scholarship. M.P. is the recipient of Philip Hemstritch Pancreatic Cancer Fellowships and a Snow Medical Fellowship. The National Health and Medical Research Council (NHMRC) supported P.T. (2016930, 1160022, 1147364, 1136974, 1129401, 1105640, 1089497, 1043501), D.H. (2012837, 2028766), K.J.M. (2037787), B.A.P. (2019139, 2037824) and T.R.C. (2000937, 1140125, 1158590, 2033065). P.T. was supported by Cancer Council NSW (RG 14-08, RG 21-12, RG 24-06). T.R.C. was supported by a Cancer Institute NSW Career Development Fellowship (CDF171105), and Cancer Council NSW (RG19-09, RG23-11). O.J.S. and J.P.M. were supported by Cancer Research UK core funding to the CRUK Scotland Institute (A17196 and A31287) to O.J.S. laboratory (A21139) and to J.P.M. laboratory (A29996).

### ACKNOWLEDGEMENTS

We thank the staff at the following facilities at the Garvan Institute of Medical Research: Australian BioResources (ABR), Biological Testing Facility (BTF), Garvan Molecular Genetics (GMG), Garvan Genomics Platform (Flow Cytometry Hub, Sequencing Hub), Garvan Data Science Platform (DSP), Tissue Culture Facility under G. Lehrbach and R.J.L, the Imaging Platform under D.S.B. and the Histology and Biospecimen Facility under the leadership of A.Z. We also would like to thank the staff of the ACRF INCITe Centre under co-leadership of T.G.P. and P.T. We also thank C. Winchester for critical reading of the manuscript.

## CORRESPONDENCE

Corresponding author. (P.T.), (D.H) (K.J.M.)

## PATIENT AND PUBLIC INVOLVEMENT

Patients and/or the public were involved in the design and dissemination plans of this research.

## AUTHOR CONTRIBUTIONS

**Investigation, validation, and formal analysis:** K.J.M., C.R.C., D.A.R., L.M.C., N.E.P.M., S.E.M., V.L., A.E.H., A.M.H.T., M.N., A.M., J.St., B.A.P., N.K., S.R., K.G., M.Tr., V.M.T., S.H., V.J., A.A., D.S.B., C.V., X.Q.W., M.M.N., B.M., A.L.P., S.G., S.V.C., D.C.F., A.Z., M.Ta., A.D.S., APGI, APMA, A.J.C., S.M., C.S., S.L.L., D.R.C., G.S., P.A.P., L.D.G., R.J.L., A.N., N.P., A.J.G., A.V.B., J.Sa., C.E.C., T.R.J.E., L.C., L.G.H., O.J.S., J.P.M., T.G.P., Y.W., T.A.C., S.A.K., A.B., M.D., J.L., T.R.C., M.P., C.J.B., D.H. and P.T. **Conceptualisation and funding acquisition:** K.J.M., C.R.C., D.A.R., M.N., B.A.P., A.V.B., O.J.S., J.P.M., T.R.C., M.P., D.H. and P.T. **Writing and visualisation:** K.J.M., D.H. and P.T.

## DATA AND MATERIALS AVAILABILITY

All data needed to evaluate the conclusions in the paper are present in the paper and/or the Supplementary Materials. Materials from APGI and APMA can be provided by APGI and APMA pending scientific review and a completed material transfer agreement. RNA-seq data will be deposited in the GEO repository.

## ETHICS STATEMENTS

Human Ethics approval for acquisition of data and biological material was obtained from human research ethics committees from Sydney Local Health District (RPA Zone) Human Research Ethics Committee X16-0293, University of Melbourne Health Science Human Ethics Subcommittee: 1748955, North Shore Private Hospital Ethics Committee: NSPHEC 2016-016 and Garvan Institute of Medical Research:1627. All experiments using patient-derived CAFs were approved by UNSW Sydney Human Ethics Committee (approval HC180973) and were performed in accordance with the relevant guidelines and regulations. All patients provided written informed consent.

Animal experiments were conducted in accordance with the Garvan/St. Vincent’s Animal Ethics Committee guidelines (19/10, 19/13, 22/08, 22/09, 22/10, 25/10, 25/11, 25/12) and in compliance with the Australian code of practice for care and use of animals for scientific purposes.

