## Supplementary_Material_Figures for "Pulsed priming with the FAK inhibitor narmafotinib enhances both gemcitabine/Abraxane and FOLFIRINOX chemotherapy response in pancreatic cancer"

**This PDF File includes:**

Supplementary Materials and Methods

Legends for Supplementary Tables S1 to S7

Supplementary Tables S8 and S9

Legends for Supplementary Figures S1 to S15

Legends for Supplementary Movies S1 to S4

Consortium Members of the APGI and APMA

References

#### Materials and methods

***Statistical Analysis:*** Statistical analysis was performed using GraphPad Prism (GraphPad Software, Inc., CA) with significance given as; ns p≥0.05, *p<0.05, **p<0.01, ***p<0.001 and ****p<0.0001. Following normality confirmation using a Shapiro-Wilk test, data were analysed using an un-paired t test with Welch's correction for normally distributed data, or a Mann-Whitney test for non-normally distributed data. For data normalised to 1, a one-sample t-test was performed. FUCCI cell cycle data analysis was assessed using a two-way analysis of variance (ANOVA) with Tukey correction for multiple comparisons. Kaplan-Meier curves were compared using a log-rank Mantel-Cox test. For RNA-seq, p-values were adjusted using the Benjamini-Hochberg procedure, with adjusted $P < 0.05$ and a fold change of >1.25 considered significant. For association of pY397-FAK with clinico-pathological features in the Australian Pancreatic Genome Initiative (APGI) cohort of PC patients p-values were determined using Chi-square test. For all other datasets, p-values were determined by ordinary one-way ANOVA with a Tukey correction for multiple comparisons for normally distributed data or a Kruskal-Wallis test with Dunn's multiple comparison for non-normally distributed data.

***Human Ethics:***

Human Ethics approval for acquisition of data and biological material was obtained from human research ethics committees from Sydney Local Health District (RPA Zone) Human Research Ethics Committee X16-0293, University of Melbourne Health Science Human Ethics Subcommittee: 1748955, North Shore Private Hospital Ethics Committee: NSPHEC 2016-016 and Garvan Institute of Medical Research:1627. Additional information on patient sex and age, PDAC stage and pathological classification (TNM) as well as neoadjuvant chemotherapy and treatment duration of the Royal North Shore Hospital/Australian Pancreatic Matrix Atlas (APMA) cohort (*1, 2*) is provided in Table S1. Patient information on the APGI and International Cancer Genome Cohort (ICGC) cohort has been described previously (*3-5*) with mutational status of the patient-derived models derived from whole-genome sequencing detailed in Table S6 (*3*). All experiments using patient-derived CAFs were approved by UNSW Sydney Human Ethics Committee (approval HC180973) and were performed in accordance with the relevant guidelines and regulations. All patients provided written informed consent. All patient-derived CAFs tested negative for mycoplasma monthly.

***Animals:*** Animal experiments were conducted in accordance with the Garvan/St. Vincent's Animal Ethics Committee guidelines (19/10, 19/13, 22/08, 22/09, 22/10, 25/10, 25/11, 25/12) and in compliance with the Australian code of practice for care and use of animals for scientific purposes. Mice were kept in individually ventilated cages on a 12 h light/dark cycle and fed *ab libitum.* In this manuscript, we utilized subcutaneous, intrasplenic and orthotopic xenograft PC models. The subcutaneous KPC model is a rapid model, which we and others previously optimized and employed for rapid assessment of early tumour growth and response to treatments (*6-8*), while the other models were optimised to monitor tumour progression, metastasis and response to treatment.

Determination of the number of mice required to detect statistical significance was performed in line with 3Rs requirements. For intravital imaging studies we enrolled 3-5 mice per group. For survival studies 10 mice per group were enrolled. For timed endpoint studies we enrolled at least 5 mice per group. For intrasplenic studies, 20 mice per treatment group were enrolled. Mice were censored in the event of non-tumour-related complications. Randomisation *via* a random number generator was used to allocate mice to treatment groups in an un-biased manner. Mice were measured and treated at random to reduce any potential confounding factors. Tissues analysis was performed on blinded samples where mouse identification and treatment assignment were withheld.

***Phase Ia clinical trial AMP945-101:*** Amplia Therapeutics Phase Ia clinical trial AMP945-101 (ACTRN12620000894998) was a randomised, double blind, placebo-controlled study of the safety, tolerability and pharmacokinetics of single and repeat doses of narmafotinib administered orally to healthy adult volunteers. This first-in-man study assessed ascending once daily single doses of narmafotinib (AMP945 15, 30, 60, 125 mg), a food effect cohort (30 mg) and multiple ascending once daily doses for 7 days (25, 50, 100 mg). A total of 56 volunteers were dosed at Nucleus Networks (Melbourne Australia). Plasma and urine samples were analysed to measure concentrations of narmafotinib at Agilex (Adelaide, Australia). Plasma pharmacokinetic analyses were conducted to determine $AUC_{0\text{-}inf}$, $AUC_{0\text{-}t}$, $AUC_{0\text{-}24}$, $C_{max}$, $T_{max}$ and $t_{½}$. Safety and tolerability were assessed by the nature, incidence, and severity of adverse events (AEs), withdrawals, physical examination findings, vital signs (blood pressure, heart rate, respiratory rate, temperature), 12-lead electrocardiograms (ECGs), concurrent medications, semen sample analysis, and safety laboratory test results. The pharmacodynamic biomarker autophosphorylation of pY397-FAK was measured from skin using the Mesoscale Discovery (MSD) platform.

***AlphaScreen Assay:*** The primary inhibitory potency of narmafotinib was assessed against recombinant FAK (residues 411-688) using an AlphaScreen assay to detect the FAK catalysed phosphorylation of a synthetic peptide. The inhibitory activity of narmafotinib was measured at 11 concentrations in duplicate and assay performance was monitored by the routine inclusion of the pan-kinase inhibitor staurosporine and PF-562271. All assays were performed with the

ATP concentration fixed at $K_m$ (80 μM). In this assay narmafotinib exhibited high levels of inhibitory potency (IC50 = 2.2 nM, n = 12) and was similar to PF-00562271 (IC50 = 1.1 nM, n = 39). It should be noted IC50 values in the 1-2 nM range are at the theoretical lower limit of measurement for this kinase assay; however, the measured values for narmafotinib were consistent with time and compound batch.

***Narmafotinib Kinase Screen:*** Narmafotinib was screened against a panel of 468 kinases (including 403 non-mutant kinases) at Eurofins DiscoverX Corporation (San Diego, USA) using the KINOMEscan™ assay technology. For a detailed description of this assay technology see (*9*).

A Selectivity Score (S-score, Table S3) was calculated based on the acquired data and indicates that narmafotinib is highly selective for FAK. A selectivity score is a quantitative measure of compound selectivity calculated by dividing the number of kinases that the compound binds to by the total number of distinct kinases tested, excluding mutant variants. For a more detailed discussion of selectivity scores see (*10*).

The S-score value can be calculated using % Ctrl as a potency threshold and provides a quantitative method of describing compound.

- S(35) = (number of non-mutant kinases with % Ctrl <35)/(number of non-mutant kinases tested)
- S(10) = (number of non-mutant kinases with % Ctrl <10)/(number of non-mutant kinases tested)
- S(1) = (number of non-mutant kinases with % Ctrl <1)/(number of non-mutant kinases tested)

***Drug Treatment Schedules:*** Stock solutions of narmafotinib (Amplia Therapeutics Limited, in-kind) were prepared at 10 mM in DMSO for *in vitro* utilisation at a range of concentrations from 5 nM to 1 μM in cell culture medium with DMSO as vehicle control. For *in vivo* studies narmafotinib was dissolved in 0.5% w/v Hydroxypropyl methylcellulose, 0.5% benzyl alcohol, and 0.4% Tween 80 in sterile water, and was administered twice daily by oral gavage for 3 days, using a 22-gauge feeding tube (Instech Laboratories, FTP-22-25) coated in sucrose solution (24%, Sigma-Aldrich, S9378). In subcutaneous models, treatment of narmafotinib (10 mg/kg) commenced 6 days after KPC cell injection (average tumour volume of 30 $mm^3$), 23-29 days for TKCC10lo cells (average tumour volume of 50 $mm^3$), 38.5 days after TKCC10lo PDX implantation (average tumour volume of 180$mm^3$) and 8 days for TKCC2.1lo cells (average tumour volume of 75 $mm^3$) as per schematics (Figure 4A and E, 5B, 6B and Supplementary Figures S7A and S7E, S13B, S14A, S15B). In pancreatic orthotopic models, narmafotinib treatment began when tumours were palpable, and detectable by IVIS imaging (average flux of $1x10^9$) 4-weeks post-intrapancreatic injection for TKCC10lo cells (Figure 7A and B) or upon detection of palpable tumours in the syngeneic orthotopic KPC model (Supplementary Figure S10A). For intrasplenic studies mice were treated with narmafotinib from 1 day prior to intrasplenic KPC cancer cell injection for three days to mimic FAK priming while cells are transiting in the circulation prior to administration of gemcitabine/Abraxane or FOLFIRINOX (Figure 4H,K). Nab-paclitaxel (Abraxane®, Specialised Therapeutics) and gemcitabine (Selleck Chemicals) were dissolved in sterile saline and administered by intraperitoneal injection at 30 mg/kg and 70 mg/kg respectively. For FOLFIRINOX, in solution oxaliplatin (Clifford Hallam Healthcare Pty Ltd, 5 mg/kg), irinotecan (Clifford Hallam Healthcare Pty Ltd, 25 mg/kg), calcium leucovorin (Clifford Hallam Healthcare Pty Ltd 100 mg/kg) and 5-fluorouracil (Baxter Healthcare Pty Ltd, 25 mg/kg) or saline vehicle were administered by intraperitoneal injection. All treatments were administered as per the treatment schedule outlined in the corresponding Figures and Figure Legends.

***Patient-derived models:*** The Kinghorn Cancer Centre (TKCC) patient-derived xenografts (PDXs, n=45) and patient-derived cell lines (PDCLs, n=19) were obtained *via* the APGI and have been described previously (*3-5, 11*). PDXs were generated by implanting fresh tumour tissue pieces (1-2 x 1-3 $mm^3$) subcutaneously or orthotopically into NOD.Cg-$Prkdc^{scid}IL2rg^{tm1Wjl}$/SzAusb mice. For implantation of cryopreserved tumours, the fresh patient tumour was sectioned and cryopreserved in freezing media (90% xenograft media [RPMI 1640 + 10% FBS + 1.5% HEPES + 1% P/S] and 10% DMSO) for later thawing and transplantation. Engraftment rates were between 25 to 80% and PDXs were expanded *in vivo* for 2–6 months to obtain P1 PDXs. Once a P1 PDX tumour reached ~700–1500 $mm^3$, it was harvested and directly re-transplanted for expansion in later serial generations. Early-passage PDXs were cryopreserved in freezing media as above for subsequent expansion and treatment studies.

PDCLs were generated from PDX tumours grown in NOD.Cg-$Prkdc^{scid}IL2rg^{tm1Wjl}$/SzAusb mice. Following mechanical and enzymatic dissociation of PDXs (collagenase, Stem Cell Technologies, USA), PDCLs were grown in flasks coated with 0.2 mg/mL rat tail collagen (BD Biosciences, USA). Cultured cells were stained with biotinylated anti-mouse MHCI antibody (1:200 dilution; eBiosciences, US) in combination with Streptavidin AlexaFluor 647 secondary antibody (1:1,000; Invitrogen, US) and anti-mouse CD140a-PE antibody (1:300; BD Biosciences, US) and sorted on a FACS Aria III Cell sorter (BD Biosciences, US) to eliminate mouse stroma and enrich for epithelial cells. Short tandem repeat (STR) profiling (cellbankaustralia.com) was used to confirm PDCLs as unique. Mutational data for the PDCLs were obtained from sequencing data of the APGI project as part of the ICGC and are summarised in Supplementary Table S6 (*3*).

PDCLs are made available for research purposes upon reasonable request *via* the APGI website: https://www.pancreaticcancer.net.au/bioresource-pdcls/

***Isolation and validation of syngeneic KPC cancer cells and syngeneic KPC-educated CAFs:*** KPC mice (*Pdx1-Cre, LSL-Kras*$^{G12D/+}$*, LSL-Trp53*$^{R172H/+}$) (*12, 13*) were crossed with C57BL/6JAusb mice purchased from Australian BioResources for >10 generations to generate a cohort of KPC mice on C57BL/6JAusb background. All mice were bred at Australian BioResources, and genotyping was performed at the Garvan Molecular Genetics facility. Mice with the appropriate genotypes were enrolled into the study at ~6 weeks of age and weighed, palpated, and monitored once weekly until detection of a palpable tumour, after which monitoring was increased to at least 3x weekly. Mice were euthanised upon reaching the study endpoint, which included the development of signs of ascites, overnight weight loss of ≥10% or weight loss of ≥20% compared to the maximum body weight, hunching posture, and signs of pain. Tissues and organs were harvested from endpoint animals and primary cancer cell and CAF lines were generated as previously described (*7, 13*). Briefly, primary tumours were surgically excised, mechanically dissociated in a tissue culture hood using a scalpel blade and plated into tissue culture flasks in DMEM supplemented with 10% FBS and 1% penicillin/streptomycin at 37 °C in 20% $O_2$, 5% $CO_2$. Primary cancer cell and CAF cultures were then enriched and purified using a FACS Aria III Cell Sorter (BD Biosciences, USA) as previously described (*14*). Single-cell suspensions of 1 × $10^6$ cells were incubated with the anti-CD16/CD32 antibody (Mouse BD Fc Block; 1:200, BD Biosciences) in FACS buffer (PBS, 2% FBS, 2% HEPES) to block non-specific antibody binding. Cells were then incubated on ice for 10 min. Single-cell suspensions were then pelleted, washed with PBS and resuspended and incubated in FACS buffer containing anti-CD31-Biotin (1:100, BD Biosciences, clone 390) on ice for 20 min. Following centrifugation (1,200 rpm, 5 min) and removal of the supernatant, cells were washed and then re-suspended in FACS buffer containing the following cocktail: anti-EpCAM-PerCP/Cy5.5 (1:500, BioLegend®, clone G8.8), anti-CD45-APC-eFluor780 (1:500, Thermo Fisher, clone 30-F11), anti-CD140a-APC (1:100; BioLegend®, Clone: APA5), anti-Podoplanin-PE (GP38) (1:1,000, BioLegend®, Clone: 8.1.1) and BV786-streptavidin (1:500, BD Biosciences), for 20 min on ice. Cells were then washed twice in FACS buffer before being resuspended in FACS buffer containing DAPI (1:1,000; Invitrogen) to discriminate live and dead cells. CAFs were isolated according to the following cell surface markers: $CD140a^{+}/GP38^{+}/EpCAM^{-}/CD45^{-}/CD31^{-}/DAPI^{-}$. Cancer cells were isolated according to the following cell surface markers: $CD140a^{-}/GP38^{-}/EpCAM^{+}/CD45^{-}/CD31^{-}/DAPI^{-}$. Following isolation, cancer cells and CAFs were re-plated on a tissue culture dish and expanded. Cancer cells and CAFs were further characterised by RNA-sequencing (RNA-seq) and immunofluorescence.

For RNA-seq, RNA was extracted from cancer cells and CAFs using the QIAGEN RNeasy Mini Kit (no. 74104) according to the manufacturer's instructions (n=3 biological replicates per cell line). RNA concentration and integrity were assessed using the Agilent 4200 TapeStation system. Library preparation was performed using the TruSeq Stranded mRNA Kit according to the manufacturer's protocol (Roche) and paired-end sequencing was performed using the Illumina NovaSeq 6000. Sequence reads were aligned using STAR (*15*) to the mouse reference genome assembly GRCm39 and expression counts estimated using RSEM (*16*). Downstream analysis was performed in R (4.4.2) using Bioconductor packages DESeq2 (1.46.0) (*17*) for differential expression analysis. P-values were adjusted using the Benjamini-Hochberg procedure to control the false discovery rate.

For immunofluorescence, 10,000 cells per coverslip were seeded onto circular glass coverslips (Agar Scientific #AGL46R24) and incubated in media at 37 °C with 5% $CO_2$ for 72 h. PBS washed coverslips were fixed in 4% v/v paraformaldehyde (ProSciTech #C004) at room temperature (RT) for 10 minutes and washed twice again. Cells were then permeabilised using ice-cold methanol for 10 min. Washed coverslips were incubated in blocking buffer (2.5% BSA, 5% donkey serum and 0.02% glycine) for 1 hour at RT followed by an overnight incubation at 4 °C in either anti-Cdh1 antibody (BD Biosciences 610181) or anti-α-Sma antibody (Abcam ab5694) diluted 1:200 in blocking buffer. Coverslips were then washed and incubated in either Cy3 AffiniPure Donkey anti-Mouse IgG (Jackson ImmunoResearch #715165150) or Cy3 AffiniPure Donkey anti-Rabbit IgG (Jackson ImmunoResearch #711165152) diluted 1:500 in blocking buffer for 1 hour at RT. Washed coverslips were then stained with DAPI (Sigma-Aldrich #D9542) diluted 1:100 in PBS for 20 minutes at RT. Coverslips were then washed with distilled water and mounted onto glass slides using ProLong Diamond Antifade Mountant (Invitrogen #P36965) for imaging. Imaging was performed on the Leica Biosystems DMI6000 SP8 basic confocal microscope with 40x immersion oil objectives. Cy3 and DAPI were excited with 552 nm and 405 nm lasers, respectively, and the signal was acquired using PMT detectors set to an emission range of 560 nm – 665 nm and 411 nm – 476 nm, respectively.

***Cell Culture:*** Primary KPC cells have previously been isolated from end-stage *Pdx1-Cre, LSL-Kras*$^{G12D/+}$*, LSL-Trp53*$^{R172H/+}$ KPC mice (*12, 13, 18*), whilst Telomerase-immortalised fibroblasts (TIFs) were also generated previously (*2, 6, 19, 20*). KPC cancer cells, KPC-educated CAFs and TIFs were maintained in Dulbecco's Modified Eagle Medium (DMEM, high glucose, pyruvate, Gibco) supplemented with 10% Foetal Bovine Serum (FBS, Hyclone) and 10 mM HEPES (Gibco). Human CAF lines were grown in IMDM (Gibco) supplemented with 10% FBS and 4 mM Glutamine. TKCC10lo cells were maintained in 1:1 M199 media / Ham's F12 medium (Gibco) supplemented with 7.5% FBS, 15 mM HEPES, 2 mM Glutamine, 1x MEM vitamins, 25 ng/ml apo-transferrin, 0.2 IU/ml insulin, 6.5 mM Glucose, 40 ng/ml Hydrocortisone, 20 ng/mL EGF, 0.5 pg/mL Triiodothyronine and 2 μg/mL O-phosphoryl ethanolamine. TKCC2.1lo cells were maintained in RPMI (Gibco) supplemented with 10% FBS and 20 ng/mL EGF. All cells were cultured in the presence of Penicillin/Streptomycin

(100 U/mL and 100 μg/mL, respectively) and maintained at 37 °C and 5% $CO_2$. Experiments were conducted at 20% oxygen in a HeraCell 150i $CO_2$/$O_2$ incubator for KPC cancer cells, KPC-educated CAFs, human PDAC patient CAFs and TIFs and at 5% oxygen for TKCC10lo and TKCC2.1lo lines. All cell lines were confirmed to be mycoplasma free.

***Generation of stable cell lines:*** Generation of stable cell lines expressing the pPBDEST-Lyn-FAK biosensor was achieved by co-transfection of KPC cells with the pPBDEST-Lyn-FAK vector and pCMV-hyPBase transposase (obtained from the Welcome Trust Sanger Institute (*2, 21, 22*)), using Lipofectamine 3000 Reagent as per the manufacturer's instructions. KPC-FUCCI cells were also engineered to express the FUCCI cell cycle reporter (mKO2-hCdt1 and mAG-hGeminin, (*5*)) or Luciferase-GFP (pLV430G) using a 3rd generation lentiviral packaging system, as previously achieved (*6, 7*). Following transfection and transduction, respectively, positive cells were selected by Fluorescence Activated Cell Sorting (FACS).

***Organotypic Invasion assay:***

*Collagen Extraction:* Collagen I was extracted from rat tails as previously described (*2, 6, 7*). Briefly, once rat tails were defrosted, pronged tweezers were used to pull collagen tendons free of the epithelium and skeletal structure. Extracted tendons from 10-12 tails were then solubilised in 1500 mL of 0.5 M acetic acid for 48-72 h at 4 °C on a magnetic stirrer. The mixture was then filtered through a strainer to remove sheath, prior to precipitation with 10% (w/v) sodium chloride on a magnetic stirrer over 6-8 h. Once an opaque homogenous white solution was formed, the mixture was centrifuged for 30 min at 4 °C at 10,000 rpm. The resulting collagen precipitate was dissolved in 400-600 mL of 0.25 M acetic acid at 4 °C on a magnetic stirrer overnight and the solution dialysed in 4 L of 17.4 mM acetic acid for 4 days with acid refreshed every 12 h.

*Contraction Assays:* Organotypic matrices were generated as described (*2, 6, 7*). Briefly, 8x10$^4$ TIFs/matrix were embedded into acid-extracted rat tail collagen (~2.5 mg/mL) in the presence of 1x MEM, 8.8% FBS and neutralised with sodium hydroxide. Matrices were allowed to set at 37 °C before detachment and allowed to contract for 12 days in DMEM with 10% FBS, 10 mM HEPES and Penicillin/Streptomycin (100 U/mL and 100 μg/mL, respectively). For early treatment (priming), matrices were treated during contraction with vehicle, 5 nM, 10nM, 20 nM, 50 nM or 100 nM narmafotinib at Day 0 and refreshed at Day 6. For KPC-educated CAFs, 2x10$^5$ CAFs/matrix were embedded into collagen matrices (~1.5 mg/mL collagen) and allowed to contract the matrices as described above for TIFs. For human CAFs, 1x10$^5$ cells were embedded into collagen matrices (~1.5 mg/mL collagen) and allowed to contract in IMDM (10% FBS, 4 mM Glutamine). For TIF-cleared invasions, matrices were incubated with 10 μg/mL Puromycin on Day 12 for three days, prior to subsequent washing with PBS.

*Invasion Assays:* After 12 days of contraction, vehicle or 20 nM narmafotinib was washed out and 1x10$^5$ KPC cells were seeded onto contracted matrices. After 72 h of cancer cell growth, seeded matrices were moved to an air-liquid interface on a metal grid and KPC cancer cells were allowed to invade into the matrix for 14 days (Supplementary Figure S5A). For late treatment regimens, medium was supplemented with vehicle or 20 nM narmafotinib and refreshed 3 times per week during cell invasion (Supplementary Figure S5A). Chronic treatment involved narmafotinib treatment during both matrix contraction and during invasion (Supplementary Figure S5A). Matrices were then fixed in 10% neutral buffered formalin and processed for paraffin block embedding, sectioning and histological and immunohistochemistry (IHC) analysis (Garvan Histopathology and Biospecimen Facility). Using cell numbers calculated from H&E staining, cell invasion index was calculated as total number of cells invading divided by the cells on top of and within the matrix and normalised to average invasion index of vehicle cells.

$$Invasive\ Index = \frac{Number\ of\ invaded\ cells}{Cells\ on\ top\ of\ and\ within\ the\ matrix}$$

All organotypic assays were performed using at least 3 biological repeats and 3 replicates per repeat.

***ICGC and APGI Patient Data:*** For IHC analysis of patient TMAs from the APGI cohort tumour cores of deceased patients (3 per patient) with complete survival datasets (total 226) were scored for stromal structure/integrity (Picrosirius Red) and analysed using TWOMBLI (*23*). For pTyr-397-FAK, DAB intensity and coverage was manually scored (range 0-4) by 2 independent researchers under pathologist guidance in both stromal and tumour compartments to generate a combined score of both intensity and coverage. Kaplan-Meier curves were generated, using GraphPad Prism and a log-rank test performed to determine significance. In order to generate the data displayed in Table S2 patients were subdivided into low (combined score $\leq$ 2) *versus* high (combined score > 2) pTyr-397-FAK. Significance of correlation between scores for pTyr-397-FAK and clinico-pathological features was assessed using Chi-square test.

***Subcutaneous KPC Cell Injections for Intravital Imaging and Tumour Growth Studies:*** For intravital imaging studies, $1 \times 10^6$ KPC-FAK cells or KPC-FUCCI cells in PBS per mouse were injected into the rear flank of 7-9 week old female BALB/c-Fox1nuAusb mice whilst under anaesthesia (isoflurane 3%, $O_2$ 1 L/min, with continuous vacuum to remove excess isoflurane). Tumour size was monitored three times per week using callipers. Tumours were allowed to develop for 6 days to an average volume of ~25 $mm^3$ for KPC tumours before treatment schedules commenced. For KPC-FAK derived tumours, mice were treated with narmafotinib

(10 mg/kg) or vehicle twice daily for three days with the final treatment 4 h prior to skin flap surgery and subsequent intravital imaging (Figure 4A). For KPC-FUCCI derived tumours, mice were treated with narmafotinib (10 mg/kg) or vehicle twice daily for three days before a single treatment with gemcitabine (70 mg/kg) and Abraxane (30 mg/kg) or saline, followed by skin flap surgery and imaging 24 h post treatment (Figure 4E). For skin flap surgery, mice were terminally anaesthetised using a mix of 10 mg/kg xylazine and 50 mg/kg zoletil and with additional maintenance of anaesthesia (gaseous isoflurane 3%, $O_2$ 1 L/min, with continuous vacuum to remove excess isoflurane). Subcutaneous tumours were surgically exposed by a small incision around the tumour which was further expanded using blunt dissection to separate the epidermis and dermis from the peritoneal wall thereby creating a skin flap. The skin flap was expanded to a distance from the body suitable for intravital imaging. Once the tumours were surgically exposed, imaging was performed with mice restrained on a heated stage at 37 °C and maintained under anaesthesia, for a maximum of 40 min. Following intravital imaging tumours were formalin fixed for histological processing.

For tumour growth studies, $1x10^6$ KPC cells in PBS per mouse were injected into the rear flank of 7-9 week old female BALB/c-Fox1nuAusb mice whilst under anaesthesia (isoflurane 3%, $O_2$ 1 L/min, with continuous vacuum to remove excess isoflurane). Tumour size was monitored three times per week using callipers. Tumours were allowed to develop for 6 days to an average volume of ~25 $mm^3$ for KPC tumours before treatment schedules commenced. For narmafotinib priming studies, treatment schedules began with twice daily oral gavage of narmafotinib (10 mg/kg) or vehicle (commencing Day 1: PM, finishing Day 4: AM) followed by gemcitabine (70 mg/kg) and Abraxane (30 mg/kg) or saline administered by IP injection on Days 7 and 10, with the cycle recommencing after Day 12 (Figure S7A). For chronic narmafotinib treatment studies, treatment schedules began with twice daily oral gavage of narmafotinib (10 mg/kg) or vehicle (commencing on Day 1 twice daily AM and PM every day) followed by gemcitabine (70 mg/kg) and Abraxane (30 mg/kg) or saline administered by IP injection on Days 7 and 10, with the cycle recommencing after Day 12 (Figure S7A). At study endpoint (tumour volume of 350mm$^3$), mice were euthanised for tumour harvest.

***Intrasplenic KPC cell studies:*** KPC cells (5x10$^5$ cells in 50 µl of PBS) were injected into the spleen of BALB/c-Fox1nuAusb mice (anesthetized with 3% isoflurane, $O_2$ 1 L/min, with continuous vacuum to remove excess isoflurane) as previously described (*6, 24*). Briefly, a left subcostal incision was made through the skin and the peritoneum exposing the spleen, into which tumour cells were injected using a 29-gauge needle. The injection site on the spleen was then sealed with cyanoacrylate, and the peritoneal wall and skin were individually sutured. For analgesia, mice were treated subcutaneously with buprenorphine (0.075 mg/kg) and topically with bupivicaine (8 mg/kg). Mice were subjected to three treatments with narmafotinib (10 mg/kg) or vehicle by oral gavage before intrasplenic injection (Day 1), followed by three subsequent treatments (Figure 4H,K). For gemcitabine/Abraxane studies, gemcitabine (70 mg/kg) and Abraxane (30 mg/kg) or saline were administered by IP injection on Days 7 and 9 (Figure 4H) until study endpoint (Figure 4H). For FOLFIRINOX studies, in solution oxaliplatin (Clifford Hallam Healthcare Pty Ltd, 5 mg/kg), irinotecan (Clifford Hallam Healthcare Pty Ltd, 25 mg/kg), calcium leucovorin (Clifford Hallam Healthcare Pty Ltd 100 mg/kg) and 5-fluorouracil (Baxter Healthcare Pty Ltd, 25 mg/kg) or saline vehicle were administered by intraperitoneal injection on Days 7 and 8 until study endpoint (Figure 4K). On Day 10 mice, were euthanized for counting of visible metastases and livers were then fixed in formalin for histological processing. Quantification of metastatic area in the liver was performed on H&E sections using QuPath (*25*).

***Subcutaneous Injections of PDCLs:*** 1.5x$10^6$ TKCC10lo or 1x$10^6$ TKCC2.1lo cells in PBS/Matrigel (1:1) per mouse were injected into the rear flank of 7-9 week old female NOD.Cg-Prkdc$^{scid}$IL2rg$^{tm1Wjl}$/SzAusb mice whilst under anaesthesia (isoflurane 3%, $O_2$ 1 L/min, with continuous vacuum to remove excess isoflurane). Tumour size was monitored three times per week using callipers. Tumours were allowed to develop for 23-29 days to an average volume of 50 mm$^3$ for TKCC10lo or for 8 days to an average volume of 75 mm$^3$ for TKCC2.1lo cells before treatment schedules commenced.

Treatment schedules began with twice daily oral gavage of narmafotinib (10 mg/kg) or vehicle (commencing Day 1: PM, finishing Day 4: AM) followed by gemcitabine (70 mg/kg) and Abraxane (30 mg/kg) or saline administered by IP injection on Days 7 and 10, with the cycle recommencing after Day 12 (Figure 5B) until study endpoint. For FOLFIRINOX studies, in solution oxaliplatin (Clifford Hallam Healthcare Pty Ltd, 5 mg/kg), irinotecan (Clifford Hallam Healthcare Pty Ltd, 25 mg/kg), calcium leucovorin (Clifford Hallam Healthcare Pty Ltd 100 mg/kg) and 5-fluorouracil (Baxter Healthcare Pty Ltd, 25 mg/kg) or saline vehicle were administered by intraperitoneal injection on Days 8 and 9, with the cycle recommencing after Day 12 (Figure 6B) until study endpoint.

For survival studies with the TKCC10lo PDCL model, mice were euthanised for tumour harvest when reaching the study endpoint of a tumour volume of 600mm$^3$. For timed endpoint studies, mice were treated with narmafotinib (10 mg/kg) or vehicle twice daily for three days (commencing Day 1: PM and finishing Day 4: AM) over two cycles with the final treatment 4 h prior to tumour collection for timed endpoint mice (TKCC10lo and TKCC2.1lo PDCL models). In the context of gemcitabine/Abraxane chemotherapy, mice were treated twice daily via oral gavage of narmafotinib (10 mg/kg) or vehicle (commencing Day: 1 PM, finishing Day 4: AM) followed by gemcitabine (70 mg/kg) and Abraxane (30 mg/kg) or saline administered by IP injection on Days 7 and 10, for three cycles (TKCC10lo model) or for six cycles

(TKCC2.1lo model) with tumours collected 48 h after the final chemotherapy treatment. For FOLFIRINOX, in solution oxaliplatin (Clifford Hallam Healthcare Pty Ltd, 5 mg/kg), irinotecan (Clifford Hallam Healthcare Pty Ltd, 25 mg/kg), calcium leucovorin (Clifford Hallam Healthcare Pty Ltd 100 mg/kg) and 5-fluorouracil (Baxter Healthcare Pty Ltd, 25 mg/kg) or saline vehicle were administered by intraperitoneal injection on Days 8 and 9, for two cycles with tumours collected 24 h after the final treatment.

***Subcutaneous implantation of PDXs:*** Cryopreserved early-passage tumour pieces (~2 mm x 2 mm size) of the TKCC10lo PDX model were thawed and implanted subcutaneously into the rear flank of 7–9 weeks old female NOD.Cg-Prkdc$^{scid}$IL2rg$^{tm1Wjl}$/SzAusb mice whilst under anaesthesia (isoflurane 3%, $O_2$ 1 L/min, with continuous vacuum to remove excess isoflurane) in order to expand the PDXs. Tumour size was monitored three times per week using callipers. Once expanded PDXs reached a size of approximately 1 cm$^3$, mice were euthanised for harvest of PDX tumours, which were sectioned (~2 mm x 2 mm size) and directly re-implanted subcutaneously into the rear flank of 7–9 weeks old female BALB/c-Fox1nuAusb mice whilst under anaesthesia (isoflurane 3%, $O_2$ 1 L/min, with continuous vacuum to remove excess isoflurane) for treatment studies. Tumour size was monitored three times per week using callipers. Tumours were allowed to develop for approximately 38.5 days to an average volume of 180 mm$^3$ before treatment schedules commenced. Treatment schedules began with twice daily oral gavage of narmafotinib (10 mg/kg) or vehicle (commencing Day 1: PM, finishing Day 4: AM) followed by gemcitabine (70 mg/kg) and Abraxane (30 mg/kg) or saline administered by IP injection on Days 7 and 10, with the cycle recommencing after Day 12. At study endpoint (tumour volume of 1,500mm$^3$), mice were euthanised for tumour harvest.

***Orthotopic Injections:*** For orthotopic survival experiments, 7-9 week old female NOD.Cg-Prkdc$^{scid}$IL2rg$^{tm1Wjl}$/SzAusb mice were anesthetised (isoflurane 3%, $O_2$ 1 L/min, with continuous vacuum to remove excess isoflurane) and TKCC10lo-Luc cells (1x10$^6$ in 50 µL PBS/Matrigel (1:1) per mouse), were injected into the pancreas during open laparotomy (*2*). For orthotopic experiments in C57BL/6J mice, syngeneic KPC cancer cells and syngeneic KPC-educated CAFs (125 cancer cells and 375 CAFs [1:3 ratio for cancer cells and CAFs, respectively] in 50 µL PBS/Matrigel [1:1] per mouse) were injected into the pancreas during open laparotomy (*2, 7*). Briefly, a left subcostal incision was made through the skin and peritoneum exposing the pancreas, into which tumour cells were injected using a 29G needle. The peritoneal wall was then sutured using vicryl resorbable sutures and clipped. For analgesia, mice were treated subcutaneously with buprenorphine (0.075 mg/kg) and topically with bupivicaine (8 mg/kg). Treatment started when tumours were both palpable or visible by IVIS monitoring (average flux 1x10$^9$) for the TKCC10lo PDCL model or upon detection of a palpable tumour in the syngeneic model (C57BL/6J).

Treatment schedules began with twice daily oral gavage of narmafotinib (10 mg/kg) or vehicle (commencing Day 1: PM, finishing Day 4: AM) followed by gemcitabine (70 mg/kg) and Abraxane (30 mg/kg) or saline administered by IP injection on Days 7 and 10, with the cycle recommencing after Day 12 (Figure 7A). For FOLFIRINOX, treatment began with twice daily oral gavage of narmafotinib (10 mg/kg) or vehicle (commencing Day 1: PM, finishing Day 4: AM) followed by in solution oxaliplatin (Clifford Hallam Healthcare Pty Ltd, 5 mg/kg), irinotecan (Clifford Hallam Healthcare Pty Ltd, 25 mg/kg), calcium leucovorin (Clifford Hallam Healthcare Pty Ltd 100 mg/kg) and 5-fluorouracil (Baxter Healthcare Pty Ltd, 25 mg/kg) or saline vehicle on Days 8 and 9 (Figure 7B). Tumour growth was monitored using IVIS imaging once per week with additional close monitoring and palpation of all animals at least three times per week.

For survival studies, study endpoint was determined upon development of ascites, overnight weight loss of ≥ 10% or total weight loss of ≥ 20% over the entirety of the experiment, hunching posture or signs of pain. At endpoint animals were euthanised and the pancreatic tumour, liver, lungs, and spleen were harvested, visible metastases were quantified, and tissues formalin fixed for histological processing. Mice were excluded from study as censored events if endpoint occurred due to non-PDAC related symptoms. For timed endpoint PDCL studies, mice were treated with narmafotinib (10 mg/kg) or vehicle twice daily for three days (commencing Day 1: PM and finishing Day 4: AM) over five cycles with the final treatment 4 h prior to tumour collection for timed endpoint mice. In the context of gemcitabine/Abraxane chemotherapy, mice were treated twice daily via oral gavage of narmafotinib (10 mg/kg) or vehicle (commencing Day: 1 PM, finishing Day 4: AM) followed by gemcitabine (70 mg/kg) and Abraxane (30 mg/kg) or saline administered by IP injection on Days 7 and 10, for six cycles with tumours collected 48 h after the final chemotherapy treatment. For timed endpoint syngeneic KPC studies, mice were treated over four cycles twice daily via oral gavage of narmafotinib (10 mg/kg) or vehicle (commencing Day: 1 PM, finishing Day 4: AM) followed by in solution oxaliplatin (Clifford Hallam Healthcare Pty Ltd, 5 mg/kg), irinotecan (Clifford Hallam Healthcare Pty Ltd, 25 mg/kg), calcium leucovorin (Clifford Hallam Healthcare Pty Ltd 100 mg/kg) and 5-fluorouracil (Baxter Healthcare Pty Ltd, 25 mg/kg) or saline vehicle on Days 8 and 9 with tumours collected 24 h after the final treatment.

***RNA-seq of TKCC10lo PDCL subcutaneous tumours:*** TKCC10lo subcutaneous tumours were isolated at timed endpoint after 3 treatment cycles, snap frozen, and stored at −80 °C. RNA extraction was performed on 5 tumours per treatment arm using the QIAGEN RNeasy Mini Kit (no. 74104) as per the manufacturer’s instructions. The Agilent 4200 TapeStation system and a Qubit 3.0 Fluorometer (Thermo Fisher Scientific), were then used to determine

RNA concentration and integrity and library preparation was performed using the Illumina Stranded mRNA Library Preparation Kit according to the manufacturer's protocol. Paired-end sequencing was then performed using the NovaSeq S4 Flow Cell (300 cycles).

Nextflow pipeline nf-core/rnaseq (3.16.0) (*26*) was used to process RNA-seq data. Here, sequence reads were aligned to human reference genome assembly GRCh38 using STAR (*15*) and expression counts estimated using RSEM (*16*). Downstream analysis was performed in R (4.4.2) using Bioconductor packages DESeq2 (1.46.0) (*17*) for differential expression analysis and fgsea (1.32.4) for GSEA. For Cancer Hallmarks GSEA, the consensus list by Menyhart *et al.* was used with adaptation of data visualisation (*27*). P-values were adjusted using the Benjamini-Hochberg procedure to control the false discovery rate.

***Immunoblot:*** KPC cancer cells, KPC-educated CAFs, TKCC10lo PDCLs andTKCC2.1lo PDCLs were cultured for 2 h in the presence of vehicle or ascending concentrations of narmafotinib. Syngeneic KPC-educated CAFs were cultured for 2 h in the presence of vehicle, 50nM, 100nM, 200 nM, 300 nM and 500 nM narmafotinib. For protein extraction, cells were rinsed twice in PBS and lysates prepared in RIPA protein lysis buffer (50 mM HEPES, 1% Triton X-100, 0.5% sodium deoxycholate, 0.1% SDS, 0.5 mM EDTA, 50 mM NaF, 10 mM $Na_3VO_4$ and 1 x protease inhibitor cocktail (Roche)). Protein concentration was determined by Bradford assay and the lysate volumes were adjusted accordingly to a final concentration of 1 µg/µL. Protein separation was performed by gel electrophoresis using 4-12% Bis-Tis Protein Gels. Separated proteins were transferred onto PVDF membranes, which were blocked overnight at 4 °C in BSA dissolved in Tris-buffered saline, 0.1% Tween 20 (TBST). After rinsing with TBST membranes were incubated overnight at 4 °C in primary antibody solution (TBS/BSA). Antibodies and their respective dilutions are provided in Table S8. After rinsing with TBST, membranes were then incubated with horseradish peroxidase (HRP)-linked secondary antibody (1:5,000, diluted in 1% skim milk/TBST, GE Healthcare Limited), for 2 h at room temperature, and rinsed with TBST. Ultra-Enhanced Chemiluminescence (ECL) or ECL reagent were used to visualise HRP signal imaged on a Fusion FX (Vilber). Densitometry analysis of protein signal was performed in Image J (NIH). All Western Blot assays were performed using at least 3 biological repeats.

| **Antibody** | **Company** | **Product** | **Dilution** |
|---|---|---|---|
| Total FAK | BD Bioscience | 610088 | 1:500 |
| pTyr-397 FAK | Invitrogen | 44-625G | 1:500 |
| GAPDH (14C10) | Cell Signalling | 2118 | 1:10,000 |
| GAPDH (14C10) | Acris | ACR001P | 1:10,000 |

**Supplementary Table S8. Primary antibodies utilised for immunoblotting**

***Histology and Immunohistochemistry:*** Organotypic matrices and tumour tissues were fixed in 10% neutral buffered formalin then processed on the Leica Peloris 3 and paraffin-embedded using the Leica HistoCore Arcadia H&C. Sections were cut at a thickness of 4 μm onto positively charged slides and incubated in a 60 °C oven to maximise adhesion. H&E staining was performed using the Leica ST5010 Autostainer XL with Haematoxylin (Haematoxylin Harris non-toxic [acidified], Australian Biostain) and Eosin (Eosin Phloxine Alcoholic 1%, Australian Biostain).

For Picrosirius Red staining, the sections were dewaxed in xylene and rehydrated in graded ethanol washes. Following haematoxylin counterstaining sections were stained with 0.02% phosphomolybdic acid and 0.1% Picrosirius Red (Polysciences) for fibrillar collagen. Sections were then rinsed in acidified water and dehydrated in graded ethanol prior to cover slipping. Slides were scanned using an Aperio slide scanner. For APMA tumour sections, analysis of

Picrosirius Red coverage was performed using Image J (see Macro: Picrosirius Red Coverage), whilst an in-house MATLAB (Mathworks, US) script was used to analyse intensity and coverage of Picrosirius Red stained organotypic matrices and tumours.

For IHC on the Leica Bond RX, organotypic matrix and tumour sections were dewaxed using Bond Dewax Solution (Leica, AR2992) on a Leica Bond RX followed by heat-induced epitope retrieval (HIER) at 93 °C for organotypic matrix sections and 100 °C for tumour sections, with epitope retrieval solution 2 (pH=9, Leica AR9640) for 30 min, except for cleaved caspase-3 which was performed for 20 min. The details for primary antibody dilution and incubation time are provided in Table S9. IHC staining was carried out on the Leica Bond RX autostainer followed by haematoxylin counterstaining on a Leica Autostainer XL and cover slipping on a Leica Coverslipper (CV5030). Samples were scanned using an Aperio slide scanner, Hamamatsu NanoZoomer S210 Digital slide scanner or the Olympus Slideview VS-220 scanner. Cleaved caspase-3 and Ki67 stained organotypic matrix and tumour sections were analysed for positive and negative cells, and for pTyr-397-FAK (pTyr-397) DAB intensity was assessed using QuPath (*25*).

| **Antibody** | **Company** | **Product** | **Dilution** | **Incubation (minutes)** |
|---|---|---|---|---|
| Ki67 (SP6) Organotypic matrices | ThermoFisher Scientific | RM-9106-S1 | 1:500 | 60 |
| Ki67 (SP6) Mouse tissue | ThermoFisher Scientific | Ab4 Neomarker | 1:500 | 60 |
| Cleaved Caspase-3 | Cell Signaling | 9661 | 1:200 | 60 |
| pTyr-397-FAK | Abcam | ab39967 | 1:4,000 | 30 |

| | | | | |
|---|---|---|---|---|
| Human tissue | | | | |
| pTyr-397-FAK<br>Mouse tissue | ThermoFisher<br>Scientific | 700255<br>(31H5L17) | 1:4,000 | 60 |
| CD4 | Cell Signaling | 25229 | 1:100 | 60 |
| CD8 | Cell Signaling | 98941 | 1:200 | 60 |
| FoxP3 | Cell Signaling | 12653 | 1:400 | 60 |
| F4/80 | Cell Signaling | 70076 | 1:100 | 60 |

**Supplementary Table S9: Primary antibodies utilised for immunohistochemistry, with staining performed on the Leica Bond RX Autostainer**

***Imaging Techniques and Data Analysis in vitro and in vivo:***

*Polarised light imaging of Picrosirius Red staining:* Polarised light microscopy was performed on fixed, deparaffinised and rehydrated 4 μm sections stained with 0.1% Picrosirius Red (Polysciences, 29401-250). Polarised light signal of fibrillar collagen was collected using an Olympus U-Pot polariser and an Olympus U-ANT transmitted light analyser fitted to a DM4000 microscope (Leica). Quantification of the birefringence signal was performed using Image J. Briefly, Hue-Saturation Balance (HSB) thresholding was applied (high birefringence/red-orange 0>H<27 | 0>S<255 | 70>B<255, medium birefringence/yellow 28>H<47 | 0>S<255 | 70>B<255, low birefringence/green 48>H<140 | 0>S<255 | 70>B<255). Relative area of fibres was then calculated (0>H<140 | 0>S<255 | 70>B<255). Fibre orientation analysis was performed on polarised light images of Picrosirus Red stained tissue using an in-house ImageJ (NIH) macro (https://github.com/TCox-Lab/PicRed_Biref), as previously described (*28-31*). Briefly, structure tensors were derived from the local orientation and isotropic properties of pixels that make up collagen fibrils. Within each input image, these tensors were evaluated for each pixel by computing the continuous spatial derivatives in the x and y dimensions using a cubic B-spline interpolation. From this, the local predominant orientation was obtained. The peak alignment (measured in degrees) of fibres was then determined, and the frequency of fibre alignment calculated across different degree ranges spanning the peak alignment (i.e. peak alignment ± 5°, 15°, 30° and 45°).

*Second Harmonic Generation (SHG) imaging:* SHG imaging was performed on an inverted Leica DMI 6000 SP8 confocal microscope with a Titanium-Sapphire femtosecond laser (Coherent Chameleon Ultra II) excitation source, operating at 80 MHz and tuned to a wavelength of 880 nm. SHG intensity was recorded on an RLD-HyD at 440/20 nm. For organotypic matrices, 3 representative fields of view (512 px x 512 px) were imaged over a 3D z-stack (80 μm z-depth with a 2.52 μm step size). For tissue sections, 5 regions of interest of deparaffinised, rehydrated 4 μm unstained sections were imaged with a step size of 1.26 μm and 20 μm z-depth. SHG signal intensity was quantified using MATLAB (Mathworks, US). For tumour samples from the Royal North Shore Hospital/APMA cohort, samples were tile scanned using a Leica Stellaris 8 DIVE multiphoton inverted microscope. A Ti:Sapphire femtosecond pulsed laser (MaiTai eHP DeepSea, Spectra Physics) was used as excitation source, operating at 80 MHz, and tuned to a wavelength of 880 nm with SHG intensity recorded on Leica NDD 4Tune HyD detectors (424-456 nm). Individual z-stacks were then exported and SHG signal intensity was quantified using Matlab (MathWorks, USA).

*FUCCI cell cycle reporter imaging:* Imaging of the FUCCI cell cycle reporter in live tissues was performed on an inverted Leica Stellaris 8 FALCON DIVE multiphoton inverted microscope. A Ti:Sapphire femtosecond pulsed laser (MaiTai eHP DeepSea, Spectra Physics) was used as excitation source, operating at 80 MHz and tuned to a wavelength of 930 nm (SHG, mAzarmi Green) or 1100 nm (mKusabira Orange) with a 25x 0.95 NA water objective used for imaging. The signal was recorded using Leica NDD 4Tune HyD detectors (455-465 nm for SHG signal, 485-540 nm for mAzami Green and 560-650 for mKusabira Orange). Ten regions of interest (ROI, 512 px x 512 px) per tumour were imaged over a 20 µm z-stack with a step size of 1.26 µm. 3D maximum projections were then generated using Leica LASX software and analysed using QuPath to quantify percentage of red, green and yellow nuclei indicative of $G_1/G_0$, $G_2/M$ or $G_1/S$ cell cycle phase (*32*), respectively. Analysis of the FUCCI reporter was performed on 10 regions of interest (ROIs) per tumour.

*FLIM-FRET imaging of the FAK biosensor:* For *in vivo* measurements of ECFP fluorescence lifetime in subcutaneous xenografts, KPC-FAK cancer cells were injected into the flank of BALB/c-Fox1nuAusb mice. 4 h post-treatment with vehicle or narmafotinib (10 mg/kg) tumours were surgically exposed using skin flap surgery. Imaging was performed using an inverted Leica DMI 6000 SP8 confocal microscope with a Titanium-Sapphire femtosecond laser cavity (Coherent Chameleon Ultra II) excitation source tuned to a wavelength of 840 nm for ECFP excitation. Signal was recorded using RLD-HyD detectors (using bandpass emission filters at 435/40 nm for SHG signal, 483/40 nm for FLIM). FLIM data was acquired with a Picoharp 300 TCSPC system (Picoquant) and image stabilisation performed using Galene (*33*). 50 cells per condition across multiple ROIs per tumour *in vivo* were acquired with a scan speed of 400 Hz, with 2 min 30 s acquisition time and a pixel dwell time of 5 µs. Analysis of ECFP lifetimes was performed using FLIMfit (*34*) by manual selection of single cell membrane regions of interest and recording of the exponential function fit to the fluorescence decay data. Mono-exponential decay lifetime references were calculated using a Chroma slide. Lifetime maps were generated from raw data with a smoothing by a 2x2 pixel kernel, and application of a standard rainbow look-up table, with blue indicating low ECFP fluorescence lifetime and green to yellow indicating high ECFP fluorescence lifetime. The background intensity threshold was set to the average background pixel value for each image to exclude areas of unspecific signal and is shown on the lifetime maps as black. Analysis of the FAK biosensor was conducted on 50 cells per mouse.

*IVIS imaging:* Orthotopic tumour growth was monitored *via* luciferase signal imaging on an IVIS Spectrum (PerkinElmer). Luciferin (150 mg/kg, Gold Biotechnology) was administered by intraperitoneal injection, 3 min prior to imaging. Anesthetised (isoflurane 2 L, $O_2$ 1 L/min, with continuous vacuum to remove excess $O_2$ and isoflurane) mice were placed on the IVIS stage, exposing the left flank and the signal was acquired with open filters and small binning. Tumour burden was determined based on total flux.

**<u>Supplementary Table Legends</u>**

**Supplementary Table S1:** Information on the Royal North Shore Hospital/Australian Pancreatic Matrix Atlas (APMA) cohort of treatment-naïve and neoadjuvant gemcitabine/Abraxane- or FOLFIRINOX-treated PDAC patients.

**Supplementary Table S2:** Association of pY397-FAK with clinico-pathological features in the Australian Pancreatic Genome Initiative (APGI) cohort of PDAC patients. P-values were determined using Chi-square test. Significant correlations are highlighted in red.

**Supplementary Table S3.** S scores for narmafotinib screen at 1 μM.

**Supplementary Table S4:** Pharmacokinetic parameters for narmafotinib in male Swiss mice following IV (5 mg/kg) and oral (20 mg/kg) administration.

**Supplementary Table S5:** Differentially expressed transcripts determined by RNA-seq of syngeneic KPC cancer cells *versus* syngeneic KPC-educated CAFs (n=3 biological replicates per cell line). Adjusted p-value $< 0.05$, fold change $> 1.25$.

**Supplementary Table S6:** Complete mutational status of patient-derived cell lines (PDCLs) TKCC2.1lo and TKCC10lo from the Australian Pancreatic Genome Initiative (APGI) cohort of PDAC patients (*3*).

**Supplementary Table S7:** Differentially expressed transcripts determined by RNA-seq on TKCC10lo PDCL subcutaneous tumours primed with vehicle prior to gemcitabine/Abraxane or primed with narmafotinib prior to gemcitabine/Abraxane (n=5 animals per treatment group). Adjusted p-value < 0.05, fold change > 1.25.

### Supplementary Figures

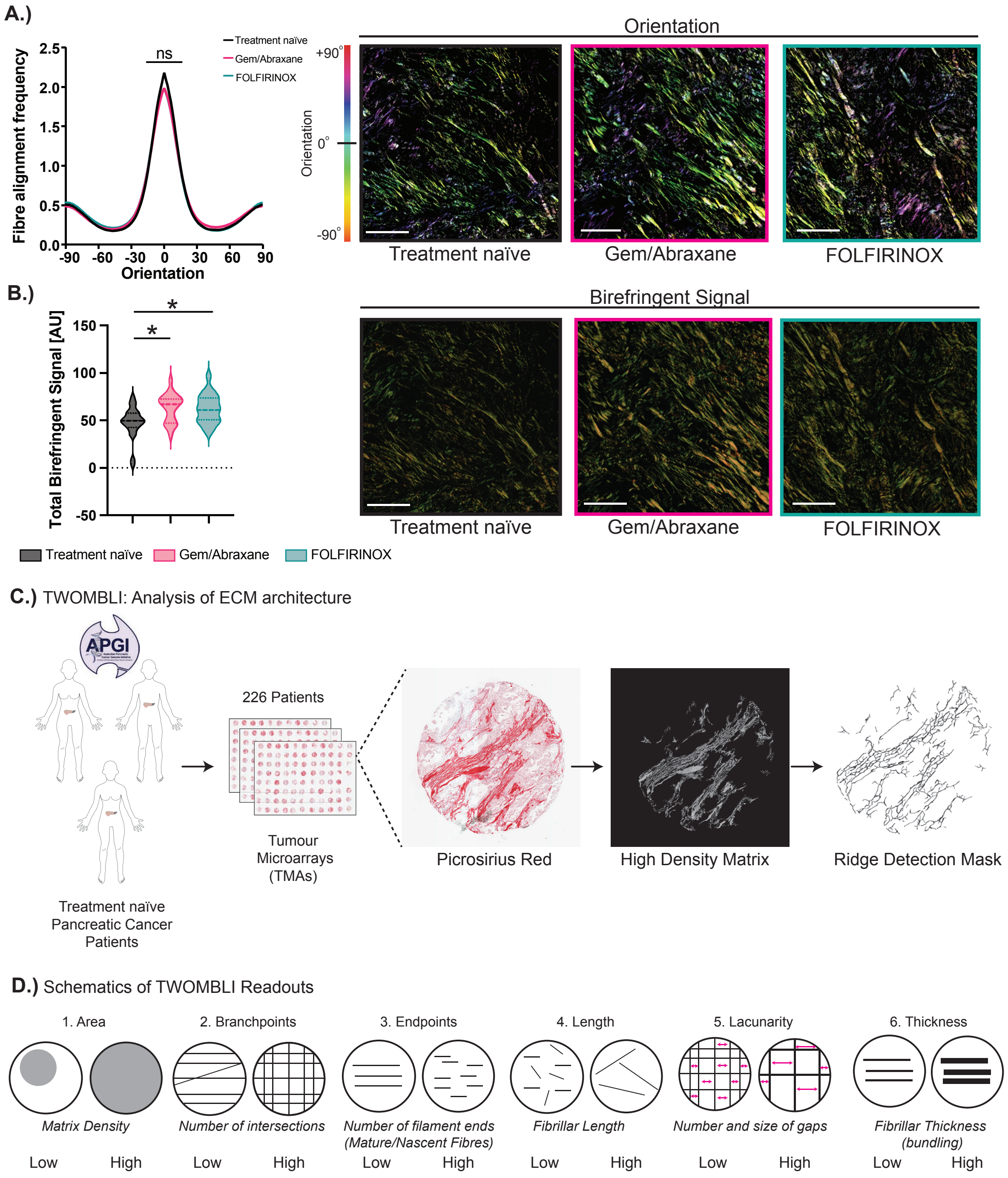


**Supplementary Figure 1**

**Supplementary Figure S1. Analysis of collagen fibre orientation and birefringence signal in PDAC patients with schematics of TWOMBLI analysis of ECM organisation and resulting fibre and matrix features.** (**A,B**) Quantification and representative images of fibre orientation/alignment analysis (**A.**) as well as birefringence signal coverage and representative birefringence images acquired by polarised light imaging of the Picrosirius Red signal (**B.**) in tumours from treatment-naïve versus neoadjuvant gemcitabine/Abraxane- or FOLFIRINOX-treated PDAC patients in the APMA cohort. Scale bar, 100 μm. n=22 treatment-naïve, n=20 neoadjuvant gemcitabine/Abraxane, n=19 neoadjuvant FOLFIRINOX patients, with ≥5 ROIs per tumour. The local orientation of fibres in (**A.**) is represented by the corresponding colour assigned to each specific angle of orientation. (**C.**) Schematic of TWOMBLI analysis of ECM architecture: input Picrosirius Red stained TMA core, output high density matrix and processed ridge detection mask. (**D.**) Schematic examples of output matrix metrics. Adapted from Wershof *et al.* (*23*). Violin plots are median (dashed line) with quartiles (dotted line), p-values determined using (**A.**) Kruskal-Wallis test with Dunn's multiple comparisons (at orientation 0°) or (**B.**) ordinary one-way ANOVA with Sídák multiple comparison test. ns, P≥0.05, *P<0.05.

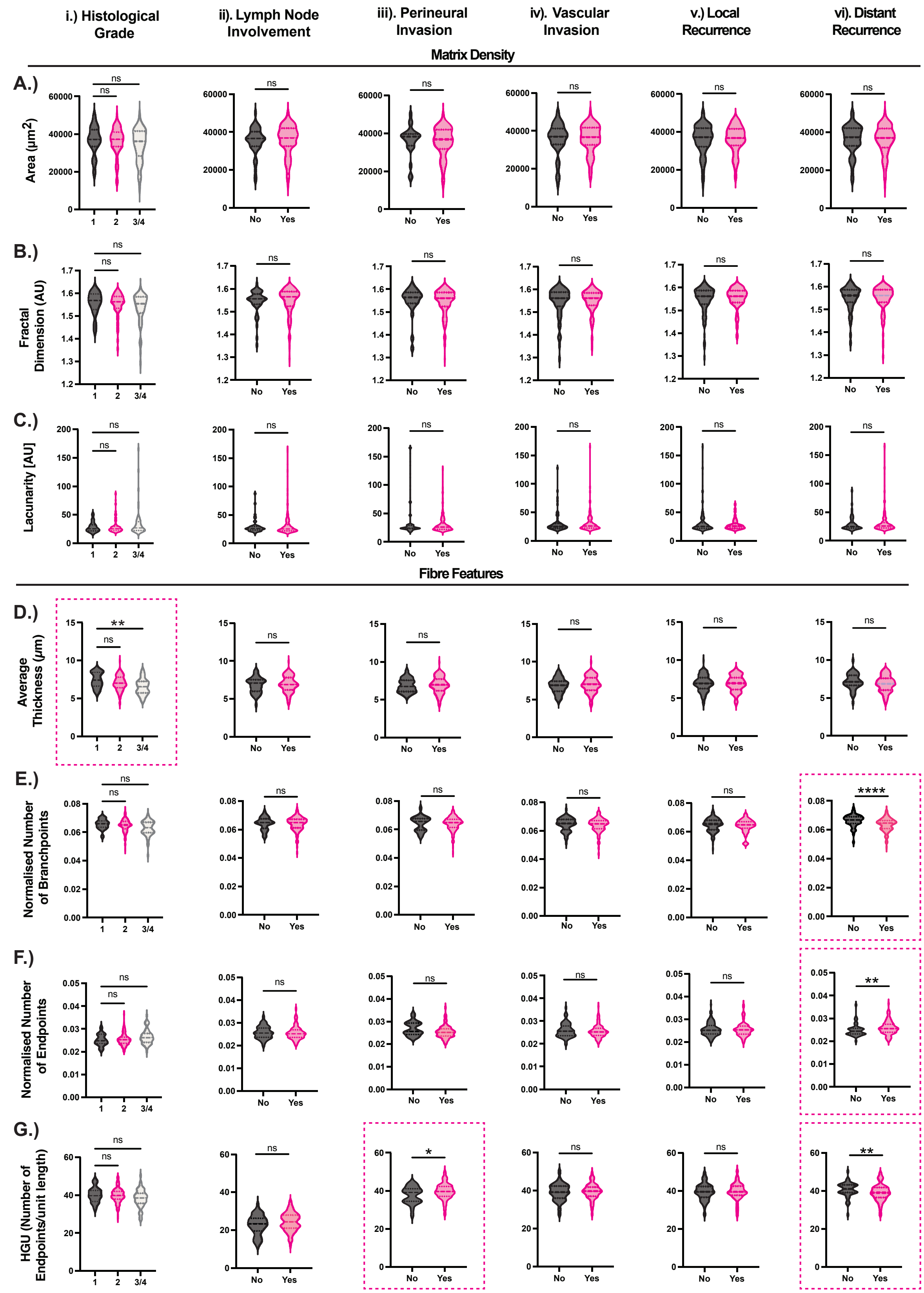


**Supplementary Figure 2**

**Supplementary Figure S2. TWOMBLI analysis of ECM organisation in relation to clinico-pathological features in PDAC patients.** (**A-G**) Analysis of collagen matrix and fibre features in PDAC patient tumours in relation to (**i.)** histological grade, **ii.)** lymph node involvement, **iii.)** perineural invasion, **iv.)** vascular invasion, **v.)** local recurrence, **vi.)** distant recurrence. Matrix features reported are area (**A.**), box counting fractal dimension (**B.**) and lacunarity (**C.**). Fibre features reported are fibre thickness (**D.**), number of branchpoints (**E.**), number of endpoints (**F.**) and Hyphal Growth Unit (HGU, **G.**). (n≥164 patients). Violin plots are median (dashed line) with quartiles (dotted line), p-values determined using one-way ANOVA with Dunnett's multiple comparison test for normally distributed data or Kruskal-Wallis test with Dunn's multiple comparisons test for non-normally distributed data (**i.)**) or unpaired t-test with Welsh correction for normally distributed data or a Mann-Whitney test for non-normally distributed data (**ii.) - vi.)**). ns, P≥0.05, *P<0.05, **P<0.01, ****P<0.0001.

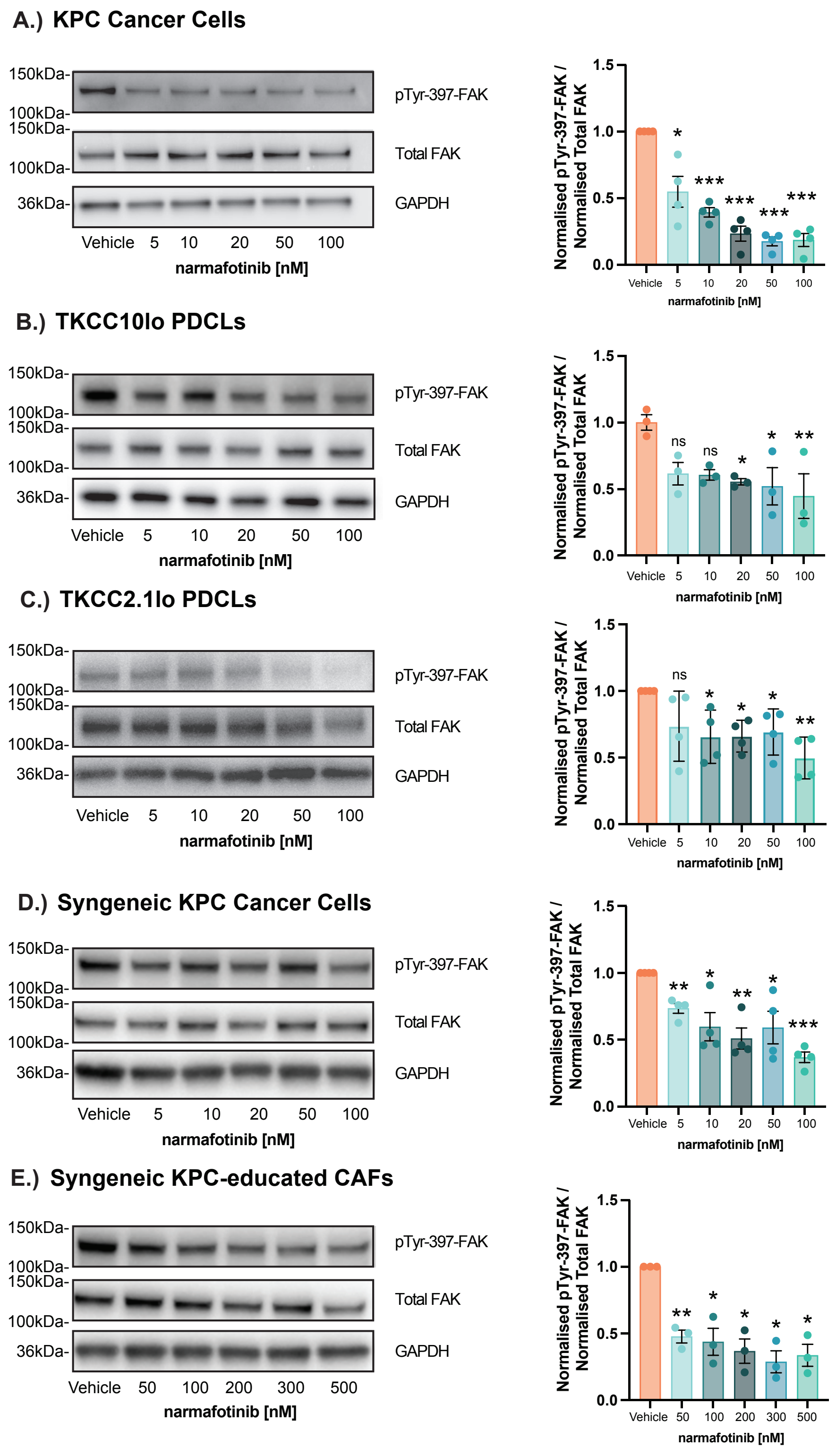


712 **Supplementary Figure 3**

713

**Supplementary Figure S3. Effect of narmafotinib on pTyr-397-FAK phosphorylation.** (**A-E**) Western blot analysis (representative blots for pTyr-397-FAK, total FAK and GAPDH) and quantification of pTyr-397-FAK in cells treated for 2 hours with vehicle or ascending concentrations of narmafotinib using KPC cells (**A.**), TKCC10lo PDCLs (**B.**), TKCC2.1lo PDCLs (**C.**), syngeneic KPC cancer cells (**D.**) and syngeneic KPC-educated CAFs (**E.**). n≥3 repeats. Results are mean ± SEM, p-values were determined using one sample t and Wilcoxon test (**A,C-E**) or ordinary one-way ANOVA with Dunnetts multiple comparison test (**B**). Unless otherwise stated, all significance is compared to vehicle. ns, P≥0.05, *P<0.05, **P<0.01, ***P<0.001.

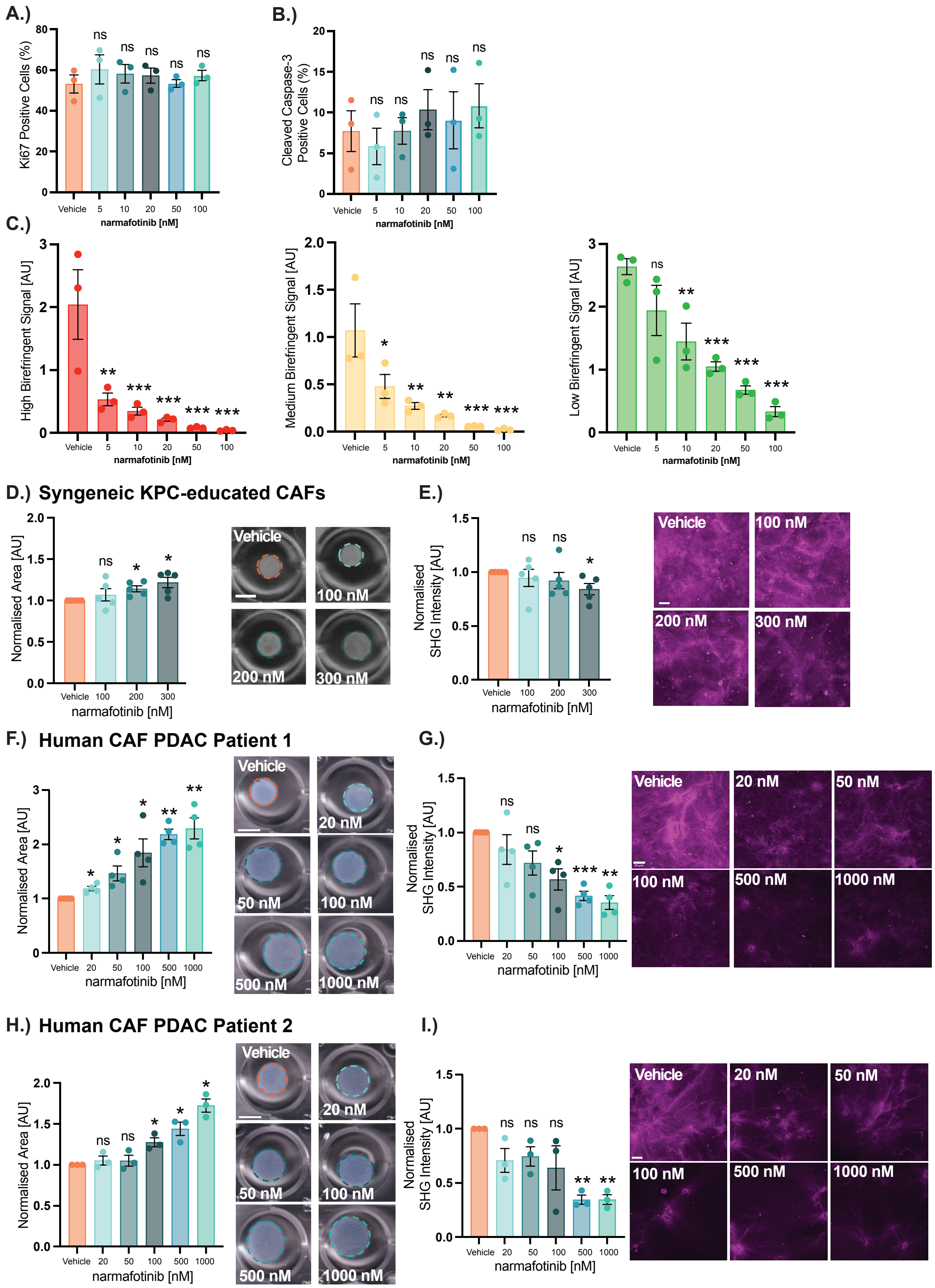


**Supplementary Figure 4**

**Supplementary Figure S4. Narmafotinib reduces collagen remodelling of murine and human CAFs.** (**A,B**) Quantification of Ki67 (**A.**) and cleaved caspase-3 (**B.**) in fibroblasts during contraction of collagen matrices in the presence of vehicle or narmafotinib. (**C.**) Quantification of high (red), intermediate (yellow) and low (green) birefringence signal as a readout of a range from highly dense to poorly dense collagen fibres in Picrosirius Red stained collagen matrices contracting in the presence of vehicle or narmafotinib. n=3 biological repeats, with three matrices per repeat. (**D-I**) Quantification and representative images of matrix contraction ([**D,F,H**], Day 12, scale bar, 1 cm) as well as quantification and representative images of SHG ([**E,G,I**], Day 12, scale bar, 50 μm) for collagen matrices contracted by syngeneic KPC-educated CAFs (**D,E**) as well as human CAFs isolated from PDAC patient 1 (**F,G**) and PDAC patient 2 (**H,I**). n≥3 biological repeats, with three matrices per repeat and 3 FOVs per matrix. Results are mean ± SEM, p-values were determined using (**A-C**) an ordinary one-way ANOVA with Dunnett's multiple comparison test or (**D-I**) a one sample t and Wilcoxon test. Unless otherwise stated, all significance is compared to vehicle. ns, P≥0.05, *P<0.05, **P<0.01 and ***P<0.001.

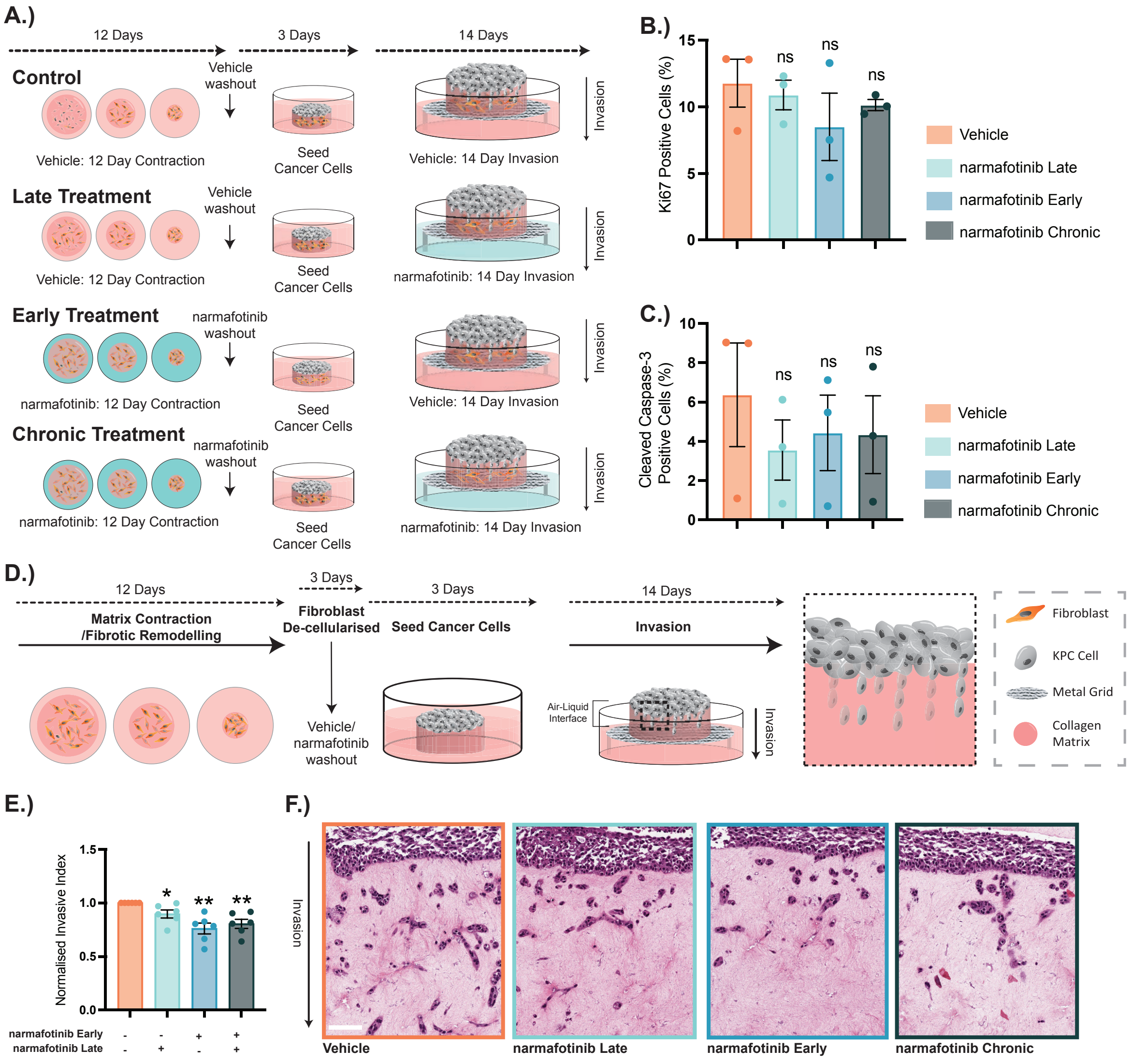


744 **Supplementary Figure 5**

**Supplementary Figure S5. Effect of narmafotinib-mediated FAK inhibition on cell proliferation, survival and invasion in 3D organotypic invasion assays.** (**A.**) Schematic representation of organotypic invasion assay, depicting treatment with vehicle (control), late narmafotinib (during invasion), early narmafotinib (during contraction) and chronic narmafotinib (during contraction and invasion). (**B,C**) Quantification of Ki67 (**B.**) and cleaved caspase-3 (**C.**) in KPC cells invading into organotypic matrices. n=3 biological repeats, with three matrices per repeat and three FOVs per matrix. (**D.**) Schematic representation of organotypic invasion assay with de-cellularised matrices. (**E,F**) Quantification (**E.**) and representative images (**F.**) of invasion into de-cellularised 3D organotypic matrices (fibroblasts removed after contraction phase) upon treatment with vehicle or 20 nM narmafotinib during KPC cell invasion, matrix contraction or matrix contraction and invasion (chronic treatment). n=6 biological repeats, with three matrices per repeat and three FOVs per matrix. Scale bar, 100 μm. Results are mean ± SEM, p-values were determined using (**B, C**) Kruskal-Wallis test with Dunn's multiple comparison or (**E**) one sample t test. Unless otherwise stated, all significance is compared to vehicle. ns, $P \geq 0.05$, $*P<0.05$ and $**P<0.01$.

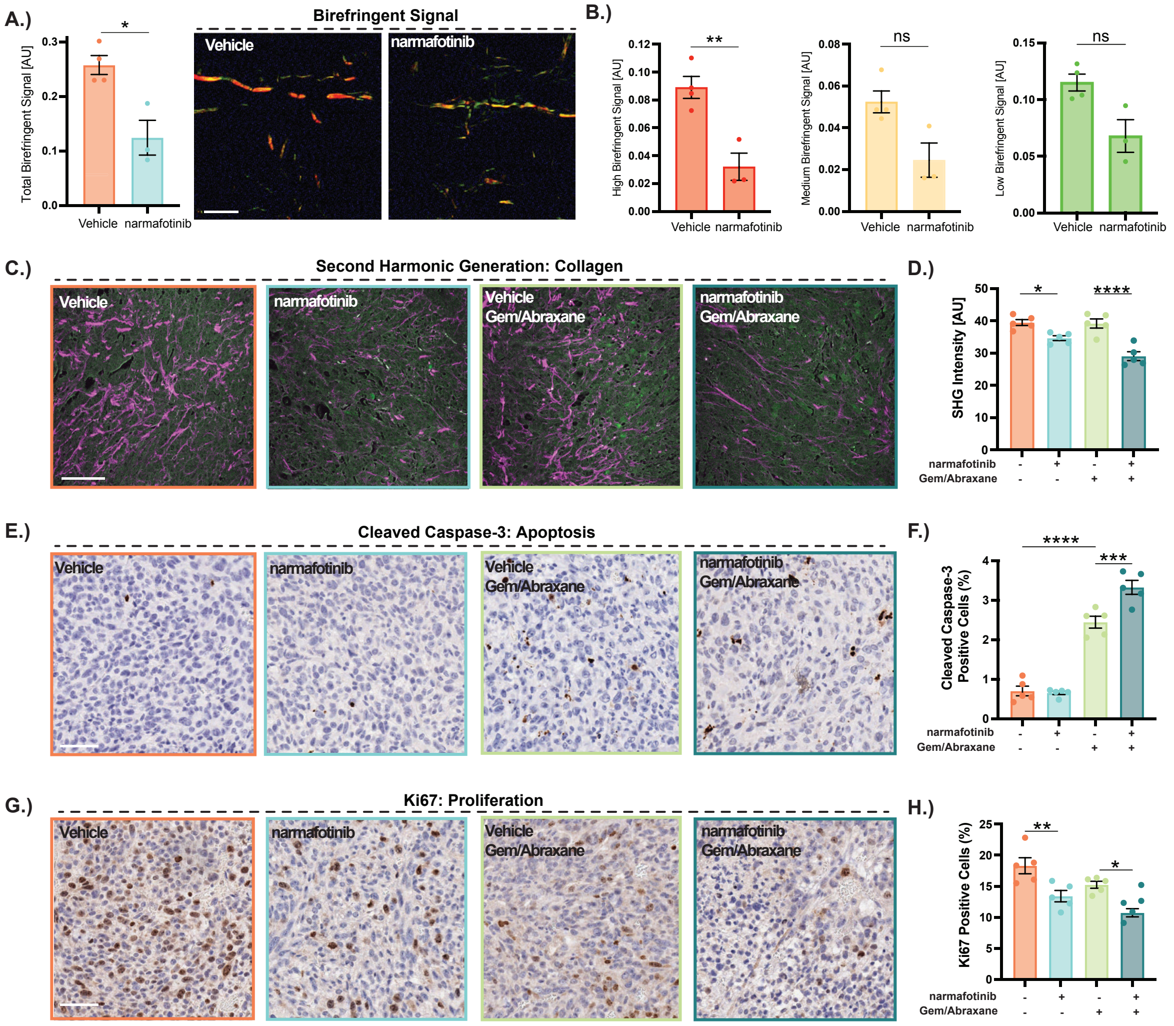


762 **Supplementary Figure 6**

**Supplementary Figure S6. Narmafotinib priming reduces collagen remodelling and deposition, cell survival and cell proliferation in subcutaneous KPC tumours.** (**A.**) Quantification of total birefringence signal and representative polarised light images of tumours stained with Picrosirius Red upon treatment with vehicle or narmafotinib from KPC-FAK-FRET intravital imaging study (scale bar, 100 μm). **(B.)** Quantification of high (red), intermediate (yellow) and low (green) birefringence signal as a readout of a range from highly mature to immature collagen fibres in Picrosirius Red stained tumours treated with vehicle or narmafotinib from KPC-FAK-FRET intravital imaging study. Vehicle n=4 mice, narmafotinib n=3 mice with 8 ROIs per tumour. (**C-H**) Representative images and quantification of SHG (**C,D**), cleaved caspase-3 (**E,F**) and Ki67 (**G,H**) in tumours treated with vehicle or narmafotinib prior to saline or gemcitabine/Abraxane from KPC-FUCCI study. n=5 FOV per animal, 5 animals per treatment group. Scale bar, 100 μm (**A,C**), 50 μm (**E,G**). Results are mean ± SEM, p-values were determined using (**A,B**) an unpaired two-tailed t-test with Welch correction for unequal variance for normally distributed data or a Mann-Whitney test for non-normally distributed data or (**D,F,H**) a one-way ANOVA with Šidák multiple comparisons test. ns, $P \geq 0.05$, *$P<0.05$, **$P<0.01$, ***$P<0.001$, ****$P<0.0001$.

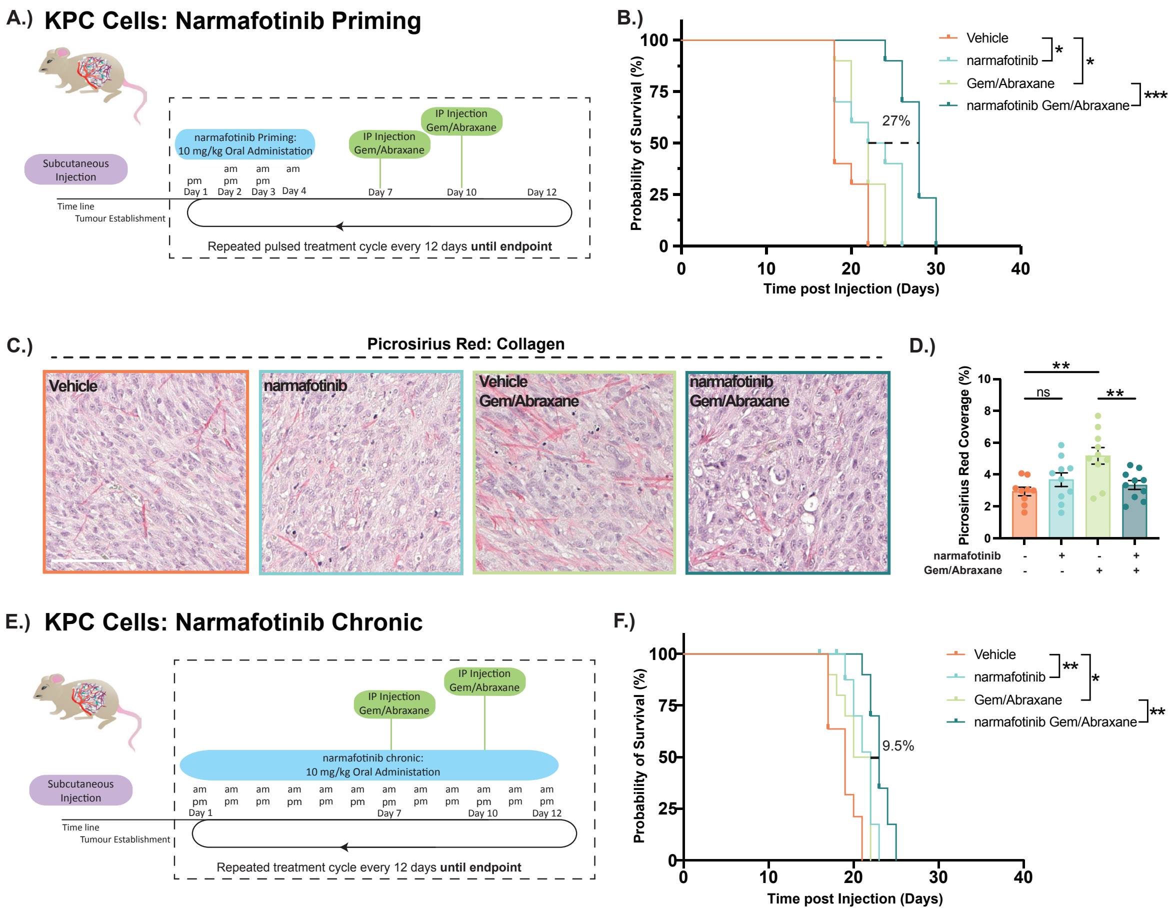


781 **Supplementary Figure 7**

**Supplementary Figure S7. Narmafotinib priming extends survival in a subcutaneous KPC tumour model.** (**A.**) Treatment schedule and timeline for KPC subcutaneous survival study using narmafotinib priming. (**B.**) Kaplan-Meier analysis of survival in mice with KPC subcutaneous tumours treated with vehicle (orange), narmafotinib priming (blue), gemcitabine/Abraxane alone (light green) or narmafotinib priming prior to gemcitabine/Abraxane (dark green). n=10 mice per group. Groups were compared using a log-rank Mantel-Cox test. (**C,D**) Representative images of Picrosirius Red staining (**C.**) and quantification of Picrosirius Red coverage (**D.**) in tumours treated with vehicle, narmafotinib, gemcitabine/Abraxane alone or narmafotinib priming prior to gemcitabine/Abraxane from KPC subcutaneous survival study. n=6 FOV per animal, n≥9 animals per group. Scale bar, 100 µm. Results are mean ± SEM, p-values were determined using an ordinary one-way ANOVA with Sídák multiple comparisons test. (**E.**) Treatment schedule and timeline for KPC subcutaneous survival study using chronic daily narmafotinib treatment. (**F.**) Kaplan-Meier analysis of survival in mice with KPC subcutaneous tumours treated with chronic vehicle (orange), chronic narmafotinib (blue), gemcitabine/Abraxane alone (light green) or chronic narmafotinib in combination with gemcitabine/Abraxane (dark green). n=10 mice per group. Groups were compared using a log-rank Mantel-Cox test. ns, P≥0.05, *P<0.05, **P<0.01, ***P<0.001.

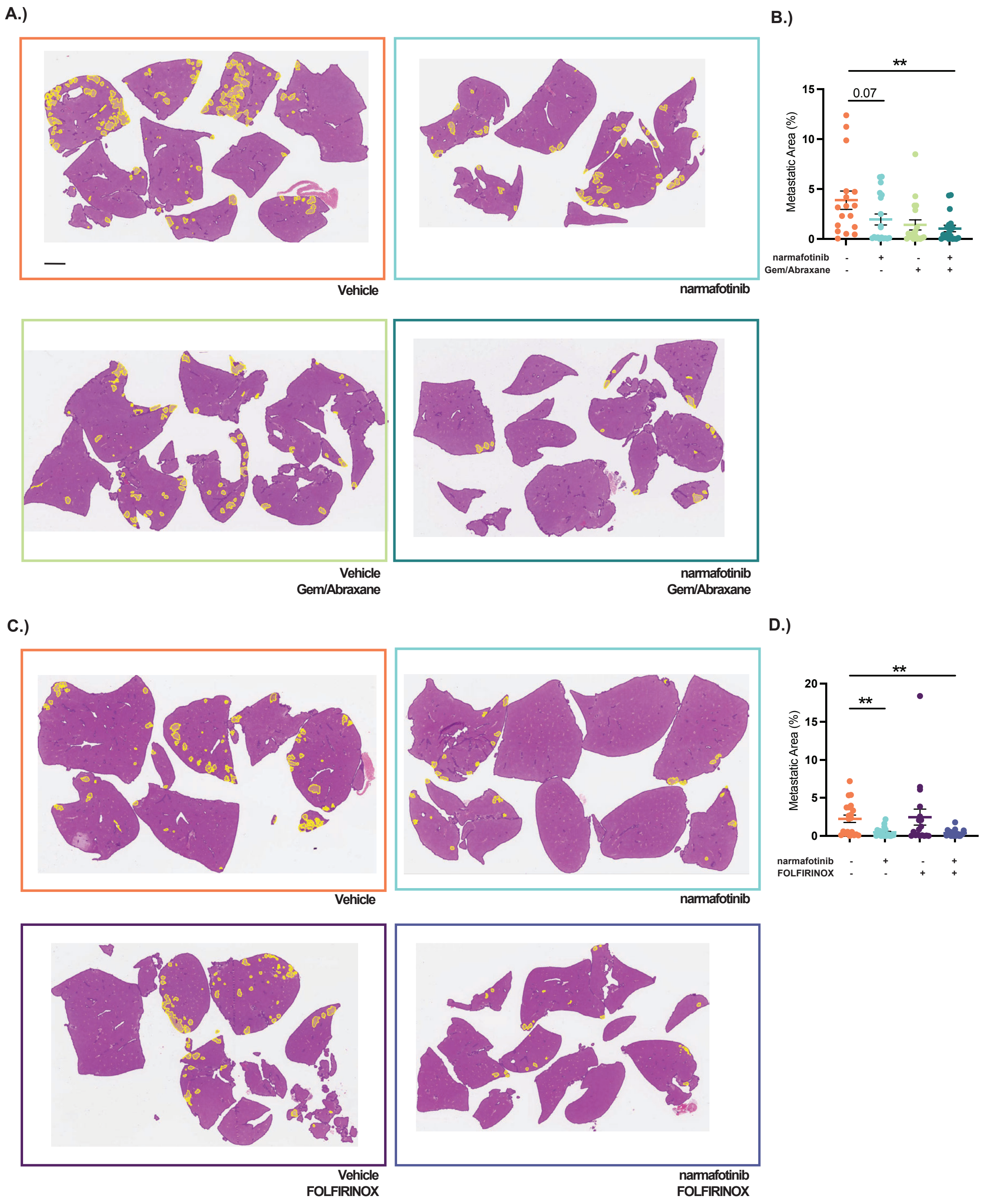


**Supplementary Figure 8**

**Supplementary Figure S8. Narmafotinib priming reduces KPC liver metastasis.** (**A,B**) Representative scanned liver sections stained with H&E (**A.**) and quantification of metastatic area in the liver (**B.**) following intrasplenic KPC cell injection into mice treated with vehicle, narmafotinib, gemcitabine/Abraxane alone or narmafotinib prior to gemcitabine/Abraxane. n≥17 mice per treatment group. (**C,D**) Representative scanned liver sections stained with H&E (**C.**) and quantification of metastatic area in the liver (**D.**) following intrasplenic KPC cell injection into mice treated with vehicle, narmafotinib, FOLFIRINOX alone or narmafotinib prior to FOLFIRINOX. Metastases are outlined in yellow (**A,C**). n≥17 mice per treatment group. Scale bar, 2 mm. Results are mean ± SEM, p-values determined using a Kruskal-Wallis test with Dunn's multiple comparisons. *P<0.05, **P<0.01.

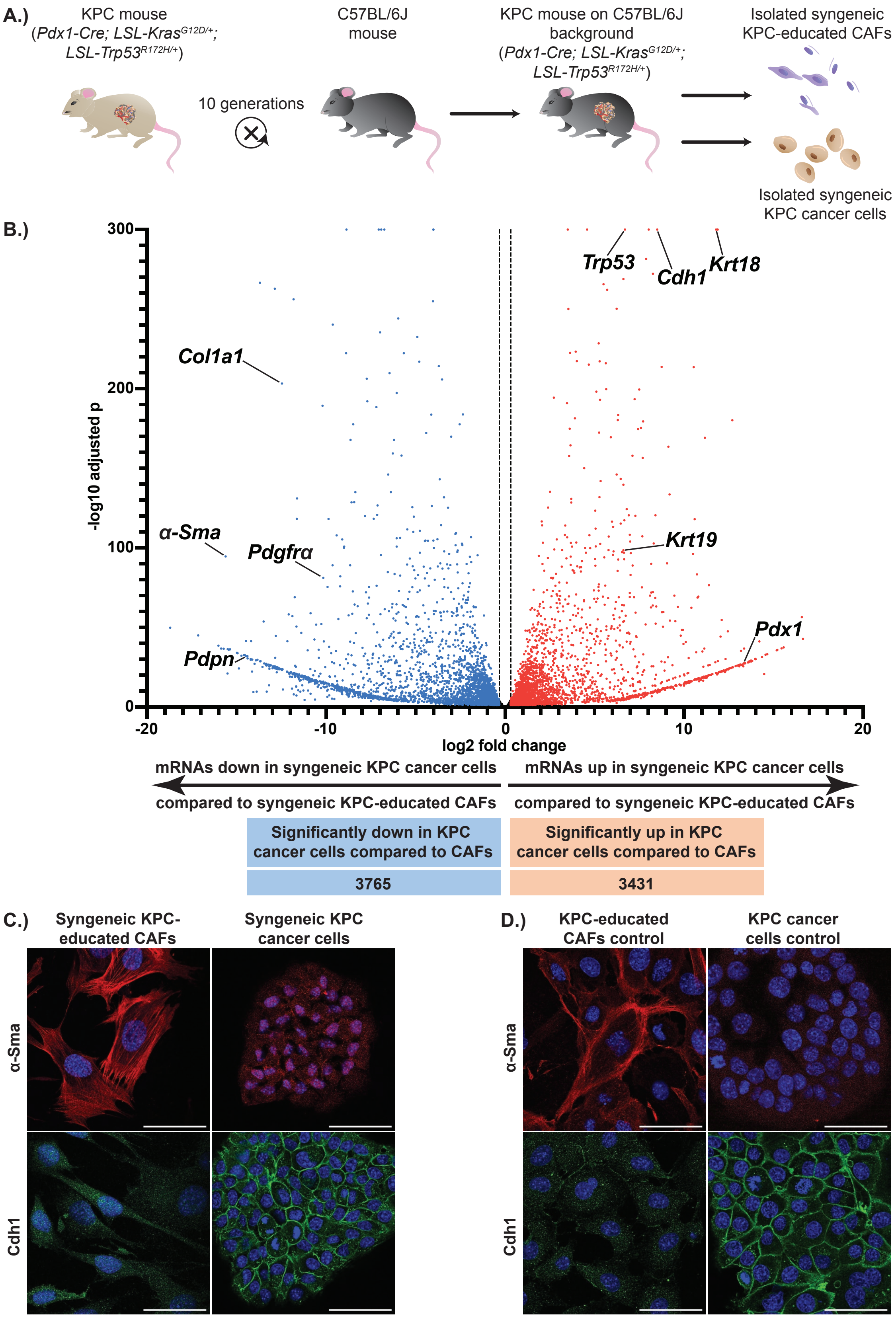


816 **Supplementary Figure 9**

817

**Supplementary Figure S9. Isolation and characterisation of syngeneic KPC cancer cells and syngeneic KPC-educated CAFs.** (**A.**) Schematics depicting the backcrossing of the genetically engineered KPC mouse model (*Pdx1-Cre; LSL-Kras$^{G12D/+}$; LSL-Trp53$^{R172H/+}$*) of PDAC for 10 generations to the C57BL/6J background followed by isolation of syngeneic KPC cancer cells and syngeneic KPC-educated CAFs from end-stage tumours of KPC mice on the C57BL/6J background. (**B.**) Volcano plot showing significantly down-regulated (blue) and significantly up-regulated transcripts (red) in syngeneic KPC cancer cells compared to syngeneic KPC-educated CAFs (n=3 biological replicates per cell line, adjusted $p<0.05$, >1.25 fold change [vertical dashed lines]). Genes with a -log10 adjusted p > 300 were plotted at -log10 adjusted p = 300. (**C,D**) Representative immunofluorescence images of syngeneic KPC-educated CAFs and syngeneic KPC cancer cells (**C.**) as well as KPC-educated control CAFs and KPC control cancer cells (**D.**) stained with anti-α-Sma (top panel, red) or anti-Cdh1 (bottom panel, green) and nuclear counterstain with DAPI (blue). Scale bar, 50 μm.

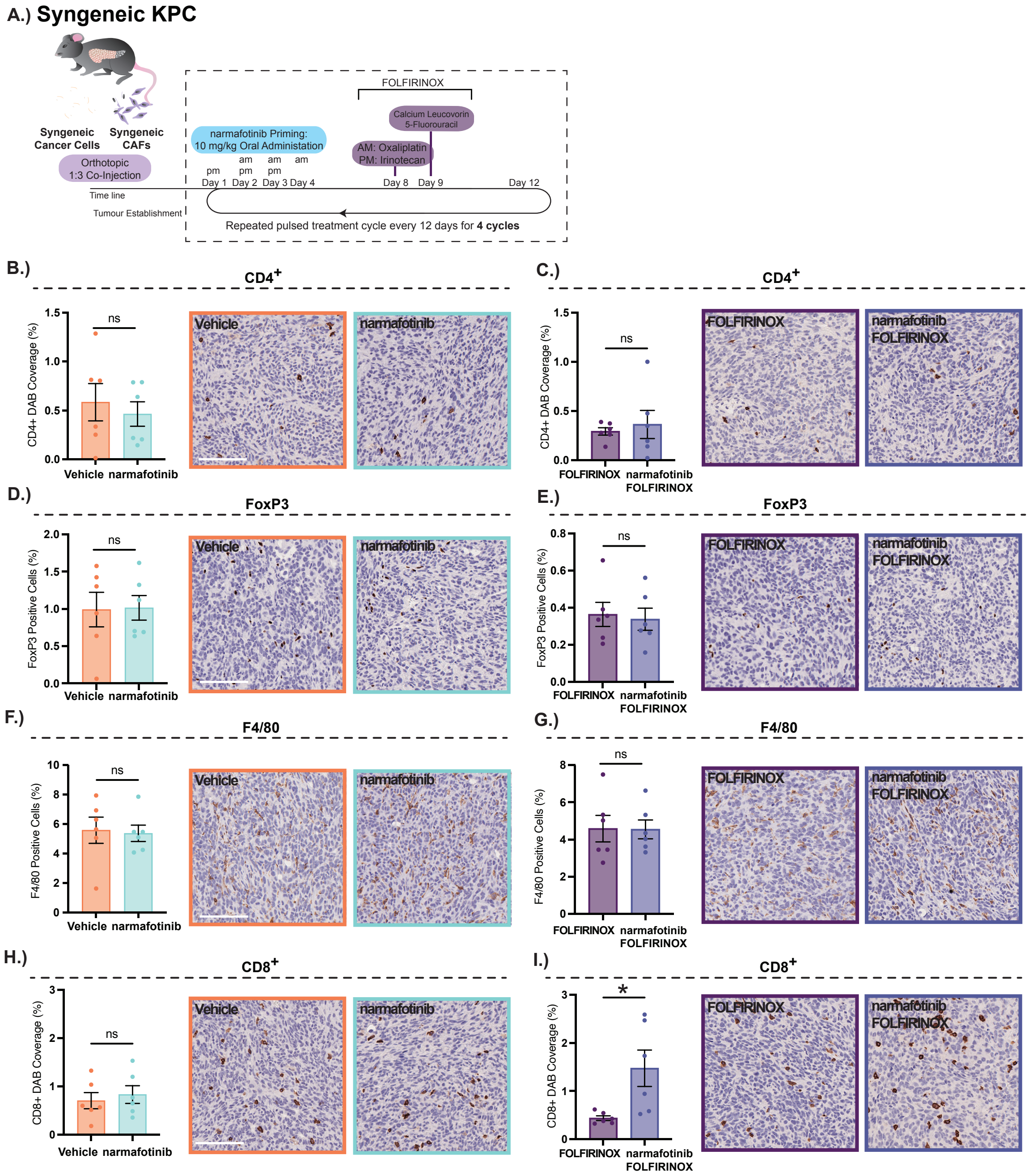


833 **Supplementary Figure 10**

834

**Supplementary Figure S10. Narmafotinib priming increases $CD8^+$ T cell abundance in a syngeneic orthotopic KPC mouse model.** (**A.**) Treatment schedule and timeline for the syngeneic orthotopic KPC model. (**B-I**) Quantification and representative immunohistochemistry images for CD4 (**B,C**), FoxP3 (**D,E**), F4/80 (**F,G**) and CD8 (**H,I**) of primary pancreatic tumours from a syngeneic orthotopic KPC mouse model treated for 4 cycles with vehicle, narmafotinib, FOLFIRINOX alone or narmafotinib prior to FOLFIRINOX. n=6 FOV per animal, 6 animals per group. Scale bar, 100 µm. Results are mean ± SEM, p-values determined using an unpaired t-test with Welch's correction. ns,$P \geq 0.05$, *$P<0.05$.

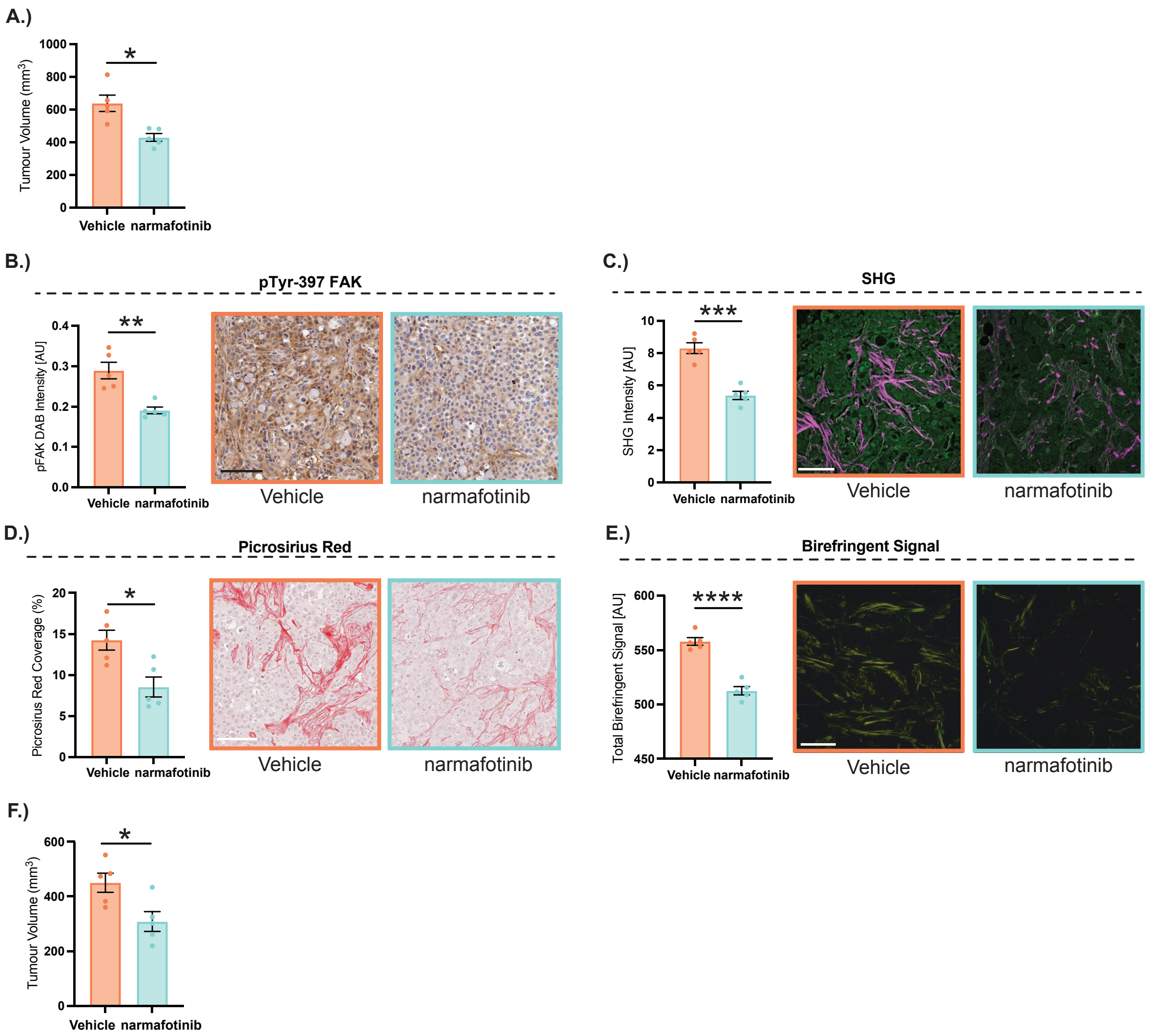


845 **Supplementary Figure 11**

**Supplementary Figure S11. Narmafotinib priming reduces tumour growth and ECM remodelling in subcutaneous TKCC10lo PDCL tumours.** (**A.**) Quantification of subcutaneous TKCC10lo tumour size at timed endpoint following two cycles of treatment with vehicle or narmafotinib. (**B-E**) Quantification and representative images of TKCC10lo PDCL tumours stained with pTyr-397-FAK (**B.**), imaged with SHG (**C.**) and stained with Picrosirius Red and imaged using transmitted light (**D.**) or polarised light (**E.**) following two cycles of treatment with vehicle or narmafotinib. (**F.**) Quantification of subcutaneous TKCC10lo tumour size at timed endpoint following two cycles of treatment with vehicle or narmafotinib. Scale bar, 100 µm. n=5 mice per treatment group, 6 FOV per tumour. Results are mean ± SEM, p-values determined using unpaired two-tailed t-test with Welch's correction for unequal variance. ns, $P \geq 0.05$, *$P<0.05$, **$P<0.01$, ***$P<0.001$, ****$P<0.0001$.

**A.)** Volcano Plot comparing mRNA expression between narmafotinib/GemAbraxane and Vehicle/GemAbraxane

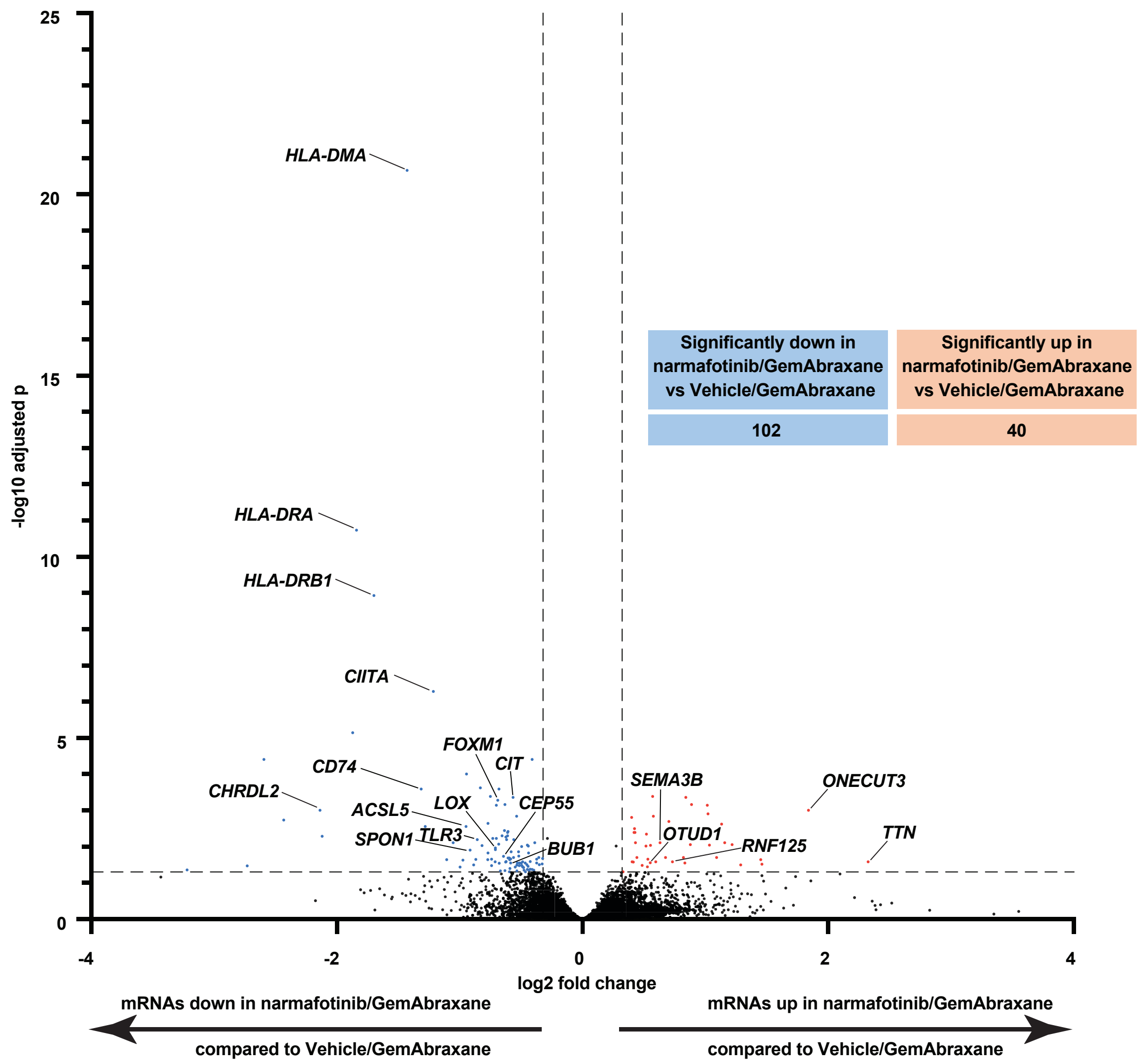


**B.)** RNA-seq 'HALLMARK' GSEA analysis

| pathway | padj | NES |
|---|---|---|
| HALLMARK_E2F_TARGETS | 3.55E-31 | -2.8257437 |
| HALLMARK_G2M_CHECKPOINT | 3.02E-23 | -2.6497053 |
| HALLMARK_INTERFERON_GAMMA_RESPONSE | 6.80E-21 | -2.624788 |
| HALLMARK_MYC_TARGETS_V1 | 4.59E-15 | -2.4095144 |
| HALLMARK_INTERFERON_ALPHA_RESPONSE | 7.43E-10 | -2.3700756 |
| HALLMARK_MYC_TARGETS_V2 | 0.00018479 | -2.0830562 |
| HALLMARK_ALLOGRAFT_REJECTION | 2.30E-05 | -2.0293601 |
| HALLMARK_MITOTIC_SPINDLE | 1.61E-07 | -2.0270361 |
| HALLMARK_SPERMATOGENESIS | 0.00362037 | -1.8317777 |
| HALLMARK_DNA_REPAIR | 0.00153599 | -1.734406 |
| HALLMARK_IL6_JAK_STAT3_SIGNALING | 0.01679046 | -1.6833149 |
| HALLMARK_MTORC1_SIGNALING | 0.00348011 | -1.6381133 |
| HALLMARK_PROTEIN_SECRETION | 0.0120196 | 1.57003802 |
| HALLMARK_MYOGENESIS | 0.00066244 | 1.75099435 |

**C.)** GSEA showing downregulated 'Cancer Hallmarks'

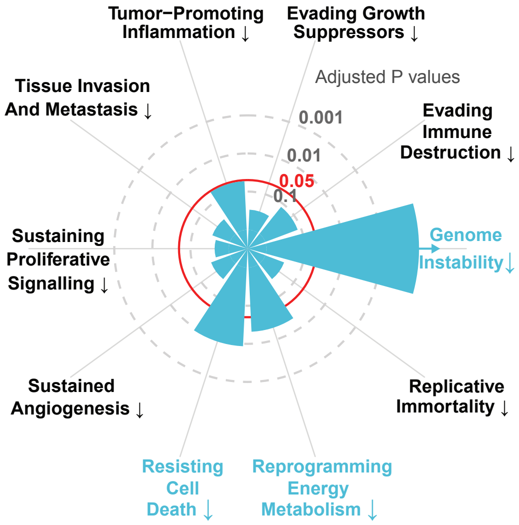


**Supplementary Figure 12**

**Supplementary Figure S12. Gene expression changes in TKCC10lo subcutaneous tumours upon narmafotinib priming prior to gemcitabine/Abraxane compared to vehicle priming prior to gemcitabine/Abraxane. (A.)** Volcano plot showing significantly down-regulated (blue) and significantly up-regulated transcripts (red) in TKCC10lo subcutaneous tumours upon narmafotinib priming prior to gemcitabine/Abraxane compared to vehicle priming prior to gemcitabine/Abraxane following 3 cycles of treatment (n=5 per treatment group, adjusted $p<0.05$ [horizontal line], >1.25 fold change [vertical dashed lines]). **(B.)** Gene Set Enrichment Analysis depicting significantly de-regulated 'HALLMARKS' pathways in TKCC10lo subcutaneous tumours upon narmafotinib priming prior to gemcitabine/Abraxane compared to vehicle priming prior to gemcitabine/Abraxane (n=5 per treatment group, adjusted $p<0.05$). **(C.)** Gene Set Enrichment Analysis depicting significantly de-regulated Cancer Hallmarks pathways (*27*) in TKCC10lo subcutaneous tumours upon narmafotinib priming prior to gemcitabine/Abraxane compared to vehicle priming prior to gemcitabine/Abraxane (n=5 per treatment group). Red circle indicates adjusted p=0.05 and blue segments indicate downregulation of Cancer Hallmarks in TKCC10lo subcutaneous tumours upon narmafotinib priming prior to gemcitabine/Abraxane compared to vehicle priming prior to gemcitabine/Abraxane.

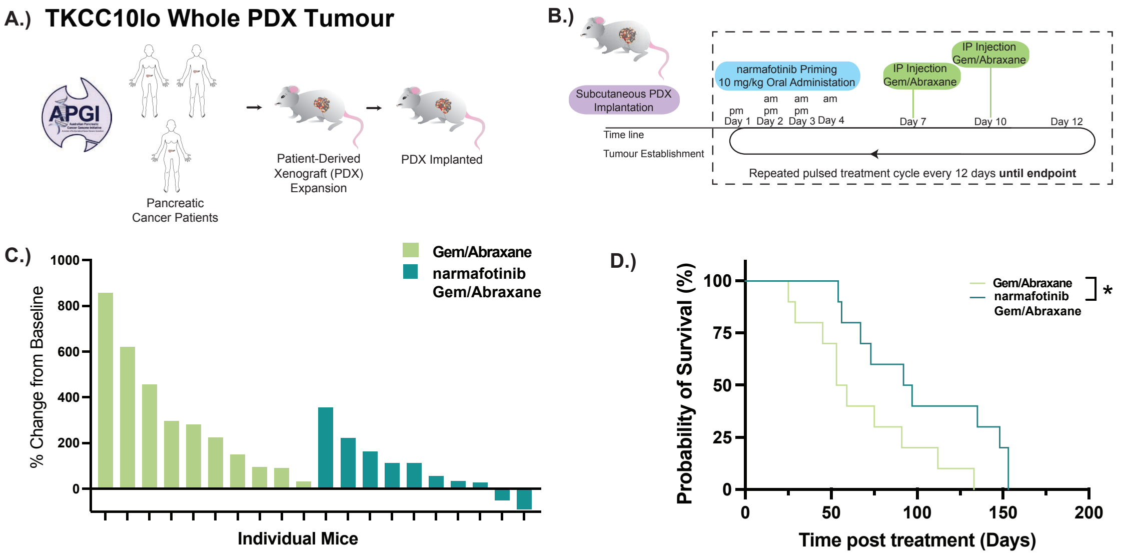


**Supplementary Figure 13**

879

**Supplementary Figure S13. Narmafotinib priming enhances the response to chemotherapy in TKCC10lo PDX tumours.** (**A,B**) Schematics depicting the generation of patient-derived xenografts (PDXs) *via* the Australian Pancreatic Genome Initiative (APGI) (**A.**) as well as treatment schedule and timeline for the TKCC10lo PDX model (**B.**). (**C.**) Waterfall plot of PDX tumour growth as a change from baseline following three cycles of treatment with gemcitabine/Abraxane or narmafotinib priming prior to gemcitabine/Abraxane. (**D.**) Kaplan-Meier analysis of survival in mice with implanted TKCC10lo PDX tumours treated with gemcitabine/Abraxane alone or narmafotinib priming prior to gemcitabine/Abraxane. n=10 mice per treatment group. p-values were determined using a log-rank Mantel-Cox test (**D.**). *P<0.05.

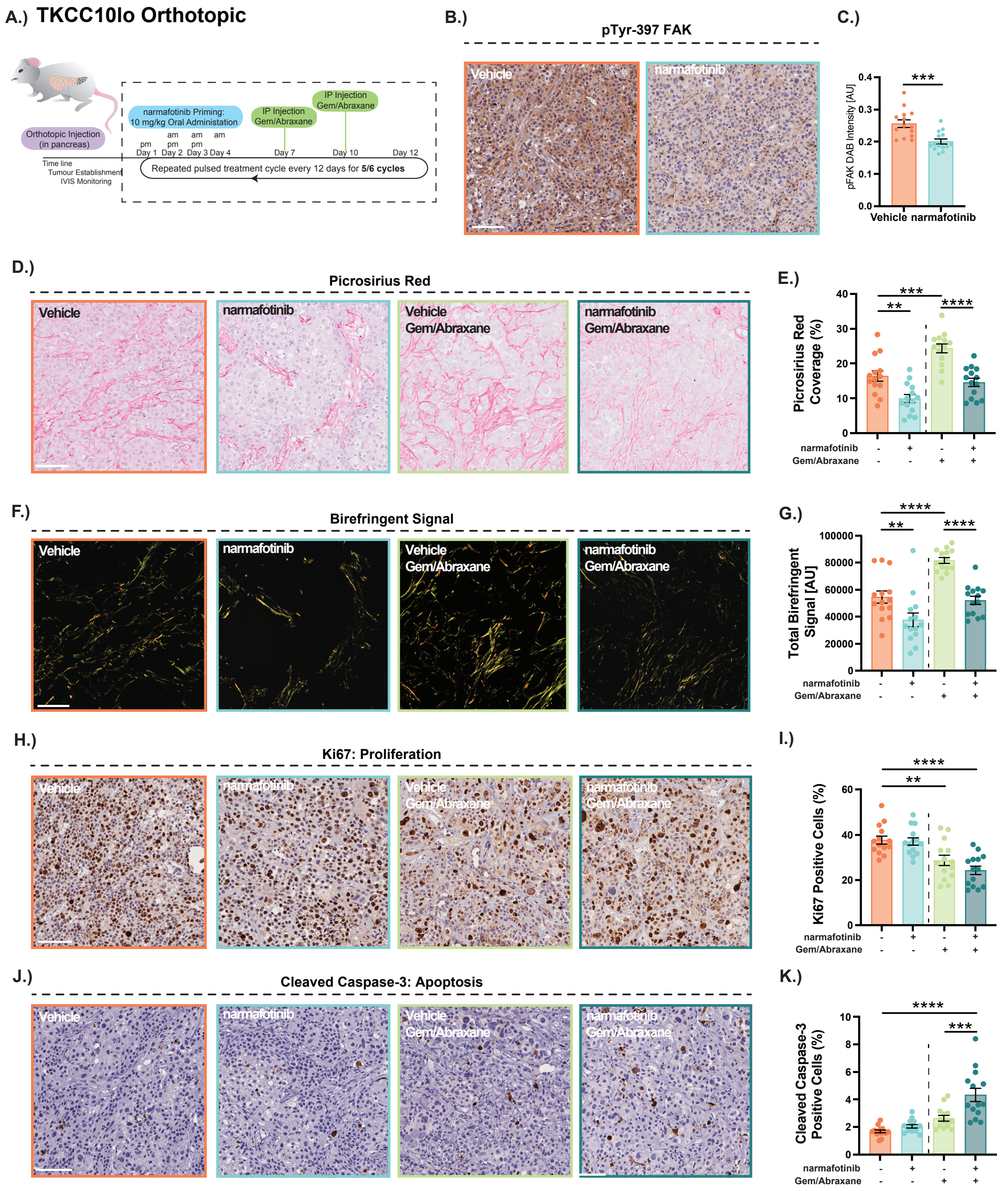


892 **Supplementary Figure 14**

**Supplementary Figure S14. Narmafotinib priming reduces collagen remodelling and deposition and increases apoptosis in orthotopic TKCC10lo PDCL tumours.** (**A.**) Treatment schedule and timeline for the orthotopic TKCC10lo PDCL model to isolate timed endpoint tumours. (**B,C**) Representative images (**B.**) and quantification (**C.**) of pTyr-397 FAK staining in timed endpoint TKCC10lo orthotopic PDCL tumours following 5 cycles of treatment with vehicle or narmafotinib. (**D-G**) Representative images and quantification of Picrosirius Red coverage following brightfield imaging (**D,E**) and birefringence coverage following polarised light imaging of Picrosirius Red (**F,G**) in timed endpoint TKCC10lo orthotopic PDCL tumours following 5 cycles of treatment with vehicle or narmafotinib or following 6 cycles of treatment with gemcitabine/Abraxane or narmafotinib priming prior to gemcitabine/Abraxane. (**H-K**) Representative immunohistochemistry images and quantification of Ki67 (**H,I**) and cleaved caspase-3 (**J,K**) in timed endpoint TKCC10lo orthotopic PDCL tumours following 5 cycles of treatment with vehicle or narmafotinib or following 6 cycles of treatment with gemcitabine/Abraxane or narmafotinib priming prior to gemcitabine/Abraxane. n=6 FOV per animal, n=14 animals per treatment group. Scale bar, 100 µm. Results are mean ± SEM, p-values were determined using (**C.**) an unpaired t-test with Welch's correction or (**E,G,I,K**) an ordinary one-way ANOVA with Sídák multiple comparison. ns, $P \geq 0.05$, $^{*}P<0.05$, $^{**}P<0.01$, $^{****}P<0.0001$.

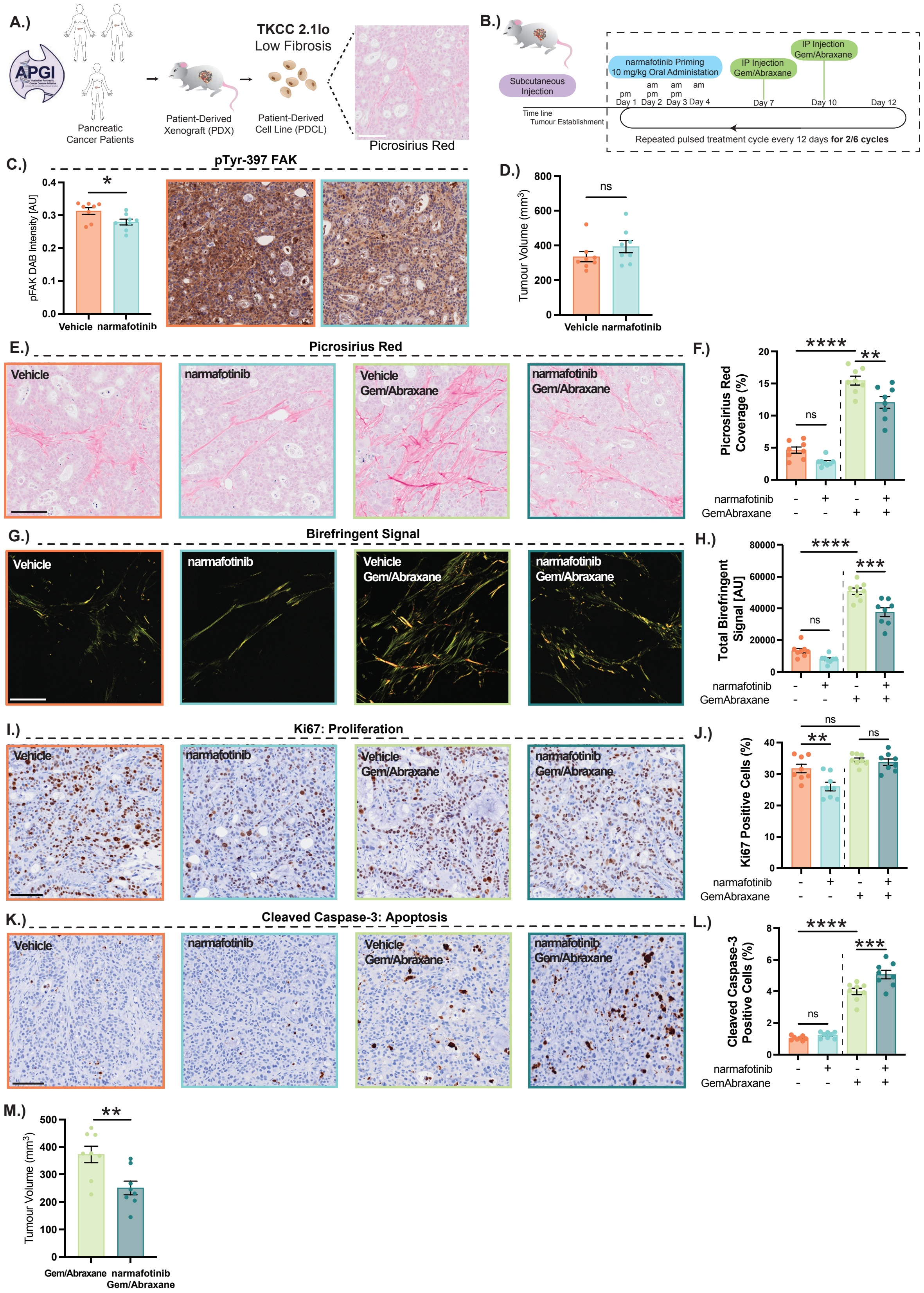


913 **Supplementary Figure 15**

**Supplementary Figure S15. Narmafotinib priming reduces tumour growth, collagen remodeling and deposition and increases apoptosis in subcutaneous TKCC2.1lo PDCL tumours.** (**A.**) Patient-Derived Xenografts (PDXs) and Patient-Derived Cell Lines (PDCLs): TKCC2.1lo PDCL-based tumour, with low collagen levels (Picrosirius Red). (**B.**) Treatment schedule and timeline for TKCC2.1lo model. (**C.**) Quantification and representative images of FAK activity in tumours treated with vehicle or narmafotinib, collected 4 hours post treatment. (**D.**) Quantification of timed endpoint TKCC2.1lo subcutaneous tumour volumes following 2 cycles of treatment with vehicle or narmafotinib. (**E-H**) Representative images and quantification of Picrosirius Red coverage upon brightfield imaging (**E,F**) and birefringence coverage upon polarised light imaging of Picrosirius Red (**G,H**) in timed endpoint TKCC2.1lo subcutaneous tumours following 2 cycles of treatment with vehicle or narmafotinib or following 6 cycles of treatment with gemcitabine/Abraxane or narmafotinib priming prior to gemcitabine/Abraxane. (**I-L**) Representative immunohistochemistry images and quantification of Ki67 (**I,J**) and cleaved caspase-3 (**K,L**) in timed endpoint TKCC2.1lo subcutaneous tumours following 2 cycles of treatment with vehicle or narmafotinib or following 6 cycles of treatment with gemcitabine/Abraxane or narmafotinib priming prior to gemcitabine/Abraxane. (**M.**) Quantification of timed endpoint TKCC2.1lo tumour volumes following 6 cycles of treatment with gemcitabine/Abraxane or narmafotinib priming prior to gemcitabine/Abraxane. n=6 FOV per animal, n=8 animals per treatment group. Scale bar, 100 μm. Results are mean ± SEM. p-values were determined using (**C,D,M**) an unpaired t-test with Welch's correction or (**F,H,J,L**) an ordinary one-way ANOVA with Sídák multiple comparisons. ns, $P \geq 0.05$, $^{*}P<0.05$, $^{**}P<0.01$.

#### Supplementary Movie Legends

**Supplementary Movie S1.** Representative z-stacks of SHG images of collagen deposition and organisation in PDAC patients from the APMA cohort who were treatment naïve (left) or received neoadjuvant gemcitabine/Abraxane (centre) or neoadjuvant FOLFIRINOX (right, scale bar 100 μm).

**Supplementary Movie S2.** Representative z-stack of SHG images of organotypic matrices treated with vehicle or narmafotinib, where narmafotinib reduces fibrillar collagen (scale bar, 50 μm).

**Supplementary Movie S3.** 3D reconstructions of FUCCI cell cycle reporter images in live KPC tumours, where narmafotinib priming prior to gemcitabine/Abraxane chemotherapy increases the proportion of cells in $G_2$/M (scale bar, 100 μm).

**Supplementary Movie S4.** Representative z-stacks of SHG images of KPC tumours, where narmafotinib priming reduces fibrillar collagen content in KPC tumours visualised by SHG imaging (scale bar, 100 μm).

#### <u>Consortium Members of the APGI</u>

**Garvan Institute of Medical Research** Amber L. Johns[1], Anthony J Gill[1,5], Lorraine A. Chantrill[1,22], Paul Timpson[1,] Angela Chou[1,5], Marina Pajic[1], Tanya Dwarte[1], David Herrmann[1], Claire Vennin[1], Thomas R Cox[1], Brooke Pereira[1], Shona Ritchie[1], Daniel A Reed[1], Cecilia R Chambers[1], Xanthe Metcalf[1], Max Nobis[1], Gloria Jeong[1], Ruth J. Lyons[1]. **QIMR Berghofer Medical Research Institute** Nicola Waddell[2], John V. Pearson[2], Ann-Marie Patch[2], Katia Nones[2], Felicity Newell[2], Pamela Mukhopadhyay[2], Venkateswar Addala[2], Stephen Kazakoff[2], Oliver Holmes[2], Conrad Leonard[2], Scott Wood[2]. **University of Melbourne, Centre for Cancer Research** Sean M. Grimmond[3], Oliver Hofmann[3]. **Royal North Shore Hospital** Jaswinder S. Samra[5], Nick Pavlakis[5,] Jennifer Arena[5], Hilda A. High[5]. **Bankstown Hospital** Ray Asghari[6], Neil D. Merrett[6], Amitabha Das[6]. **Liverpool Hospital** Peter H. Cosman[7], Kasim Ismail[7]. **St Vincent's Hospital** Alina Stoita[8], David Williams[8], Allan Spigellman[8]**. Westmead Hospital** Duncan McLeod[9], Judy Kirk[9]. **Royal Prince Alfred Hospital, Chris O'Brien Lifehouse** James G. Kench[10], Peter Grimison[10], Charbel Sandroussi[10], Annabel Goodwin[7,10]. **Prince of Wales Hospital** R. Scott Mead[1,11], Katherine Tucker[11], Lesley Andrews[11]. **Fiona Stanley Hospital** Michael Texler[12], Cindy Forrest[12], Mo Ballal[12,13], David Fletcher[12]**. St John of God Healthcare** Maria Beilin[13], Kynan Feeney[13] Krishna Epari[13] Sanjay Mukhedkar[13]. **Epworth HealthCare** Nikolajs Zeps[23]. **Royal Adelaide Hospital** Nan Q Nguyen[14], Andrew R. Ruszkiewicz[14], Chris Worthley[14]. **Flinders Medical Centre** John Chen[15], Mark E. Brooke-Smith[15], Virginia Papangelis[15]. **Envoi Pathology** Andrew D. Clouston[16]. **Princess Alexandra Hospital** Andrew P. Barbour[17], Thomas J. O'Rourke[17], Jonathan W. Fawcett[17], Kellee Slater[17], Michael Hatzifotis[17], Peter Hodgkinson[17]. **Austin Hospital** Mehrdad Nikfarjam[18]. **Johns Hopkins Medical Institutes** James R. Eshleman[19], Ralph H. Hruban[19], Christopher L. Wolfgang[19]. **ARC-Net Centre for Applied Research on Cancer** Aldo Scarpa[20], Rita T.

Lawlor[20], Vincenzo Corbo[20], Claudio Bassi[20]. **University of Glasgow** Andrew V Biankin [21], Nigel B. Jamieson[21] David K. Chang[1, 21,] Stephan B. Dreyer[21].

[1]The Kinghorn Cancer Centre, Garvan Institute of Medical Research, 370 Victoria Street, Darlinghurst, Sydney, New South Wales 2010, Australia.

[2]QIMR Berghofer Medical Research Institute, 300 Herston Rd, Herston, Queensland 4006, Australia.

[3]University of Melbourne, Centre for Cancer Research, Victorian Comprehensive Cancer Centre, 305 Grattan Street, Melbourne, Victoria 3000, Australia.

[4] Institute for Molecular Bioscience, University of QLD, St Lucia, Queensland 4072, Australia.

[5]Royal North Shore Hospital, Westbourne Street, St Leonards, New South Wales 2065, Australia.

[6]Bankstown Hospital, Eldridge Road, Bankstown, New South Wales 2200, Australia.

[7]Liverpool Hospital, Elizabeth Street, Liverpool, New South Wales 2170, Australia.

[8] St Vincent's Hospital, 390 Victoria Street, Darlinghurst, New South Wales, 2010 Australia.

[9]Westmead Hospital, Hawkesbury and Darcy Roads, Westmead, New South Wales 2145, Australia.

[10]Royal Prince Alfred Hospital, Missenden Road, Camperdown, New South Wales 2050, Australia.

[11]Prince of Wales Hospital, Barker Street, Randwick, New South Wales 2031, Australia.

[12]Fremantle Hospital, Alma Street, Fremantle, Western Australia 6959, Australia.

[13] St John of God Healthcare, 12 Salvado Road, Subiaco, Western Australia 6008, Australia.

[14] Royal Adelaide Hospital, North Terrace, Adelaide, South Australia 5000, Australia.

[15] Flinders Medical Centre, Flinders Drive, Bedford Park, South Australia 5042, Australia.

[16] Envoi Pathology, 1/49 Butterfield Street, Herston, Queensland 4006, Australia.

[17] Princess Alexandra Hospital, 199 Ipswich Rd, Woolloongabba QLD 4102.

[18] Austin Hospital, 145 Studley Road, Heidelberg, Victoria 3084, Australia.

[19] Johns Hopkins Medical Institute, 600 North Wolfe Street, Baltimore, Maryland 21287, USA.

[20] ARC-NET Center for Applied Research on Cancer, University of Verona, Via dell'Artigliere, 19 37129 Verona, Province of Verona, Italy.

[21] Wolfson Wohl Cancer Research Centre, Institute of Cancer Sciences, University of Glasgow, Garscube Estate, Switchback Road, Bearsden, Glasgow, Scotland G61 1BD, United Kingdom.

[22] Wollongong Hospital, Illawarra and Shoalhaven Local Health District, Loftus Street, Wollongong NSW 2500.

[23] Epworth HealthCare, 89 Bridge Rd, Richmond VIC 3121, Australia.

#### Consortium Members of the APMA

Paul Timpson[1], Thomas R. Cox[1], Marina Pajic[1], Anthony J. Gill[1,2], Jaswinder S. Samra[1,2], Brooke A. Pereira[1], David Herrmann[1], Amber L. Johns[1], Gloria Jeong[1], Shona Ritchie[1], Daniel A. Reed[1], Cecilia R. Chambers[1], Janett Stoehr[1], Morghan C. Lucas[1], Joanna N. Skhinas[1], Lea Abdulkhalek[1], Max Nobis[1], Tatjana Schmitz[1], Victoria Lee[1], Xanthe L. Metcalf[1], Sean M Grimmond[3], Kym Pham Stewart[3], Mehreen Arshi[1], Angela M Steinmann[1], Nicola Blackburn[1], Ruth J. Lyons[1]

[1]The Kinghorn Cancer Centre, Garvan Institute of Medical Research, 370 Victoria Street, Darlinghurst, Sydney, New South Wales 2010, Australia.

[2]Royal North Shore Hospital, Westbourne Street, St Leonards, New South Wales 2065, Australia.

[3]University of Melbourne Centre for Cancer Research, Victorian Comprehensive Cancer Centre, 305 Grattan Street, Melbourne, Victoria, 3000, Australia.
